# BoltzMol-1: Towards Reliable Virtual Screening for Fast and Cost-Effective Hit Discovery

**DOI:** 10.64898/2026.07.04.736485

**Authors:** Noah Getz, Geoffrey Smith, Avene Colgan, Vincent Fan, Luca Cavalleri, Francesco Capponi, Jeremy Wohlwend, Anthony Gitter, Joshua Kritzer, Madison Maiorano, Nathan Wlodarchak, Gabriele Corso, Saro Passaro

## Abstract

We present BoltzMol-1, a small-molecule hit discovery pipeline, centered on an optimized version of Boltz-2, explicitly adapted for prospective discovery. Reliable hit discovery that generalizes across target classes (rather than only the well-characterized families that dominate existing ligand data) would broaden the range of biology accessible to small-molecule intervention and reduce reliance on resource-intensive high-throughput screening. Towards this goal, the system prioritizes compounds for rapid experimental validation by coupling model-driven ranking with streamlined procurement from commercial catalogs. To improve developability at the point of selection, we introduce a suite of ADMET models for kinetic solubility (logS), lipophilicity (logD), and Caco-2 permeability. These models act as an early triage layer, systematically filtering out compounds with unfavorable physicochemical and absorption properties prior to synthesis or purchase. Across a panel of ten targets (most with no representation in the underlying affinity training data) we observe strong prospective performance on challenging systems. Functional actives or binders were identified for 6 of 10 targets, despite modest experimental budgets of 28-96 compounds per target. These results include successes on receptors and enzymes traditionally considered difficult for structure- or ligand-based approaches. Collectively, this work establishes a practical framework for low-throughput, cost-constrained discovery campaigns capable of delivering chemically tractable binders with favorable property profiles.

**Figure 1:**
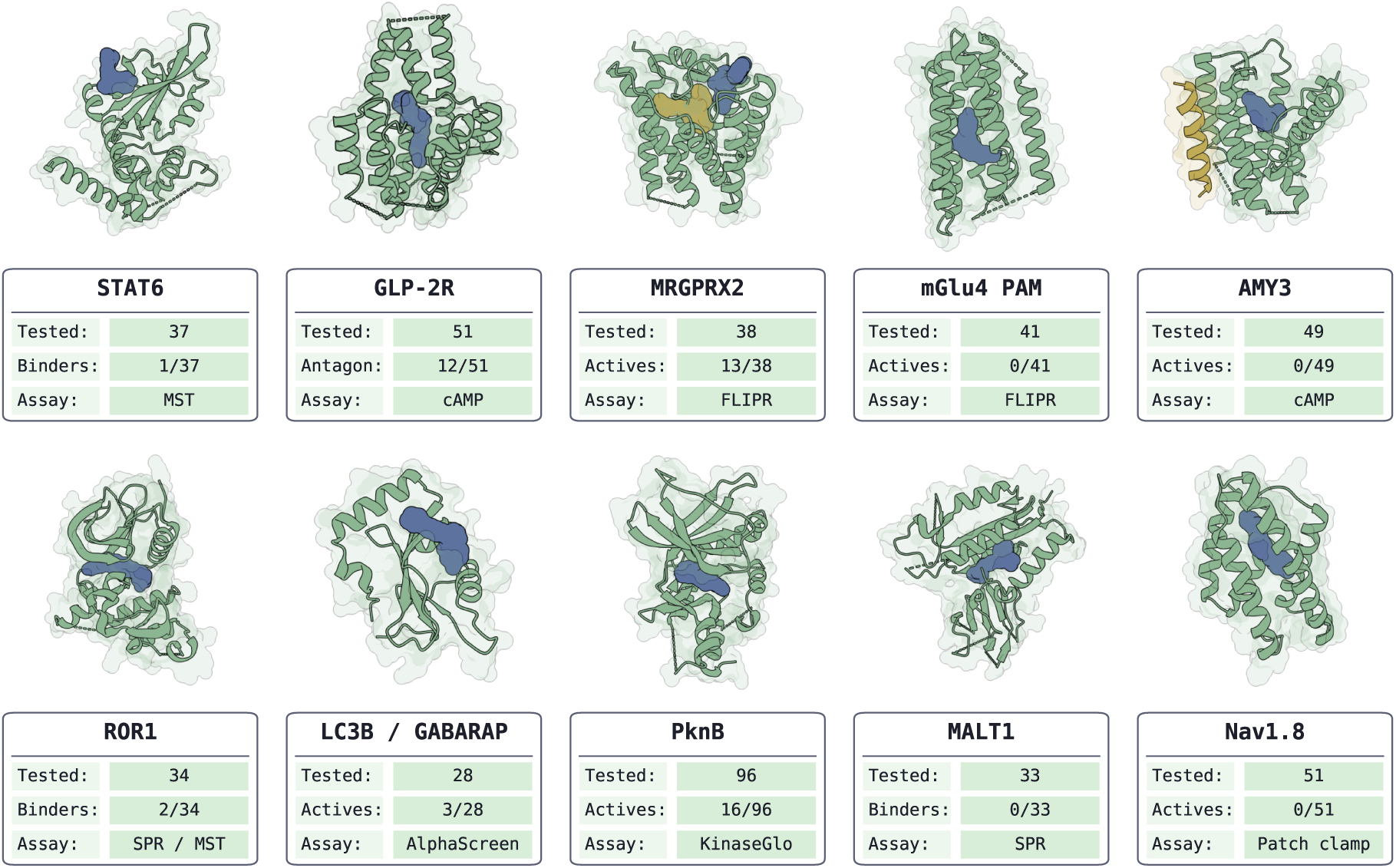
Overview of the prospective virtual-screening campaigns across all targets. For each target, the panel shows the predicted protein-ligand complex together with the number of compounds tested, the number of confirmed actives/binders, and the assays used for screening and follow-up.

## 1 Introduction

Small-molecule hit discovery remains a costly and low-yield endeavor. Conventional approaches typically rely on screening tens of thousands to millions of compounds, either experimentally or through large-scale virtual screening campaigns, to identify a small number of initial hits [Macarron et al., 2011, Gorgulla et al., 2020]. Despite these efforts, HTS hit rates are often extremely low, frequently in the range of 0.01–0.14% and often substantially lower for challenging targets with limited ligand precedent or non-classical binding sites [Zhu et al., 2013]. These challenges are further exacerbated for systems with complex conformational dynamics or poorly defined binding pockets. As a result, the process is both resource-intensive and time-consuming.

While advances in machine learning have improved the accuracy of binding prediction [Zhang et al., 2024], most existing evaluations either rely on retrospective benchmarks [Heikamp and Bajorath, 2013], whose relevance to real-world discovery success is often unclear, or are limited to well-characterized protein classes with abundant ligand data. As a result, it is difficult to assess how well these models generalize to more challenging targets and translate into experimental success [Kearnes, 2021].

Here we present BoltzMol-1, a small-molecule hit-discovery pipeline in which model predictions are used to directly select a small set of compounds for experimental testing. At its core, BoltzMol-1 ranks candidate compounds using the Boltz-2 scoring function [Passaro et al., 2025]: for each protein-ligand pair, Boltz-2 co-folds the complex and predicts a binding affinity together with a small-molecule binding-confidence score which forms a composite binding-confidence signal. This learned signal, rather than classical docking or a physics-based energy function, is the primary driver of compound ranking and selection. BoltzMol-1 is designed around the constraint that all candidates must be readily accessible from commercial catalogs (including synthesis-on-demand virtual space), ensuring that predictions can be rapidly translated into assays through direct purchase, or without lengthy synthesis. By focusing on compounds that can be acquired and tested in short cycles, the approach supports rapid experimental feedback and practical iteration, without relying on large-scale screening.

To further contextualize selected compounds, we evaluate their physicochemical and absorption-related properties using internal ADMET models for kinetic solubility (logS), lipophilicity (logD), and Caco-2 A-to-B permeability. Retrospectively, their predictions show strong correlation with experimental outcomes, indicating that they capture meaningful developability signals and could be deployed prospectively as filters to further reduce the number of compounds requiring experimental testing.

We evaluate BoltzMol-1 against a set of ten targets chosen to represent diverse and challenging hit-discovery settings. Across this panel, we observe prospective success on 6/10 (60%) of targets, identifying experimentally validated binders or functionally active compounds despite modest testing budgets. Notably, for the transcription-factor SH2 domain STAT6, we identify a binder confirmed by SpS and supported by an orthogonal control assay and preliminary SAR. Among GPCR targets, we identify 12 functionally active antagonists for GLP-2R and, for MRGPRX2, 3 antagonists and 10 agonists, all validated in primary assays, confirmed by dose-response profiling, and showing no activity in parental line counter screens. For the pseudokinase ROR1, we identify two confirmed binders validated by both MST and SPR, one of which was further validated with replicate MST and SpS measurements and via preliminary structure-activity relationship (SAR) support. In addition, we expanded previous work investigating the utility of fixed screening libraries and virtual space to two additional targets. We identify, for the ATG8-family proteins LC3B and GABARAP, three inhibitors with dose-response validation, and for serine/threonine protein kinase PknB (Protein Kinase B), sixteen active compounds. These results suggest a path toward reducing the cost and scale of early hit discovery while maintaining robustness across diverse and challenging target classes.

Our screening was carried out using the Boltz Lab platform GUI. This allows a point of entry to the BoltzMol-1 pipeline for non-computationally centred disciplines across the biological sciences. For our non-collaborative prospective screens, we additionally highlight the underlying Boltz API input that can be utilized within computational or agentic workflows in Appendix F.

## 2 Wetlab Results

Here, we summarize each target by describing its biological and therapeutic relevance, outlining the experimental screening strategy, and presenting the screening outcomes. We also report retrospective ADMET measurements for a selected subset of compounds to evaluate whether the prioritized chemistry shows tractable developability properties in addition to target-specific activity. Further methodological details and supporting analyses are provided in the Appendix. Appendix C presents target-specific experimental notes and the complete set of campaign results, Appendix A describes the binding and functional assays used for hit identification and confirmation, Appendix B details the ADMET assays used for profiling, and Appendix E examines target similarity to the training set as well as the similarity of experimentally active compounds to chemistry represented in the training data. Table 1 summarizes the targets, detected interactions, and assay workflows used for experimental validation.

**Table 1:** Summary of prospective validation campaigns across targets, showing detected interaction types and assay workflows used for screening and follow-up.

| Target | # Tested | # Actives | Classification | Experimental Workflow |
| --- | --- | --- | --- | --- |
| STAT6 | 37 | 1 | 1 binder | – SpS<br>– SpS counter-screen<br>– SAR follow-up (+3 binders) |
| GLP-2R | 51 | 12 | 12 antagonists | – cAMP primary screen<br>– cAMP dose-response<br>– cAMP parental control<br>– SAR follow-up (+11 actives) |
| MRGPRX2 | 38 | 13 | 10 agonists<br>3 antagonists | – FLIPR primary screen<br>– FLIPR dose-response<br>– FLIPR parental control<br>– SAR follow-up (+9 actives) |
| ROR1 | 34 | 2 | 1 hit<br>1 binder | – SPR, MST, nanoDSF<br>– SAR follow-up (+3 binders) |
| LC3B / GABARAP | 28 | 3 | 3 inhibitors | – AlphaScreen dose-response |
| PknB | 96 | 16 | 16 actives | – KinaseGlo primary screen<br>– KinaseGlo dose-response |
| mGlu4 PAM | 41 | 0 | N/A | – FLIPR primary screen |
| AMY3 | 49 | 0 | N/A | – cAMP primary screen |
| MALT1 | 33 | 0 | N/A | – SPR |
| Nav1.8 | 51 | 0 | N/A | – Patch clamp |

### 2.1 STAT6

#### Target Class Description

STAT6 is a signal-dependent transcription factor in the JAK–STAT pathway whose SH2 (Src homology 2) domain functions as the central recognition module for pathway activation [Mandal et al., 2015]. The SH2 pocket mediates receptor docking by binding phosphotyrosine motifs on activated cytokine receptors (notably IL-4R*α*/IL-13R*α*1 complexes) and also mediates post-phosphorylation dimer assembly through reciprocal pTyr641–SH2 engagement [Mandal et al., 2015]. Because both receptor recruitment and dimerization depend on this interface, the SH2 site is the key structurally actionable node for direct pathway interruption.

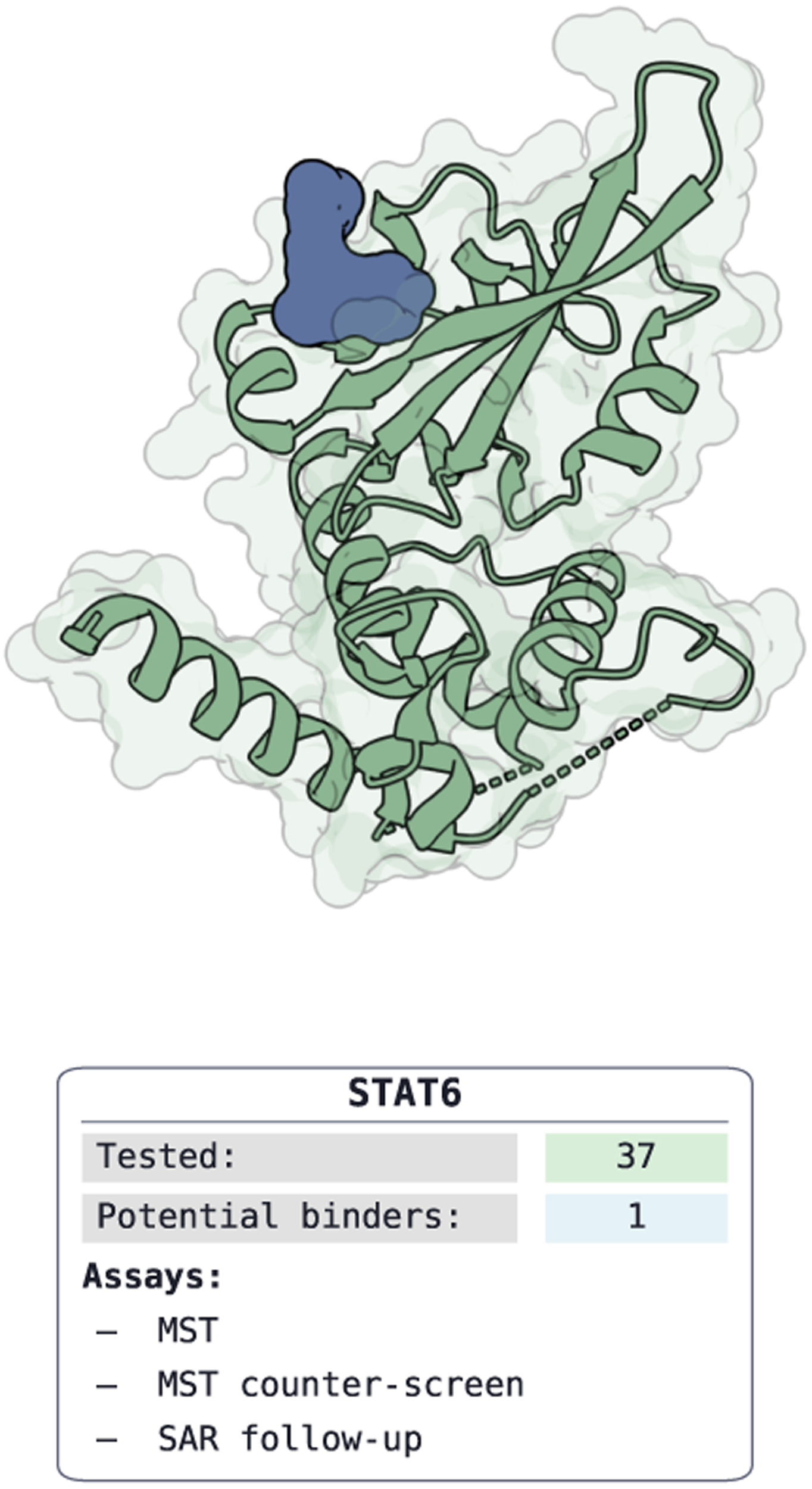

#### Therapeutic Rationale

STAT6 hyperactivation is implicated across both oncology and immunology settings, including lymphoma subtypes and Th2-skewed inflammatory disease (for example, asthma, atopic dermatitis, and broader allergic pathology) [Sharma et al., 2023]. Targeting SH2-mediated dimerization offers a direct mechanism to suppress pathological STAT6 transcriptional programs [Morlacchi et al., 2014]. A further advantage is pathway convergence: SH2-directed inhibitors act downstream of multiple upstream cytokine inputs, creating the potential to reduce signaling from IL-4/IL-13-centered circuits with a single small-molecule intervention point.

#### Results

We screened 37 compounds by SpS (Appendix A.2) and identified one binder with an SpS-measured *K_D_* = 30.3 *µ*M. To assess non-specific binding, we also ran an SpS counter-screen against p38 as a control protein and observed no detectable interaction. Follow-up evaluation of close analogs provided preliminary SAR support, suggesting that the observed activity is chemically meaningful rather than assay-specific. Additional campaign-specific details are provided in Appendix C.1, and the target’s relationship to the training distribution is discussed in Appendix E.1.

### 2.2 GLP-2R Agonist / Antagonist

#### Target Class Description

GLP-2R is a class B GPCR with exceptionally sparse structural coverage. At present, the experimentally determined structural record is effectively limited to a single peptide-bound human GLP-2R–Gs cryo-EM complex (7D68) [Sun et al., 2020], with no apo structure and no small-molecule-bound GLP-2R structure available. As a result, extracellular peptide recognition is defined, but the transmembrane geometry most relevant to oral non-peptide ligand design remains largely unresolved.

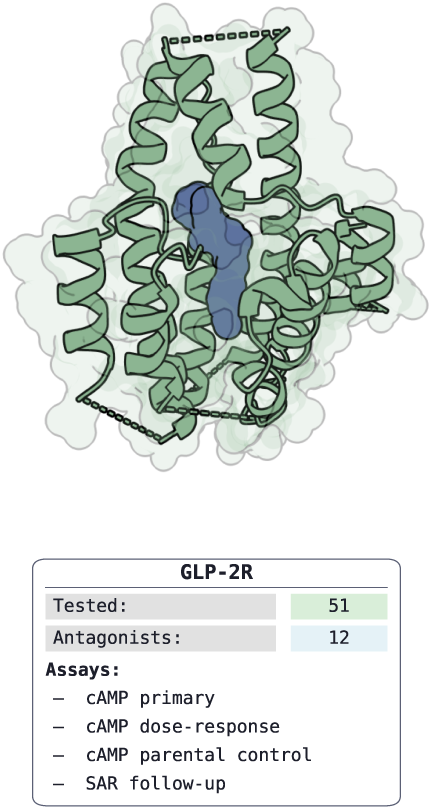

#### Therapeutic Rationale

GLP-2R is clinically validated by peptide agonism. Teduglutide is approved for short bowel syndrome [Jeppesen et al., 2012], and additional long-acting GLP-2 analogs remain in late-stage clinical development [Eliasson et al., 2022]. The remaining unmet need is modality rather than biology: converting this validated intestinal-repair and absorptive program into an orally tractable small-molecule therapy would materially expand accessibility and reduce the burden associated with chronic injectable peptide treatment.

Although initial GLP-2R antagonists would mainly be used in research to dissect GLP-2 signaling and intestinal adaptation, rather than in clinical therapy, emerging biology suggests GLP-2 signaling drives tumor growth in colorectal and small intestinal cancers [Thulesen et al., 2004]. This field of research could be further unlocked by the discovery of potent and selective tool molecules.

#### Results

Compounds were ranked purely by the aforementioned binding-confidence signal to the orthosteric GLP-2 peptide pocket; the screen received no functional bias during target set-up. The model therefore could predict binders where most confident within the orthosteric site, and whether each binder acted as an agonist or antagonist was resolved experimentally. We used a cAMP assay to evaluate agonist and antagonist activity (Appendix A.5). Compounds were first assessed in a primary screen at three concentrations, followed by dose-response confirmation and a parental cell line counter-screen. Using this workflow, we screened 51 compounds and identified 12 antagonists that were active in both the primary and dose-response assays while remaining inactive in the parental cell line counter-screen. The antagonists showed IC_50_ values ranging from 0.07 *µ*M to 24 *µ*M. Additional campaign-specific details are provided in Appendix C.2, and the target’s relationship to the training distribution is discussed in Appendix E.2.

### 2.3 MRGPRX2 Agonist / Antagonist

#### Target Class Description

MRGPRX2 is a class A GPCR expressed prominently on mast cells and implicated in IgE-independent hypersensitivity signaling [Kumar et al., 2021, McNeil et al., 2015]. Structural data now include multiple cryo-EM complexes with peptide and polycationic agonists, together with small-molecule agonist-bound states, and collectively show a shallow, solvent-exposed orthosteric pocket [Cao et al., 2021]. Although the March 2025 PSB-172656 study disclosed a potent antagonist chemotype, its receptor-bound pose was inferred by docking rather than solved in an antagonist-bound complex [Al Hamwi et al., 2025]. The inactive small-molecule recognition mode of MRGPRX2 therefore remains unresolved despite increasing medicinal-chemistry activity around the target.

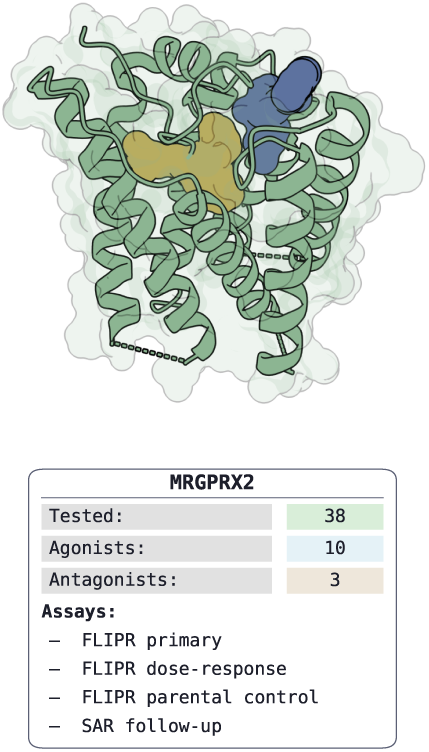

#### Therapeutic Rationale

MRGPRX2 is therapeutically compelling for both efficacy and safety reasons. It contributes to chronic urticaria, itch, and broader mast cell-driven inflammatory disease, while also mediating pseudo-allergic reactions triggered by numerous approved drugs, including fluoroquinolones, opioids, and neuromuscular blockers [Kolkhir et al., 2023]. Clinical advancement of oral antagonists such as EVO756 and INCB000262/EP262 further supports tractability and highlights the opportunity for selective small molecules that suppress pathological mast-cell activation without relying on antibody-based intervention [Evommune, Inc., 2025].

Identification of agonistic binders would have downstream implications for the safety evaluation against this target for the assessment of MRGPRX2 as an anti-target in clinical safety, where antagonists carry intrinsic therapeutic benefit.

#### Results

Compounds were ranked purely by their predicted binding-confidence signal, with no input-conditioning to elicit a desired functional response. We used a FLIPR assay to measure both agonist and antagonist activity via calcium efflux (Appendix A.6). Compounds were first evaluated in a primary screen at three concentrations, followed by dose-response confirmation and a parental cell line counter-screen. Using this workflow, we screened 38 compounds with the resulting split into agonist and antagonist classification emerging from the FLIPR assay rather than from the ranking itself. BoltzMol-1 identified 10 agonists and 3 antagonists that were active in both the primary and dose-response assays and inactive in the parental cell line counter-screen. The agonists showed EC_50_ values ranging from 2.2 *µ*M to 18 *µ*M, while the antagonists showed IC_50_ values ranging from 9.3 *µ*M to 19 *µ*M. Additional campaign-specific details are provided in Appendix C.3, and the target’s relationship to the training distribution is discussed in Appendix E.3.

### 2.4 ROR1

#### Target Class Description

ROR1 is a receptor tyrosine kinase-like orphan receptor whose intracellular kinase domain is a pseudokinase rather than a canonical active kinase [Sheetz et al., 2020]. Structural work shows that the ATP pocket is sterically occluded by aromatic side chains via degenerated catalytic motifs (mutated DFG and HRD sites) resulting in the activation loop adopting an autoinhibited-like arrangement [Sheetz et al., 2020]. This architecture makes conventional ATP-site engagement atypical and shifts tractable small-molecule binding toward a back-pocket adjacent to the blocked nucleotide site which remains accessible, rendering a type-II inhibitor binding mode a more tractable small molecule strategy [Mendrola et al., 2013].

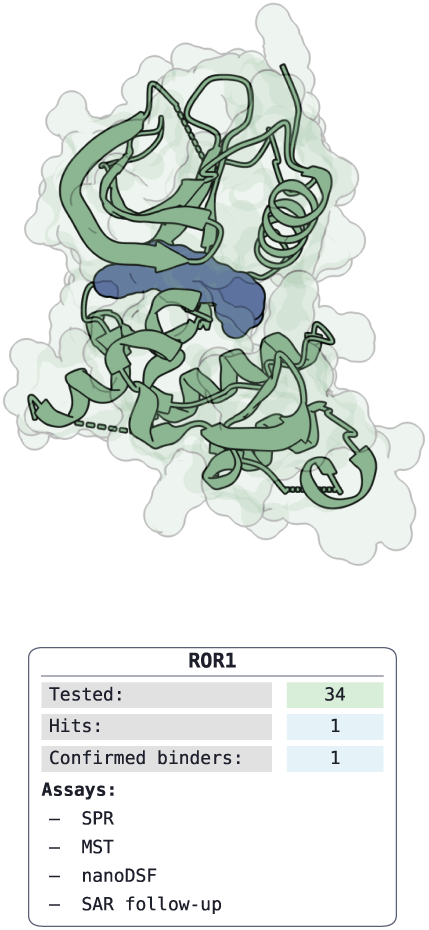

#### Therapeutic Rationale

ROR1 regulates embryonic and fetal development and is expressed at low levels in most normal adult tissues [Borcherding et al., 2014]. However, its overexpression in adult tissue is strongly associated with tumorigenesis and metastasis in hematologic and solid tumor settings [Zhao et al., 2021]. Because the pseudokinase domain is not driven by classical catalytic turnover, the therapeutic hypothesis is non-catalytic. Ligands may function as conformational modulators that perturb receptor-dependent scaffolding and non-canonical signaling states (for example, WNT5a-associated pathways) [Konopelski Snavely et al., 2023], thereby reducing tumor-supportive signaling without relying on phospho-transfer inhibition.

#### Results

We screened the prioritized compounds using three orthogonal biophysical assays: SPR, MST/SpS, and nanoDSF (Appendix A.1, Appendix A.2, and Appendix A.3). Across 34 tested compounds, we identified one confirmed binder that was active in all three assays and showed a *K_D_* = 5.13 *µ*M by SPR. Follow-up evaluation of close analogs provided preliminary SAR support, indicating that the observed activity is chemically meaningful rather than assay-specific. We also identified a second putative binder showing binding in SPR and MST with a *K_D_* = 10.1 *µ*M in SPR, although this compound did not show thermal stabilization in nano-DSF. Additional campaign-specific details are provided in Appendix C.4, and the target’s relationship to the training distribution is discussed in Appendix E.4.

### 2.5 LC3B and GABARAP

*Experiments by Madison Maiorano and Joshua Kritzer*.

#### Target Class Description

LC3B and GABARAP are ATG8-family ubiquitin-like modifiers central to autophagy [Martens and Fracchiolla, 2020]. Both share the conserved *β*-grasp fold and a LIR docking site built from HP1, the aromatic W/F/Y-recognition pocket, and HP2, the aliphatic L/I/V-recognition pocket that anchors cargo-receptor motifs such as p62, NBR1, OPTN, and NDP52 [Wirth et al., 2019]. Where LC3B and GABARAP differ matters for selectivity: HP1 is tighter in LC3B and more open in GABARAP, and the electrostatics around the docking surface diverge enough to support potential subfamily-selective ligands [Wirth et al., 2019].

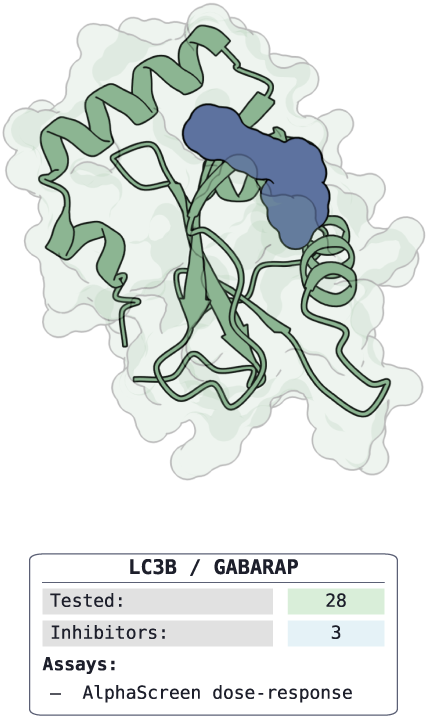

#### Therapeutic Rationale

Autophagy inhibition is an established strategy for sensitizing cancers to chemotherapy and DNA-damaging agents [Verbaanderd et al., 2017]. LC3B/GABARAP proteins are mechanistically central to this approach and mediate protein-protein interactions at every step of the autophagy pathway [Wirth et al., 2019]. Existing clinical-stage autophagy inhibitors such as chloroquine and hydroxychloroquine act lysosomotropically, and suffer from poor pharmacokinetics and high toxicity at therapeutic doses [Mauthe et al., 2018]. Targeting the LIR docking surface directly offers a more specific mechanism: blocking cargo receptor engagement at the autophagosome membrane rather than disrupting lysosomal function globally. Small molecules that modulate the LC3B or GABARAP docking surface selectively could therefore serve as both mechanistic tools to dissect these distinct branches of ATG8-family biology and, if developed further, as starting points for more targeted autophagy-modulating therapeutics.

#### Results

28 compounds predicted by the BoltzMol-1 to bind LC3B or GABARAP were purchased and tested in a competitive AlphaScreen assay measuring displacement of biotinylated peptide tracers (FYCO1S for LC3B; K1 for GABARAP) from His-tagged recombinant protein (Appendix A.9). Three compounds were classified as inhibitors, with IC_50_ values showing activity against GABARAP in the 15 to 28 *µ*M range and LC3B 15 to 40 *µ*M respectively. Additional campaign-specific details are provided in Appendix C.5, and the target’s relationship to the training distribution is discussed in Appendix E.5.

### 2.6 PknB

*Experiments by Anthony Gitter and Nathan Wlodarchak*.

#### Target Class Description

PknB is a transmembrane receptor-like serine/threonine protein kinase (STPK) belonging to a family of eleven eukaryotic-like STPKs encoded in the *Mycobacterium tuberculosis*(Mtb) genome. The intracellular kinase domain adopts the canonical bilobal fold of eukaryotic serine/threonine kinases, with an N-terminal lobe housing the ATP-binding site and a C-terminal lobe responsible for substrate recognition and catalysis. Crystal structures of the PknB kinase domain reveal an atypical ‘DFG-in, C-helix out’ active conformation, that present conformational specifics for binding pose generation, despite being a well-defined adenine-binding pocket that accommodates ATP and ATP-competitive inhibitors [Wlodarchak et al., 2018]. The extracellular region consists of four tandem PASTA (penicillin-binding protein and serine/threonine kinase associated) domains that bind muropeptide fragments of peptidoglycan, coupling cell wall remodeling signals to intracellular kinase activation. The architecture of a structurally well-characterized, catalytically active ATP-binding pocket connected to a biologically meaningful extracellular sensor makes PknB a tractable small-molecule target with a clear structural basis for inhibitor design.

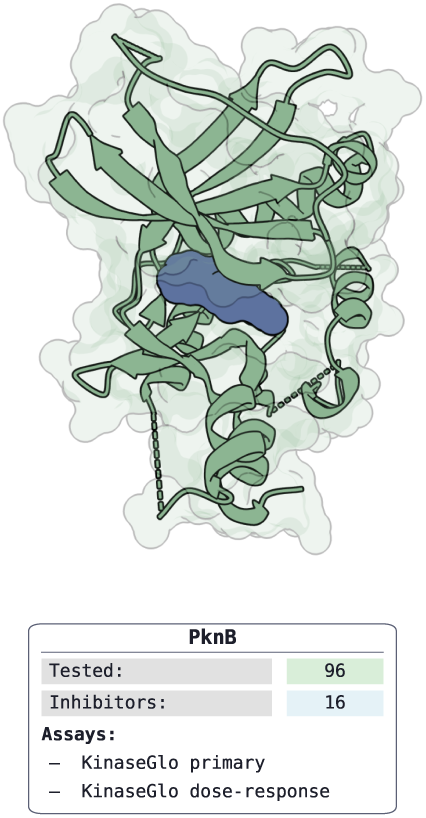

#### Therapeutic Rationale

PknB is essential for sustained mycobacterial growth. Conditional depletion of PknB causes growth arrest, morphological defects, and loss of acid-fast staining, reflecting disruption of cell wall mycolate synthesis. Its substrates span cell division regulators, penicillin-binding proteins, and metabolic enzymes, placing PknB at the apex of signaling cascades that coordinate cell shape, division, and adaptation to the host environment. Critically, PknB dependence is conserved across drug-susceptible and multidrug-resistant (MDR) clinical isolates, with the ATP/ADP-binding domain showing high sequence conservation across resistant strains. This makes PknB an attractive target in the context of the global MDR-TB crisis. Structural divergence between the Mtb STPK family and human kinomes around the ATP-binding pocket provides a selectivity window that has been exploited in medicinal chemistry campaigns. Identification of selective ATP-competitive inhibitors with antimycobacterial activity and minimal inhibition of human kinases have been reported, establishing PknB as a compelling target for novel anti-tuberculosis drug discovery [Wlodarchak et al., 2018].

#### Results

96 compounds from Enamine were purchased and tested in a biochemical KinaseGlo activity assay measuring ATP consumption by PknB 1–331. Sixteen compounds were classified as confirmed actives, with IC_50_ values ranging from 23.2 to 399 *µ*M. K*_i_* values were derived assuming competitive inhibition, ranging from 6.8 to 118 *µ*M. The reference inhibitor GSK690693 returned an IC_50_ of 0.47 *µ*M, in agreement with published values and confirming assay performance [Wlodarchak et al., 2018]. Additional campaign-specific details are provided in (Appendix C.6), and the target’s relationship to the training distribution is discussed in (Appendix E.6).

### 2.7 mGlu4 PAM

#### Target Class Description

mGlu4 is a class C GPCR for which practical small-molecule modulation is typically achieved through transmembrane allosteric sites rather than the orthosteric glutamate-binding Venus flytrap domain [Gutzeit et al., 2019]. Unlike class A GPCRs where orthosteric ligands bind within the 7TM bundle, class C receptors function as obligate dimers with glutamate binding in the extracellular VFT domain [Gutzeit et al., 2019]. Positive allosteric modulation is therefore a conformationally selective problem that depends on stabilizing the right receptor state rather than simply occupying a canonical active site.

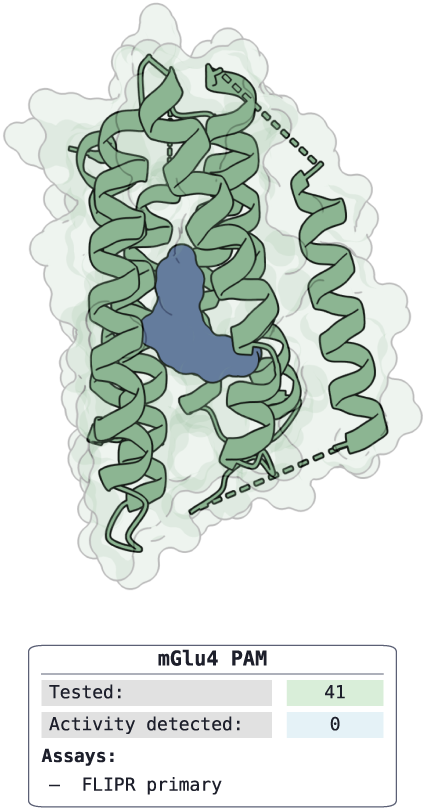

Recent cryo-EM structures have revealed that mGlu4 PAMs can engage at least two distinct allosteric sites within the 7TM region. These are either an intra-subunit pocket spanning TM3, TM5, TM6, TM7 (exemplified by PDB 8WGB) [Huang et al., 2024] or an asymmetric dimer interface between subunits (exemplified by VU0364770 in PDB 8JD6) [Wang et al., 2023]. The functional outcome (whether a compound acts as a PAM, agonist, or NAM) depends on subtle differences in binding pose and the specific receptor conformational states stabilized.

#### Therapeutic Rationale

mGlu4 positive allosteric modulators have been pursued primarily for Parkinson’s disease, where they may reduce pathological overactivity in the indirect pathway of the basal ganglia without directly replacing dopamine [Charvin, 2018]. Preclinical studies demonstrated that mGlu4 PAMs such as VU0364770 and ADX88178 improve motor function in rodent models of parkinsonism, both alone and in combination with L-DOPA [Jones et al., 2012].

Foliglurax (PXT-002331) became the first mGlu4 PAM to reach Phase II clinical trials for Parkinson’s disease. However, the AMBLED trial (2020) failed to meet its primary endpoints, with the compound unable to sufficiently distinguish itself from placebo despite showing acceptable safety [Rascol et al., 2022]. This clinical setback highlighted the translational challenges in this target class and raised questions about whether preclinical models adequately predict human efficacy.

The target remains of interest as a neuropharmacology tool and as a test case for how well computational pipelines handle class C GPCR allostery, where subtle structural differences can strongly influence subtype selectivity and efficacy mode.

#### Results

We screened 41 compounds by FLIPR assay (Appendix A.6) and detected no activity. Additional target-specific analysis is provided in Appendix E.7.

### 2.8 Amylin 3 Receptor Agonist

#### Target Class Description

AMY3 is a heteromeric amylin receptor formed by the calcitonin receptor in complex with RAMP3 [Cao et al., 2022]. This receptor-complex architecture places it in the class B GPCR signaling family while adding RAMP-dependent pharmacology that can materially reshape ligand recognition and assay behavior relative to the calcitonin receptor alone [Hay and Pioszak, 2016].

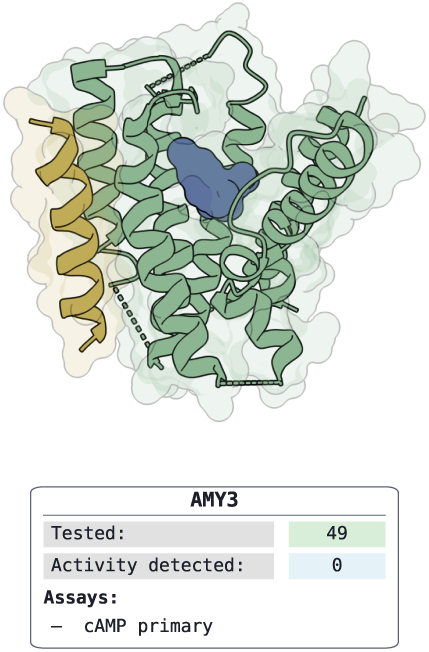

#### Therapeutic Rationale

Agonism at amylin-family receptors is therapeutically attractive for metabolic disease because it can influence satiety, gastric emptying, and broader glucoregulatory physiology [Boyle et al., 2022]. A small-molecule entry point for AMY3 would therefore be interesting both as a potential therapeutic direction and as a mechanistic probe of receptor-subtype contributions within the calcitonin/amylin receptor family. While GLP-1 receptor agonists act primarily via incretin-mediated insulin secretion and central appetite suppression, amylin receptor agonism engages complementary and partially distinct neural circuits that regulate both homeostatic and hedonic feeding [Boyle et al., 2022]. This mechanistic differentiation has driven clinical interest in combining amylin and GLP-1 pathways, exemplified by the unimolecular GLP-1/amylin co-agonist amycretin, which demonstrated additive weight loss beyond either modality alone in early-phase trials [Dahl et al., 2025].

#### Results

We screened 49 compounds by cAMP assay (Appendix A.5) and detected no activity. Additional target-specific analysis is provided in Appendix E.8.

### 2.9 MALT1

#### Target Class Description

MALT1 is the only human paracaspase and a dual-function signaling node in the CARD11-BCL10-MALT1 (CBM) complex, combining scaffold-mediated NF-*κ*B assembly with catalytic cleavage of regulatory substrates [Afonina et al., 2015, Lu et al., 2018]. This biology creates two structurally actionable modes: a catalytic cysteine-containing active site and a distinctive allosteric pocket centered on Trp580, which is not recapitulated in classical caspases. The allosteric mechanism has been structurally and pharmacologically validated, including through the MLT-747/MLT-748 chemotype series. These potent, selective allosteric inhibitors bind the Trp580 pocket (IC_50_ = 5 nM) and lock the protease in an inactive conformation, with co-crystal structures deposited in the PDB, establishing MALT1 as a tractable but still under-diversified small-molecule target [Quancard et al., 2019].

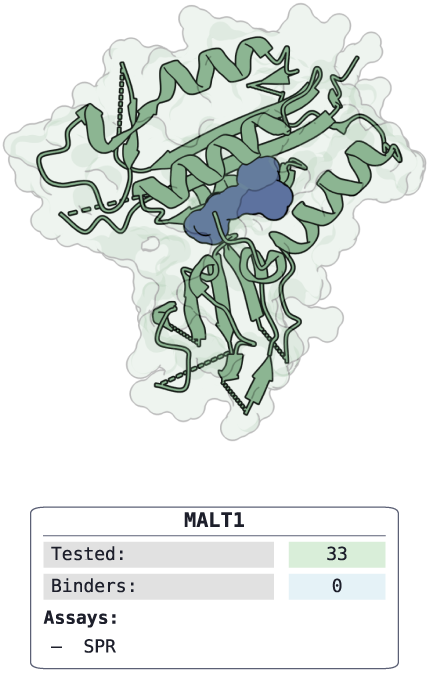

#### Therapeutic Rationale

MALT1 has strong disease genetics and pathway dependency in B-cell malignancies, particularly NF-*κ*B-addicted contexts such as ABC-DLBCL and translocation-driven MALT lymphoma, and broader relevance across additional hematologic and selected solid-tumor settings [Fontan et al., 2012]. Beyond tumor-cell intrinsic signaling blockade, MALT1 inhibition may provide a second oncology mechanism by destabilizing suppressive tumor-associated Treg programs, potentially enhancing anti-tumor immunity [Rosenbaum et al., 2019]. Despite no approved small molecules to date, multiple programs are now advancing in the clinic [Fontán and Melnick, 2012, Zhang et al., 2025], making MALT1 an attractive target that represents a rare convergence of precision-oncology and immune-modulation rationale within one target class.

#### Results

We screened 33 compounds by SPR (Appendix A.1) and identified no evidence of binding. Additional target-specific analysis is provided in Appendix E.9.

### 2.10 Nav 1.8

#### Target Class Description

Nav1.8 (SCN10A) is a voltage-gated sodium channel preferentially expressed in peripheral sensory neurons, particularly nociceptors of the dorsal root ganglia [Xie, 2025]. Unlike the nine mammalian Nav isoforms that share *>* 50% sequence identity and a conserved pore architecture, Nav1.8 contains a unique sequence motif in Voltage-Sensing Domain 2 (VSD2) that enables isoform-selective pharmacology [Xie, 2025, Wood and Iseppon, 2025]. Structural work reveals that the extracellular S3-S4 loop of VSD2 harbors a “KKGS” tetrapeptide sequence (K748, K749, G750, S751) found exclusively in Nav1.8. Other isoforms carry divergent residues at these positions. Additionally, V746 in the S3 helix is valine in Nav1.8 but leucine in other subtypes, creating a smaller side-chain that permits closer ligand engagement [Osteen et al., 2025]. This architecture enables allosteric biomolecules to bind the VSD2 extracellular cleft and stabilize the channel’s resting (closed) state, blocking sodium conductance without occluding the central pore.

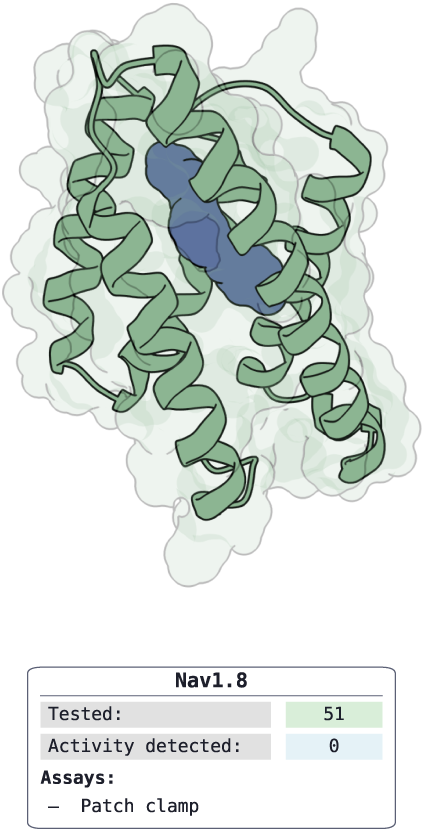

#### Therapeutic Rationale

Nav1.8 is a well validated peripheral pain target. Its restricted expression to sensory neurons absent from CNS, cardiac, and skeletal muscle tissue provides a therapeutic window unavailable to earlier non-selective sodium channel blockers [Jones et al., 2023]. Genetic validation includes gain-of-function SCN10A mutations that cause painful neuropathy, and SNPs that bias human pain sensitivity [Faber et al., 2012]. Suzetrigine (VX-548), a VSD2-targeting inhibitor, demonstrated efficacy in Phase 3 trials for acute pain and received FDA approval in January, 2025, establishing proof-of-concept for this mechanism as a non-opioid analgesic [U.S. Food and Drug Administration, 2025, Bertoch et al., 2025]. The therapeutic mechanism-of-action for VX-548 centers on state-dependent allosteric VSD2 modulation via trapping the channel in a resting conformation, reducing neuronal excitability in hyperactive nociceptors without affecting normal sensory function or other Nav-dependent tissues [Osteen et al., 2025].

#### Results

We screened 51 compounds by patch-clamp assay (Appendix A.4) and detected no activity. Additional target-specific analysis is provided in Appendix E.10.

### 2.11 Experimental ADMET Results

We subjected a subset of compounds emerging from the virtual-screening campaigns to experimental ADMET profiling to assess whether apparently promising screening matter also possessed acceptable developability properties. Despite being drawn from drug-like screening libraries, many of these compounds showed unfavorable measured profiles, including poor solubility, unfavorable lipophilicity, or limited permeability. These observations motivate the need for an ADMET filtering layer early in the selection process, so that compounds are prioritized not only for target activity but also for their likelihood of supporting downstream progression. For this subset, we experimentally measured lipophilicity, kinetic solubility, and Caco-2 permeability; assay details are provided in Appendix B, with individual protocol summaries in Appendix B.1, Appendix B.2, and Appendix B.3.

**Figure 2:**
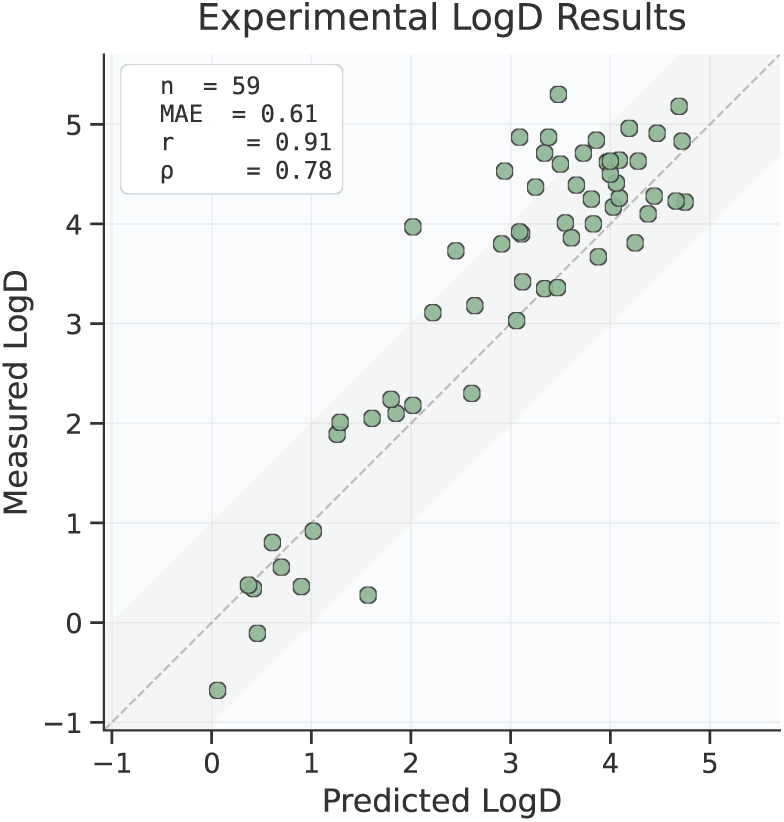
Retrospective performance of the LogD model on prospectively profiled compounds. Predicted versus measured LogD for *n* = 59 compounds. The dashed line indicates the identity.

#### LogD

Across 59 compounds, predicted and measured LogD agreed with Pearson *r* = 0.91 and MAE = 0.61 log units across the full assay range. Performance is consistent with our retrospective benchmark, where LogD behaved as a comparatively saturated task, and the prospective measurements confirm that the model produces reliable continuous estimates suitable for downstream lipophilicity-based triage.

**Figure 3:**
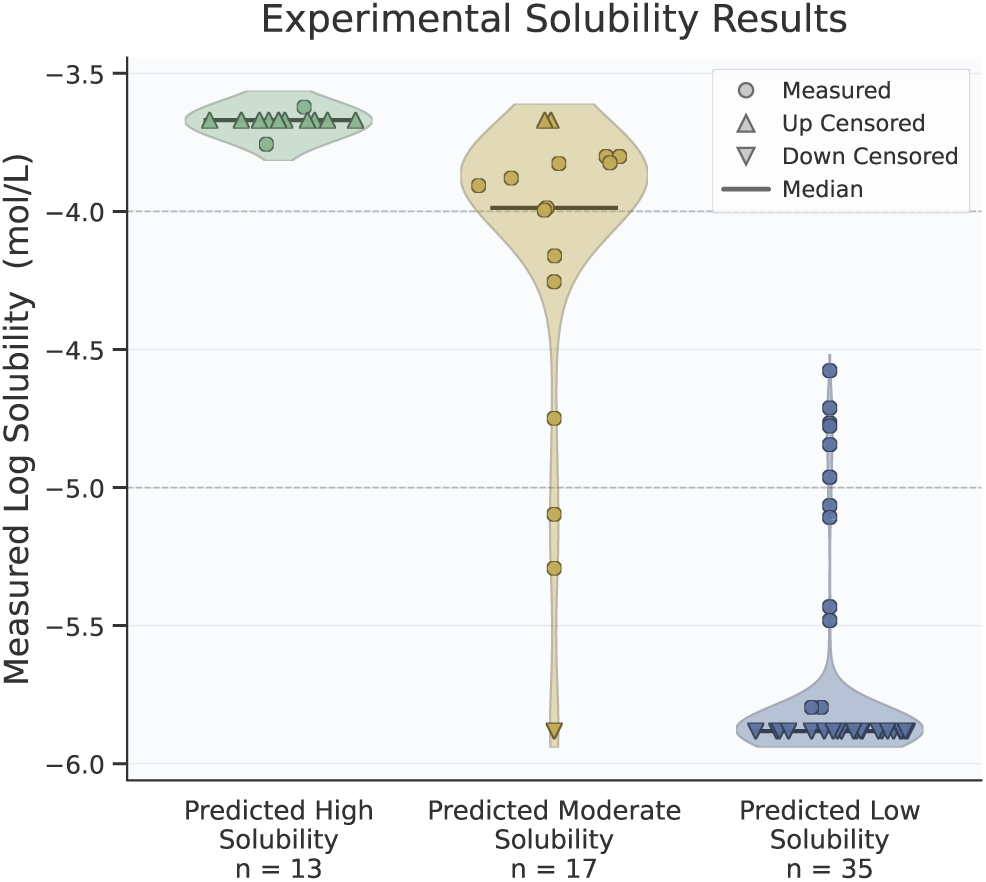
Retrospective performance of the KSol model on prospectively profiled compounds. Measured KSol distributions are shown for compounds grouped by predicted solubility bucket. Up- and down-triangles denote censored measurements at the assay’s upper and lower detection limits, respectively.

#### KSol

The KSol model returns a three-bin call rather than a continuous value, calibrated for high-precision identification of soluble compounds and high-recall exclusion of insoluble compounds. Of the 12 compounds placed in the predicted-high bin, 11 were confirmed at the assay’s upper detection limit and all 12 confirmed as highly soluble with measured KSol above *−*4 log mol/L. Of the 34 compounds placed in the predicted-low bin, 29 were confirmed as poorly soluble with measured KSol below *−*5 log mol/L, 23 of these pinned at the lower detection limit, and none mis-classified as highly soluble. The predicted-moderate bin covers the intermediate range as intended: of its 17 compounds, 10 were highly soluble with measured KSol above *−*4 and only 3 were poorly soluble with measured KSol below *−*5, with the remainder distributed between. Together, the strong separation between the high and low bins, combined with the moderate bin appropriately absorbing the most ambiguous chemistry, supports the use of the model as an early triage filter: compounds in the low solubility bin can be discarded in a virtual screen and compounds in the high solubility bin can be advanced with high confidence.

#### Caco-2 permeability

We restrict the analysis to compounds that are sufficiently soluble under the assay conditions, since insoluble compounds do not reach the nominal dosing concentration in the donor compartment and produce apparent permeability readouts that reflect precipitation rather than membrane transport; consistent with this, mass balance recovery for these compounds is very low. For the 8 compounds with measured efflux ratio above 2, likely efflux substrates whose net A*→*B permeability reflects active transport in addition to passive flux, we additionally ran the assay in the presence of the dual P-gp/BCRP efflux inhibitor GF120918. Across the resulting 16 compounds, predicted and measured Caco-2 A*→*B apparent permeability agreed with r = 0.65 and MAE = 0.21 (see Figure 4); without the inhibitor runs, agreement on the same set drops to r = 0.53 and MAE = 0.37. The model provides useful permeability estimates for triage, though it does not consistently identify which compounds are efflux substrates, and orthogonal information remains valuable for resolving this regime.

**Figure 4:**
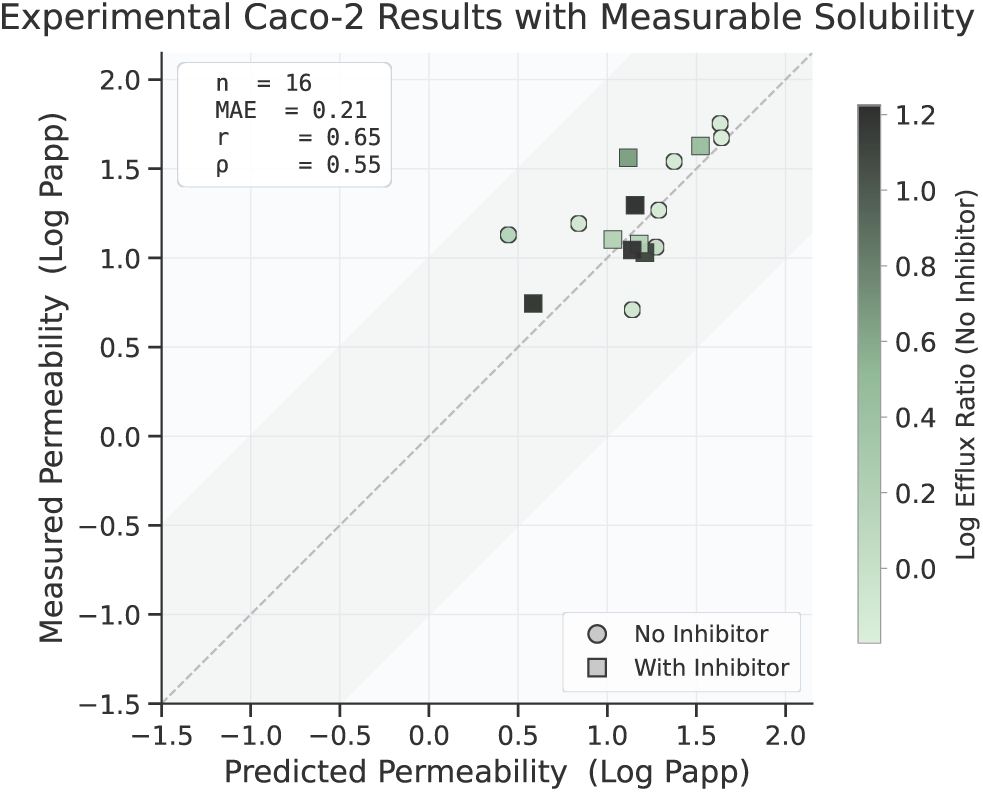
Retrospective performance of the Caco-2 permeability model on prospectively profiled compounds with measurable compound recovery. Predicted versus measured A*→*B apparent permeability, distinguishing assays performed without an efflux inhibitor (circles) from those performed with the dual P-gp/BCRP efflux inhibitor GF120918 (squares). Marker colour indicates measured log efflux ratio.

Together, these results show that developability cannot be assumed even within drug-like screening collections and support the use of ADMET models as a pre-filtering step before experimental advancement.

## 3 Methods

### 3.1 Virtual Screening

The BoltzMol-1 virtual screening pipeline is built around an optimized version of Boltz-2 [Passaro et al., 2025] to better support prospective hit discovery, and available through the Boltz API and the Boltz Lab platform. Four modifications distinguish the screening configuration used here from the default Boltz-2 setup.

First, we tune the inference-time hyperparameters and recombine the model’s outputs into a single ranking score. Boltz-2 produces multiple complementary signals, including multiple predicted binding affinities and probabilities, structural confidence estimates. We calibrate their relative weighting on internal retrospective benchmarks designed to mimic the prospective screening setting, where the model must discriminate true binders from large pools of decoy chemistry drawn from the same catalog distribution. The resulting composite score is the primary basis for compound ranking.

Second, we apply a custom set of medicinal-chemistry filters tailored to the developability requirements of an early hit-finding campaign. These filters extend beyond standard PAINS and reactive-group exclusions [Baell and Holloway, 2010] to also remove compounds with property profiles unlikely to support downstream progression, such as extreme molecular weight, excessive flexibility, or substructures associated with assay interference in the specific readouts we deploy. This filtering step is applied to the catalog before scoring, ensuring that compute is spent ranking compounds that are plausible starting points for medicinal chemistry rather than chemically problematic matter that would be discarded post hoc.

Third, we identify and exclude a set of structures we term *virtual PAINS*: compounds that the model systematically scores as strong binders across unrelated targets, independent of any genuine target-specific interaction signal. These molecules act as interference compounds for the in-silico ranking process itself, inflating top-of-list enrichment with chemistry that is unlikely to reflect true binding. We enumerate virtual PAINS by aggregating Boltz-2 scores across a diverse panel of held-out targets and flagging compounds that consistently appear in the top ranks regardless of target identity. These flagged structures, together with their close analogs, are removed from the screening pool prior to ranking.

Lastly, we optimize the runtime of Boltz to allow faster inference and scale up inference-time compute. Reducing the per-compound cost of a forward pass has a direct effect on screening throughput, allowing us to evaluate substantially larger catalog slices under a fixed compute budget.

Together, these components define BoltzMol-1 as an end-to-end small-molecule pipeline that integrates affinity prediction, medicinal-chemistry filtering, and ADMET scoring into a single ranking workflow. The pipeline supports two distinct deployment modes. In the first, BoltzMol-1 is run against a fixed, enumerable compound library, scoring and ranking a predefined catalog in full. In the second, it is deployed against a combinatorial, synthesizable chemical space far too large to enumerate exhaustively; here BoltzMol-1 leverages a generative active-learning workflow that iteratively proposes, scores, and refines candidates to explore the space efficiently under a fixed compute budget, concentrating evaluation on the most promising regions rather than scoring uniformly at random.

We deploy BoltzMol-1 using the WuXi Off-the-Shelf library in fixed-library mode for the receptor, enzyme, transcription-factor, ion-channel, and GPCR campaigns, and in the generative active-learning mode against Enamine REAL [Enamine Ltd., 2024] for the LC3B/GABARAP campaigns. PknB results use the BoltzMol-1 pipeline without its standard medicinal-chemistry filters, but with additional triage as outlined in Appendix C.6. For each target, we select the top N available compounds, where N is set by the per-target experimental budget (28-96 compounds; see Table 1).

### 3.2 ADMET Models

A promising hit must be more than simply a good binder to the target of interest. Even high-affinity compounds fail further along in development if they are insufficiently soluble to reach the target at relevant concentrations, too lipophilic to avoid off-target liabilities, or too poorly permeable to be absorbed into the bloodstream or access intracellular compartments [Waring et al., 2015]. The earliest stage of medicinal chemistry therefore relies on a small set of physicochemical and in vitro properties, often grouped under the informal label of Tier 1 ADME, to assess compounds before more resource-intensive in vitro and in vivo assays [Siramshetty et al., 2021].

We focus here on three. Aqueous solubility (LogS) measures the concentration at which a compound precipitates out of buffered aqueous solution at physiological pH. The distribution coefficient (LogD) measures the equilibrium partitioning of a compound between octanol and aqueous buffer at physiological pH and serves as the standard summary of effective lipophilicity for ionizable molecules [Waring, 2010]. Caco-2 A*→*B permeability measures the rate at which a compound is transported across a directionally polarized monolayer of cultured cells from the apical (intestinal) to basolateral (bloodstream) side, serving as an in vitro proxy for intestinal absorption [Press and Di Grandi, 2008].

#### 3.2.1 Overview

For an ADMET model to provide value within a virtual screen, it must function as a high-precision triage filter that generalizes across a large and diverse region of chemical space. Meeting this standard is difficult because public ADMET data is heterogeneous, noisy, and often assembled from assays with inconsistent protocols, endpoint definitions, and standardization practices [Parrondo-Pizarro et al., 2025]. Recent community efforts [MacDermott-Opeskin et al., 2026] have organized retrospective evaluations on unblinded lead-optimization campaigns, where compounds are typically generated under a single assay protocol against one or two chemical series [Fang et al., 2023]. Strong performance in this within-series regime can reflect genuine utility for lead optimization, but it does not directly establish performance in the hit-triage setting, where predictions are required for compounds with no close training neighbors.

A separate line of public benchmarks targets out-of-distribution generalization explicitly, most prominently the ADMET tasks in the Therapeutics Data Commons (TDC), which releases scaffold-based splits intended to estimate performance on unseen chemistry [Huang et al., 2021]. We find that these splits are not strict enough in practice: incomplete standardization places identical or near-identical molecules on opposite sides of the split, and scaffold separation does not prevent local analogs, compounds differing by a single substituent that medicinal chemists would treat as the same SAR point, from crossing the train/test boundary. Across the nine TDC ADMET regression tasks, between 43% and 80% of test compounds have either a globally similar or a locally analogous neighbor in the training set under the criteria we define. We therefore argue that the relevant notion of leakage for medicinal-chemistry data combines global similarity with local analog overlap, capturing relationships scaffold splits miss in both regimes. The full analysis, including a leakage taxonomy, worked examples, and per-task prevalence numbers, is reported in Appendix D.1. Reported performance on these splits therefore overstates generalization to the chemistry encountered in hit discovery and virtual screening.

#### 3.2.2 Data Curation

The core data corpus relies on a combination of ChEMBL [Zdrazil et al., 2024] and GOSTAR [Excelra (formerly GVK BIO)], which together provide measurements for LogS, LogD, and Caco-2 A*→*B permeability across a broad range of drug-like chemical space.

We additionally collate data from ChEMBL and GOSTAR on a set of auxiliary permeability-related endpoints. The use of these endpoints in a multi-task setup, including how task transfer to the primary endpoints is structured, is described in Section 3.2.3. The Caco-2 assay is traditionally run bidirectionally, apical-to-basolateral (A*→*B) and basolateral-to-apical (B*→*A), because the rate of transport is asymmetric: efflux transporters expressed at the apical membrane, primarily P-glycoprotein (MDR1) and to a lesser extent BCRP and MRP2, actively pump substrates from the cell back to the apical side [Saaby and Brodin, 2017]. The efflux ratio (B*→*A divided by A*→*B) therefore quantifies the contribution of active efflux to net permeability, with values above 2 indicating that the compound is a substrate for one or more apical efflux pumps. We collate data on all three readouts for training. We also include the analogous bidirectional panel run in MDCK and MDCK-MDR1 cells. MDCK is a canine kidney epithelial line widely used as a faster-growing alternative to Caco-2, with low endogenous expression of efflux transporters. MDCK-MDR1 is its MDR1-transfected variant [Polli et al., 2001]. Finally, we include PAMPA, a parallel-artificial-membrane assay with no cells or transporters that provides a clean readout of passive permeability alone [Kansy et al., 1998].

From this corpus we construct training, validation, and held-out test sets for the three primary endpoints: LogS, LogD, and Caco-2 A*→*B. Validation and test sets are constructed on a per-document basis and required to be free of leakage to any training compound under both global similarity and local analog criteria. We additionally augment the training set with data drawn from public benchmarks and competitions while maintaining our criteria for strict out-of-distribution evaluation. The full curation methodology, including the leakage criteria and external dataset integration, is described in Appendix D.2.

#### 3.2.3 Modeling

We adopt a multi-task learning framework in which a dedicated model is trained for each primary endpoint, optimizing prediction quality on that endpoint while using auxiliary endpoints as transfer signal during training [Kearnes et al., 2017]. The auxiliary set is chosen to reflect the physical-chemistry hierarchy among the endpoints rather than to maximize data volume indiscriminately. The LogD model is trained on LogD alone; the LogS model is trained jointly on LogS and LogD, since LogD is the dominant physicochemical determinant of aqueous solubility for ionizable molecules; and the Caco-2 A*→*B model is trained jointly on the full panel of permeability and lipophilicity endpoints described above (Caco-2 A*→*B, B*→*A, and efflux ratio; the analogous MDCK and MDCK-MDR1 panels; PAMPA and LogD), reflecting the fact that net Caco-2 A*→*B permeability is a composite readout shaped by both passive physicochemistry and active-transport biology.

The model architecture is a GPS-style graph transformer [Rampášek et al., 2022] that combines a local message-passing component with a global all-atom pairwise attention component. The local update is performed by a DMPNN [Yang et al., 2019] operating over directed bond edges, using chemprop-style atom and edge featurizations [Heid et al., 2024] augmented with per-atom pKa values predicted by UniPKa [Luo et al., 2024] at acidic and basic sites. The global update is performed by attention over all atom pairs, with each pair represented by a learned embedding constructed from the corresponding atom and bond features, the topological graph distance between the two atoms, and the minimum and average pairwise distances across an ensemble of RDKit conformers [Landrum and RDKit Contributors, 2024]. The pairwise representation is used as an additive bias to the attention logits and is not updated across layers.

At inference, each endpoint is surfaced according to how it is used downstream. For LogD and Caco-2 permeability, we surface a continuous predicted value, since LogD and Permeability are most informative as a graded property rather than a binary developability call. For LogS, we surface a three-bin prediction with a high-precision bin for compounds predicted to be highly soluble (LogS *> −*4), a high-recall bin for compounds predicted to be insoluble (LogS *< −*6), and an uncertain middle bin for predictions falling between the two thresholds. The top bin is calibrated to identify compounds we can advance with confidence in their developability, and the bottom bin is calibrated to comprehensively screen out the least developable chemistry. The middle bin reserves uncertainty for compounds where neither call is supported.

#### 3.2.4 Retrospective Results

We benchmark each of our three models against ADMET-AI [Swanson et al., 2024], the most widely used open-source ADMET prediction package and a strong reference point for the state of ADMET prediction. ADMET-AI returns continuous predicted values for all three endpoints, while we return continuous predictions for LogD and Caco-2 A*→*B and a three-bin classification for LogS. For LogD and Caco-2 we therefore compare directly as regressors. For LogS we align the two output formats by binning ADMET-AI’s predictions by rank: for any subset of the test set, we take the top-K and bottom-K compounds by predicted value to match the number of compounds assigned to our high and low bins for that subset, and compute precision and recall against fixed ground-truth thresholds. This forces both models to commit to the same number of pass and reject calls, isolating which compounds each model selects rather than how aggressively it bins.

The test sets used here are constructed under the leakage criteria specified in Appendix D.2 and are fully out-of-distribution for our model, reflecting the most distant kind of chemistry encountered in virtual screening over commercial catalogs. They were not constructed with ADMET-AI’s training set in mind, so the same compounds are not necessarily out-of-distribution for ADMET-AI. We analyze performance degradation on in-distribution and out-of-distribution subsets of our test set, defined relative to ADMET-AI’s training data, in Appendix D.3.

#### LogD

Figure 5 shows predicted versus measured LogD for both models. On the full test set the two models perform comparably: MAE 0.67 vs 0.68, Pearson r 0.73 vs 0.73, Spearman *ρ* 0.74 vs 0.71. We interpret this as expected. LogD is a comparatively saturated task, well-covered by public data and largely additive in its physical-chemistry determinants.

**Figure 5:**
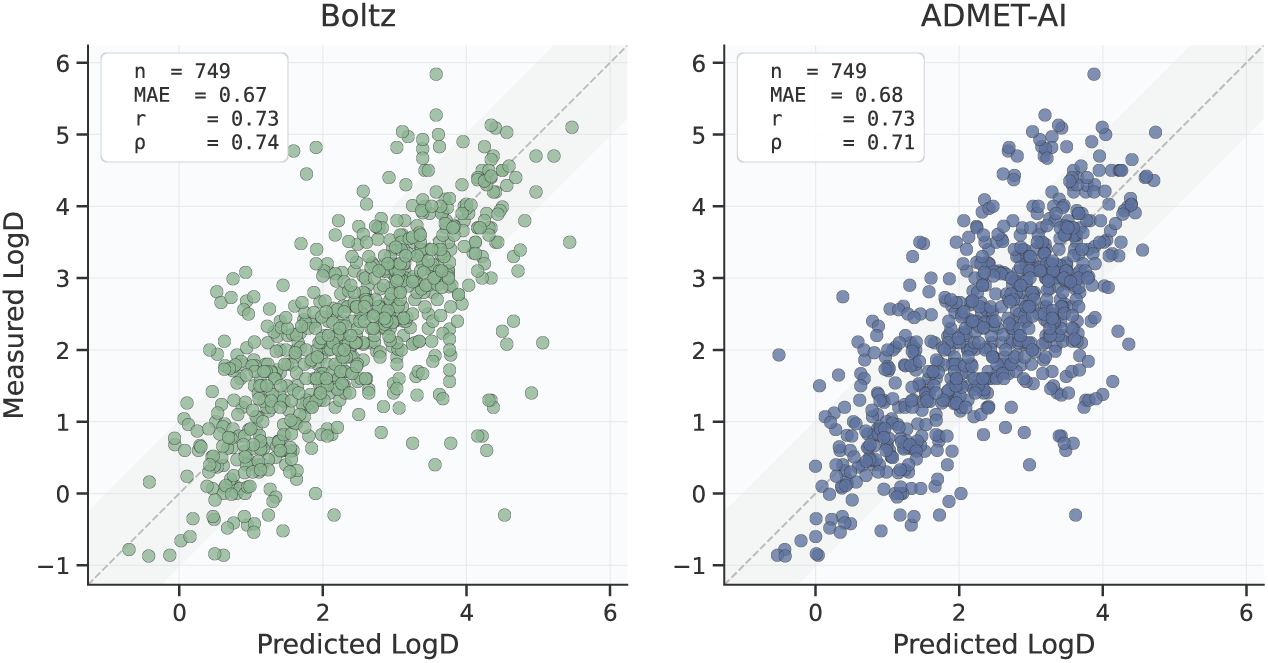
Predicted versus measured LogD for our model (left) and ADMET-AI (right). Reported metrics are MAE, Pearson r, and Spearman *ρ* across the full test set.

#### LogS

Figure 6 reports high-solubility precision and low-solubility recall for both models. Our model substantially exceeds ADMET-AI on every metric: high-solubility precision 0.74 vs 0.55 (Δ + 0.20), low-solubility recall 0.73 vs 0.50 at LogS *< −*5 (Δ + 0.22), and 0.93 vs 0.64 at the stricter LogS *< −*6 threshold (Δ + 0.29).

**Figure 6:**
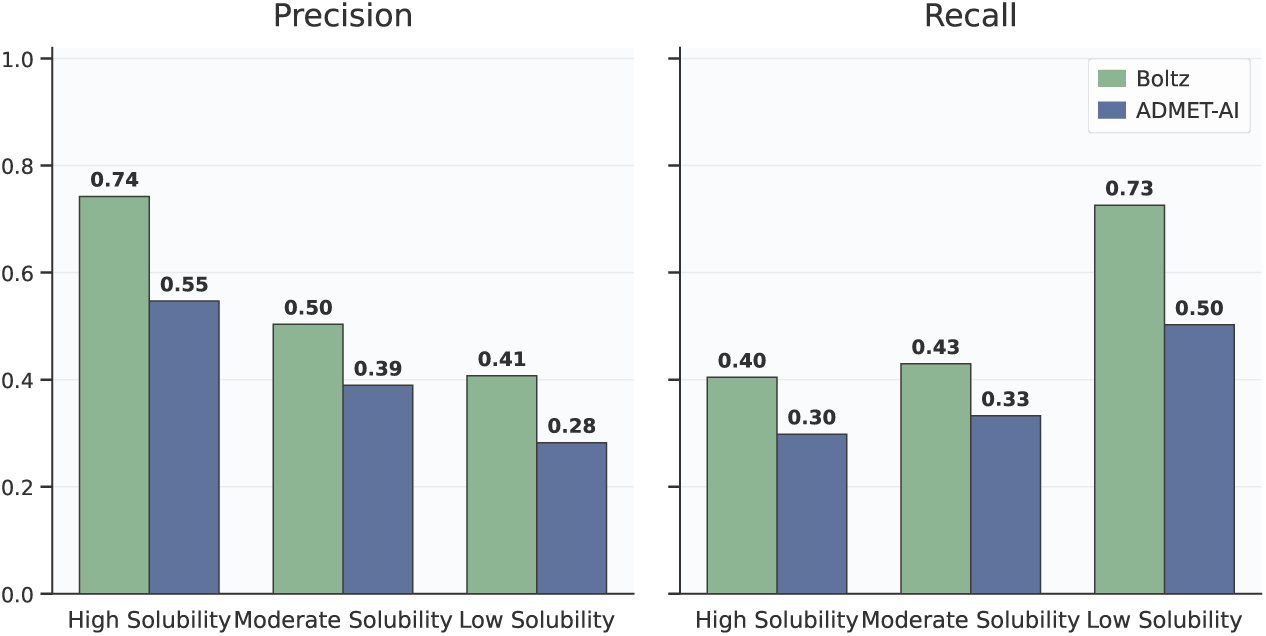
High-solubility precision and low-solubility recall for our model and ADMET-AI on the LogS test set. Truth thresholds at LogS *> −*4 for the high-solubility bin and LogS *< −*5 / *< −*6 for the low-solubility bin.

#### Caco-2 A*→*B

Figure 7 shows predicted versus measured Caco-2 A*→*B permeability for both models, coloured by measured efflux ratio. On the full test set we exceed ADMET-AI across every metric: MAE 0.46 vs 0.55, Pearson r 0.64 vs 0.48, Spearman *ρ* 0.65 vs 0.46. The gain is largest among low-efflux compounds (ER *<* 2), where passive physicochemistry dominates, while the harder regime is ER *>* 2, where accurate prediction requires modeling active efflux rather than physicochemistry alone, and absolute performance drops sharply for both models. We retain a relative advantage there, but fully capturing active efflux remains difficult, and neither model is yet at a level where it can be relied on to triage efflux-substrate chemistry without orthogonal data.

**Figure 7:**
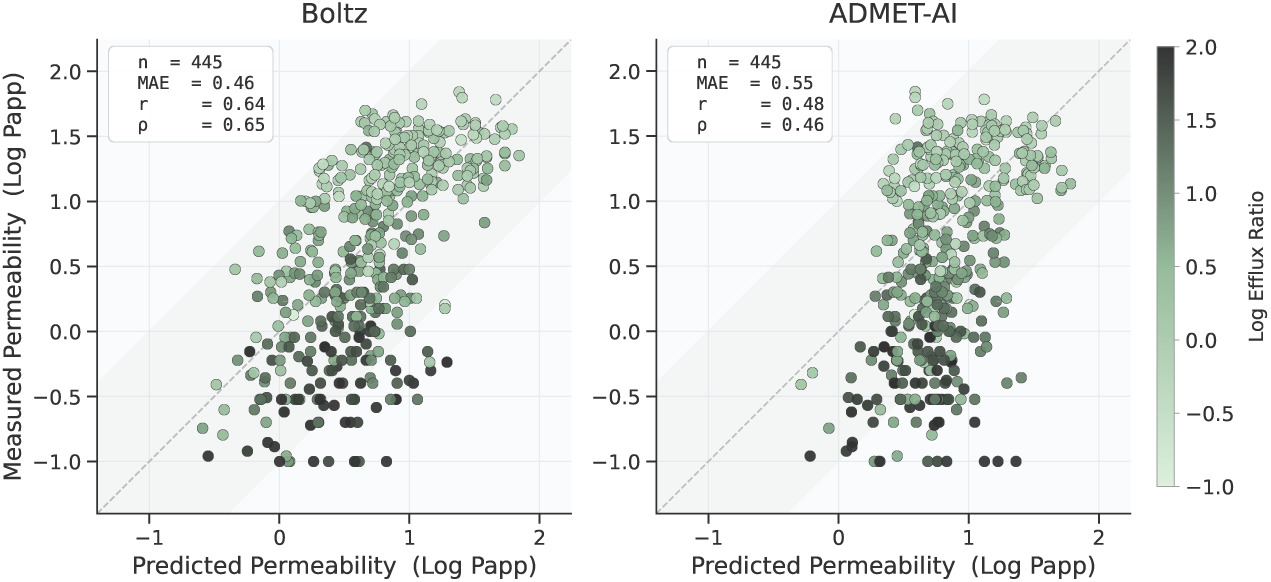
Predicted versus measured Caco-2 A*→*B permeability (log Papp) for our model (left) and ADMET-AI (right), with points coloured by log efflux ratio. Reported metrics are MAE, Pearson r, and Spearman *ρ* across the full test set.

## 4 Conclusion

We introduce a hit-finding pipeline in which affinity-model prioritization, rigorous filtering, and immediate catalog procurement together yield experimentally confirmed binders at budgets of tens rather than tens of thousands of compounds, within weeks and not months. Across ten targets spanning pseudokinases, a serine/threonine kinase, SH2-domain transcription factors, class A, B, and C GPCRs, an ion channel, a paracaspase, and ATG8-family protein–protein interaction surfaces, we observed validated activity on 60%, with most targets absent from the underlying affinity training data and several lacking any small-molecule co-crystal structure. The pipeline produced tractable starting points on systems where structure- and ligand-based approaches have historically struggled, making it well-suited to primary hit identification against validated targets, tool and probe discovery for orphan or emerging biology, generation of small-molecule equity for mechanistic and target-validation studies, and structural-biology enablement, where even modest-affinity binders can unlock co-crystallography or cryo-EM on undercharacterized systems. Retrospective ADMET analysis further indicates that early triage on solubility, lipophilicity, and permeability meaningfully reduces the developability liabilities carried into experimental testing, concentrating limited assay capacity on compounds with a more credible path forward. Together, these results define a pipeline in which model-guided prioritization, fast procurement, lightweight ADMET filtering, and iterative experimental triage combine into a hit-finding system that is broadly accessible across target classes and well-matched to the resource constraints of many discovery settings.

## Author contributions

J.K. and M.M. contributions involved activities solely related to the ATG8 targets. N.W. and A.G. contributions involved activities solely related to PknB.

## Acknowledgments

A.G. acknowledges the support of NIH award R01GM135631 and Jeanne M. Rowe. N.W. acknowledges VA Career Development Award 5IK2BX005082-03. We are grateful to Spencer S. Ericksen for assistance with the computational analysis of PknB. We thank collaborators and early platform users for feedback on target selection, assay design, and pipeline implementation. We are grateful to our collaborators at WuXi and Crelux for their experimental contributions, with special thanks to Sophie Rau and Xiaojing Guo for coordinating the experimental work.

## Declarations

- Competing interests: N.G., G.S., A.C., V.F., L.C., F.C., J.W., G.C., S.P. are affiliated with Boltz PBC. V.F. is affiliated with Boltz PBC and MIT CSAIL and his contributions to this work were performed exclusively under his Boltz PBC affiliation.
- N.W. is an inventor on U.S. patent #9540369 “Use of kinase inhibitors to increase the susceptibility of Gram+ bacteria to b-lactam antibiotics”.

## A Binding and Functional Assays

### A.1 Surface Plasmon Resonance (SPR)

SPR measures binding kinetics including association and dissociation rates to calculate the *K_D_* value, and supports diverse screening applications such as fragment and antibody screening [Patching, 2014]. The SPR assay runs on Biacore 8*K/*8*K*+.

The SPR buffer with/without DMSO were prepared and filtered by 0.22 *µ*m filtration unit (bottle top filter; Nalgene) equipped with a 0.22 *µ*m filter, check the pH of the buffer and adjust it to the indicated value with NaOH or HCl. Wash the instrument by taking the chip out of the fridge and let it adjust to room temperature, dock the chip then immediately change solution to SPR buffer without DMSO. Thaw protein aliquot on ice and dilute to the indicated final concentration in SPR buffer without DMSO. Dilute the protein to the indicated concentration in SPR immobilization buffer without DMSO. To capture the biotinylated target protein onto a SA Chip, dispense the pre-conditioning solution, wash solution and diluted protein to the respective wells of a 96 well SPR plate, then cover the plate with microplate foil. The immobilization has to be carried out at 20 C Flow cell and sample compartment temperature. After the immobilization, prepare the upper and lower solvent correction solutions by adding the indicated volume of 100% DMSO to the DMSO-free SPR buffer, vortex vigorously, prepare the other six points of solvent correction by mixing different ratios of the upper and lower solvent correction solutions (referring to “8point solvent correction”), vortex vigorously and distribute the dilutions into a 96-well deepwell plate; next, dilute compounds with DMSO in an Eppendorf tube to reach a working concentration, mix by vortexing, add 20 *µ*L of the diluted compounds to well 1 of the SPR plate, distribute 10 *µ*L (using the Multipette E3, Eppendorf) into wells 2-9 of the SPR plate, perform serial dilutions using the Integra Assist pipette (12.5 *µ*L) by mixing well 1 of the SPR plate for 15 mixing cycles at 12.5 *µ*L volume, taking 10 *µ*L compound from well 1 and transferring it to well 2, mixing well 2 for 15 cycles at 12.5 *µ*L volume, continuing in the same manner up to well 9 and discarding the last 10 *µ*L after mixing well 9, distribute 98 *µ*L SPR immobilization buffer (without DMSO) into an SPR plate using a Multipette E3 (Eppendorf, yield 1 mL per well), transfer 2 *µ*L of the serial dilution to the SPR plate containing 98 *µ*L buffer using the Cybi Selma and mix 50 times with 25 *µ*L volume, seal the SPR plate with microplate foil, shake the plate at 800 rpm for 5 minutes, and finally centrifuge the plate for 2 minutes. Then the plate was loaded to Biacore 8*K/*8*K*+ for measurement.

### A.2 Spectral Shift (SpS) and Microscale Thermophoresis (MST)

Spectral Shift technology uses a dual-emission optical system, which detects fluorescence emission simultaneously at two defined wavelengths (650 nm and 670 nm). The target molecule is labeled with an environment-sensitive near infrared dye, which experiences a subtle shift of the emission spectrum upon changes in its microenvironment under isothermal conditions by 2 distinct mechanisms. Either by binding of a ligand in close proximity to the position of the fluorophore or by inducing a conformational change in the target molecule [Langer et al., 2022].

Microscale Thermophoresis is also measured in capillaries. The fluorescence within the capillary is excited and detected through the same objective. A focused IR-Laser is used to locally heat a defined sample volume. Thermophoresis of fluorescent molecules through the temperature gradient is detected [Jerabek-Willemsen et al., 2011]. Both SpS and MST assay are run on Monolith.

In a PCR tube dilute the compounds to working concentration with DMSO. Use 2 separate Corning Microplates as ligand plate and Assay plate. Making a 1:3 dilution series by transferring 15 *µ*l of compound at working concentration into well 1 of the ligand plate. With the Eppendorf Multipette (+ Combitip 0.2 ml) distributing 11x 10 *µ*l DMSO into wells 2-12 of the ligand plate, using the Integra VIAFLO pipette (12.5 *µ*l) to do serial dilutions: take 5 *µ*l tool peptide from well 1, transfer to well 2 and mix (12 mixing cycles, 12.5 *µ*l volume). Continue in the same manner up to well 12. Discard the last 5 *µ*l after mixing of well 12. In a reaction tube prepare 550 *µ*l of 20 nM labeled protein (Use freshly labeled protein or protein from the same day) per interaction in duplicate. With the Eppendorf Multipette (+ Combitip 0.5 ml) dispense 19.5 *µ*l of protein (20 nM protein) into wells 1-24 of the Assay plate. With the Integra VIAFLO pipette (12.5 *µ*l) add 0.5 *µ*l of tool peptide from the dilution series (ligand plate) to the corresponding wells 1-12 and 13-24 of the Assay plate (to obtain duplicates) and mix well (12 mixing cycles, mixing volume 12.5 *µ*l). This results in a final DMSO concentration of 2.5%. Ensure that all 12 tips are attached to the multi-channel pipette with equal force by gentle rocking the pipette back and forth and that all tips have aspirated the same volume. Incubate the Assay plate at RT for 30 min. Fill the 2 x 12 concentrations into Monolith^TM^ NT.115 Monolith Premium Capillary Chips by dipping the chips into the solutions. Insert the chip into the instrument and fix it by using the Monolith Premium Capillary Chip Lid. Start the measurement with the following measurement parameters: LED power, auto-excitation (device sets fluorescence to 1000-1200 counts). Spectral Shift and MST at the same time. MST power, high. MST Settings, 3 s - 20 s - 1 s (initial fluorescence - MST on time - back-diffusion). Analysis Mode: Spectral shift (ratio), MST (670 nm) after 5 s.

### A.3 Nano Differential Scanning Fluorimetry (NanoDSF)

NanoDSF measures protein thermal stability by monitoring intrinsic tryptophan/tyrosine fluorescence shifts during heating, determining Tm (melting temperature). It screens optimal buffer/ligand conditions, analyzes molecular interactions, and assesses protein purity/aggregation [Magnusson et al., 2019]. The signal was measured by Prometheus NT.48.

Thaw the target protein on ice and keep it on ice thereafter, centrifuge the protein for 10 min at 15000 rpm and 4 C, carefully remove the supernatant and pipette into a fresh tube on ice, then measure protein concentration using UV. Prepare single use aliquots of the tool compound by diluting the compounds to working concentrations in PCR tubes using DMSO. Then prepare 6 PCR tubes with total volume: 2 *×* 30 *µ*l DMSO control (quadruplicates), 1 *×* 30 *µ*l per tool compound (duplicates) and 1 *×* 30 *µ*l per analyte compound (duplicates), with final DMSO concentration 2%. Use the Integra Pipette (125 *µ*l) for mixing (12 mixing cycles, 18 *µ*l mixing volume). Spin tubes down briefly, ensure samples are air bubble free, fill the capillaries by carefully sampling (fill completely and air bubble free, fill two capillaries with the same dip for duplicates), place the capillaries centered on the tray and attach the magnetic lid to fix the capillaries, close the drawer, then start the measurement with instrument settings: LED power e.g. 30% (fluorescence should lie between 1000 and 20000), start temperature 20 C, end temperature 95 C, temperature ramp 2 C*/*min.

### A.4 Patch Clamp

hNav1.8/*β*1 sodium channel activity was evaluated using automated whole-cell patch clamp on a SyncroPatch 384PE (Nanion) at room temperature [Obergrussberger et al., 2016]. HEK293 cells stably expressing hNav1.8/*β*1 (WuXi AppTec) were maintained at 37 C, 5% CO2 in DMEM supplemented with 11% FBS, G418, blasticidin, zeocin and GlutaMAX (see Table 1 for vendor/catalog details). Cells *≥* 2 days post-plating and *>* 75% confluent were harvested with TrypLE and resuspended in physiological solution prior to recording. Solutions were (mM): physiological/external – NaCl 140, KCl 4, CaCl2 2-3, MgCl2 1, glucose 5, HEPES 10, pH 7.4 (298 mOsm); internal – NaCl 10, CsF 110, CsCl 10, EGTA 10, HEPES 10, pH 7.2 (280 mOsm). Assay setup, chip priming, cell catch/seal, amplifier settings and voltage/application protocols were configured using Biomek/PatchControl software. After baseline recording following addition of 40 *µ*L vehicle (300 s), test compounds (final concentration 1.00 *µ*M; 40 *µ*L addition) were applied with exposures *≥* 300 s per concentration. VX548 was included as a positive control. A minimum of three technical replicates were obtained per compound; wells failing automated QC were excluded and retested. Data were analyzed to determine percent inhibition relative to vehicle and positive control and are reported as mean *±* SD.

### A.5 cAMP

cAMP kits are based on a competitive format involving a specific antibody labeled with cryptate (donor) and cAMP coupled to d2 (acceptor) [Hill et al., 2017]. Luminescence was measured by Perkin Elmer EnVision. Dose-response curves for GLP-2R were fit as described in Appendix A.7.

#### Transfected Cell Line

CHO-K1 cells transfected with Amylin3 and GLP-2 were grown to 80%-90% confluence in F-12 culture media (10% FBS and 1X penicillin/streptomycin). Cells were detached with 0.25% trypsin/EDTA and resuspended with plating media. Centrifuge and resuspend cells with assay buffer (DPBS, 0.5 mM IBMX, 0.1% Casein, 20 mM HEPES) to the optimized cell density (100 K cells/mL). Dispense 10 *µ*L (1 K cells/well) of cells into assay plate, which contains 10 nL of serial diluted compound stock each well. Spin for 10 seconds at 1000 rpm and incubate at 37 C for 30 minutes. Dilute donor and acceptor reagents using lysis detection buffer included in the kit. Dispense 5 *µ*L of diluted cAMP-d2 and 5 *µ*L of diluted Anti-cAMP-Cryptate with Combi multi-drop and incubate at RT for 30 minutes. Read plates at 665/615 and calculate results.

#### Parental Cell Line

CHO-K1 parental cells were grown to 80%-90% confluence in F-12 culture media (10% FBS and 1X penicillin/streptomycin). Cells were detached with 0.25% trypsin/EDTA and resuspended with plating media. Centrifuge and resuspend cells with assay buffer (DPBS, 0.5 mM IBMX, 0.1% Casein, 20 mM HEPES) to the optimized cell density (100 K cells/mL). Dispense 10 *µ*L (1 K cells/well) of cells into assay plate, which contains 10 nL of serial diluted compound stock each well. Spin for 10 seconds at 1000 rpm and incubate at 37 C for 30 minutes. Dilute donor and acceptor reagents using lysis detection buffer included in the kit. Dispense 5 *µ*L of diluted cAMP-d2 and 5 *µ*L of diluted Anti-cAMP-Cryptate with Combi multi-drop and incubate at RT for 30 minutes. Read plates at 665/615 and calculate results.

### A.6 FLIPR

Calcium mobilization was measured by FLIPR (Fluorometric Imaging Plate Reader) Penta with the calcium flux released from cells upon stimulation and the calcium-specific indicator fluo-8 according to manufacturer’s protocol [Schroeder and Neagle, 1996]. Dose-response curves for MRGPRX2 were fit as described in Appendix A.7.

#### Transfected Cell Line

HEK 293 cells transfected with mGlu4 and MRGPRX2 were grown to 80%-90% confluence in DMEM culture media containing 10% FBS and 1X penicillin/streptomycin. Cells were detached with 0.05% trypsin/EDTA and resuspended with plating media. Cells were seeded at 30 *µ*L/well in 384-well (20 K cells/well) clear-bottomed poly-D-lysine coated black tissue culture plates and incubated overnight at 37 C, 5% CO2.

On the day of the assay, cells were loaded with 10 *µ*L/well of 4X loading dye solution and incubated at 37 C, 5% CO2 for 30 minutes. Compound dilutions were prepared from 10 mM dimethylsulfoxide stocks. Compound additions (10 *µ*L of 5X) were done on the FLIPR Penta and changes in fluorescence that reflect calcium mobilization were monitored at 1-second intervals for 130 seconds (excitation wavelength=470-495 nm, emission wavelength=515-575 nm). Data were exported as the difference between maximum and minimum fluorescence observed for each well.

#### Parental Cell Line

HEK 293 parental cells were grown to 80%-90% confluence in DMEM culture media containing 10% FBS and 1X penicillin/streptomycin. Cells were detached with 0.05% trypsin/EDTA and resuspended with plating media. Cells were seeded at 30 *µ*L/well in 384-well (20 K cells/well) clear-bottomed poly-D-lysine coated black tissue culture plates and incubated overnight at 37 C, 5% CO2.

On the day of the assay, cells were loaded with 10 *µ*L/well of 4X loading dye solution and incubated at 37 C, 5% CO2 for 30 minutes. Compound dilutions were prepared from 10 mM dimethylsulfoxide stocks. Compound additions (10 *µ*L of 5X) were done on the FLIPR Penta and changes in fluorescence that reflect calcium mobilization were monitored at 1-second intervals for 130 seconds (excitation wavelength=470-495 nm, emission wavelength=515-575 nm). Data were exported as the difference between maximum and minimum fluorescence observed for each well.

### A.7 Dose-Response Curve Fitting

Concentration-response data from the cAMP (Appendix A.5) and FLIPR (Appendix A.6) assays were analysed with the drc package (version 3.0-1) [Ritz et al., 2015] in R 4.6. Each compound was tested in duplicate across a ten-point, three-fold serial dilution from a top concentration of 50 *µ*M; reference ligands (Teduglutide for GLP-2R; Cortistatin 14 and the reference antagonist for MRGPRX2) were titrated from their respective top concentrations. Per-well responses were normalized to percent activity (agonist mode; 0% = vehicle, 100% = the EC_100_ response of the reference agonist) or percent inhibition (antagonist mode; relative to the reference agonist challenge) and fit against compound concentration with a four-parameter log-logistic model (LL.4),

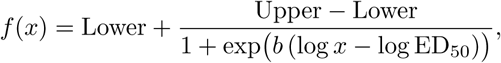

equivalent to the standard four-parameter logistic (Hill) equation with Hill slope = *−b* and midpoint potency EC_50_/IC_50_ = ED_50_. Potencies are reported as EC_50_/IC_50_ and as pEC_50_*/*pIC_50_ = *—* log_10_(ED_50_ [M]) with 95% confidence intervals from the standard error of log ED_50_; E_max_ (agonist) and maximal inhibition (antagonist) are taken as the fitted upper asymptote, expressed relative to the reference EC_100_.

### A.8 KinaseGlo

Kinase inhibition was measured using the KinaseGlo luminescence-based assay (Promega), which quantifies residual ATP remaining after the kinase reaction as a proxy for enzymatic activity. Assays were performed in 96-well white, flat-bottom plates in a total reaction volume of 50 µL.

PknB 1–331 expressed as a GST-fusion was purified and the tag was cleaved, and then was diluted to 0.25 µM in assay buffer (10 mM Tris-HCl pH 7.4, 150 mM NaCl, 1 mM DTT, 1 mM MgCl_2_). Compounds were first screened in single points in triplicate at a final concentration of 512 *µM*. The top 18 compounds were then selected for dose-response analysis. Compound stocks were prepared at 25.6 mM in DMSO and serially diluted 1:2 in DMSO across eight points. Compounds were transferred to assay plates to yield final concentrations ranging from 512 *µM* to 4 *µM* at a final DMSO concentration of 2%. GSK690693 was included on each plate as a reference inhibitor at a top concentration of 16 *µM*. Uninhibited enzyme controls (DMSO vehicle only) and no-enzyme background controls were included in each run. Compounds and enzyme were pre-incubated for 10 minutes at room temperature prior to substrate addition.

Kinase reactions were initiated by addition of a substrate mix containing ATP (final concentration 100 µM) and GarA full-length (final concentration 40 µM). Plates were incubated at 37°C for 20 minutes, then equilibrated to room temperature for 10 minutes. An ATP standard curve (100%, 50%, 25%, 12.5%, 6.25%, 3.125%, 1.5625%, and 0% of the reaction ATP concentration) was included on each plate to confirm assay linearity and enzyme loading.

Following the kinase reaction, 25 µL of KinaseGlo reagent was added to each well and plates were incubated at room temperature for 15 minutes in the dark. Luminescence was measured with an integration time of 1 second per well. Raw luminescence units were normalized to the uninhibited enzyme control (0% inhibition) and no-enzyme background (100% inhibition). IC_50_ values were determined by nonlinear regression using GraphPad Prism 10, fitted to a four-parameter logistic model. *K_i_* values were derived from IC_50_ values using the Cheng-Prusoff equation,

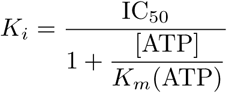

where *K_m_*(ATP) for PknB was taken as 42 µM.

### A.9 AlphaScreen for LC3B and GABARAP

Purified LC3B or GABARAP was diluted in assay buffer (25 mM HEPES, 150 mM NaCl, 0.1% Tween-20, 1 mg/mL BSA pH 7.3) to 200 nM and 100 nM (LC3B) or 100 nM and 50 nM (GABARAP) [Eglen et al., 2008]. 2.5 µL of the more concentrated protein (200 nM LC3B or 100 nM GABARAP) was added in duplicate to a white, 384-well polystyrene plate (AlphaPlate, Revvity) to wells intended for inhibitor compounds. 5 µL dilute protein (100 nM LC3B or 50 nM GABARAP) was added to control wells. Inhibitor compounds were dissolved at 20 mM in DMSO. From the 20 mM stock, 500 µL solutions of each compound were prepared in assay buffer with 7.5% DMSO in an Eppendorf tube. The compound solutions were serially diluted 1:1 in assay buffer with 7.5% DMSO. 15 µL of each serially diluted compound was added to wells with 2.5 µL of concentrated protein. 5 µL of non-biotinylated K1 or non-biotinylated FYCO1S were added to wells with 5 µL dilute protein, acting as positive control inhibitors for GABARAP and LC3B, respectively [Olsvik et al., 2015]. Plates were covered in foil, spun down at 1200 rpm for 3 minutes, and then incubated at room temperature for 45 minutes. Biotinylated K1 was diluted to 100 nM (concentrated) and 50 nM (dilute) in assay buffer, or biotinylated FYCO1S was diluted to 150 nM (concentrated) and 75 nM (dilute) in assay buffer. 5 µL of dilute biotinylated K1 or biotinylated FYCO1S was added to control wells with GABARAP or LC3B, respectively. 2.5 µL of concentrated biotinylated K1 or biotinylated FYCO1S was added to wells with inhibitor compounds and GABARAP or LC3B, respectively. Three blank wells containing 10 µL assay buffer and 5 µL assay buffer with 7.5% DMSO, as well as three “no-inhibitor” control wells containing 5 µL dilute protein, 5 µL dilute biotinylated peptide, and 5 µL assay buffer with 7.5% DMSO were included in each plate. Plates were covered in foil, spun down, and incubated for 45 minutes. AlphaScreen streptavidin donor beads and nickel chelate acceptor beads (Revvity) were each diluted to 200 mg/mL (concentrated) and 100 mg/mL (dilute) in assay buffer. 5 µL dilute acceptor beads were added to each control well, and 2.5 µL concentrated acceptor beads were added to each inhibitor compound well. In the dark, 5 µL dilute donor beads were added to each control well, and 2.5 µL concentrated donor beads were added to each inhibitor compound well. Plates were covered with foil, spun down, and incubated for 1 hour before being read on a plate reader (Tecan Spark, AlphaScreen method, excitation at 680 nm and emission at 520-620 nm). Using GraphPad Prism 10 software, data was normalized using blank wells as 0% bound and no-inhibitor control wells as 100% bound, then fit with IC_50_ curves. Average IC_50_ values and standard errors of the mean were calculated based on at least three independent trials.

## B ADMET Assays

### B.1 logD

Log D (pH 7.4) was determined using a miniaturized 1-octanol/buffer shake-flask method with LC-MS/MS semi-quantification [Andrés et al., 2015]. Test compounds and QC standards were prepared as 10 mM solutions in DMSO and plated (2 *µ*L/well) into 96-well plates. To each well, buffer-saturated 1-octanol (149 *µ*L) and octanol-saturated 0.1 M phosphate buffer (pH 7.4; 149 *µ*L) were added to give a nominal compound concentration of 66.7 *µ*M. Samples were vortexed 2 min and shaken at 800 rpm at room temperature for 1 h, then centrifuged (4,000 rpm, 5 min) to effect phase separation. Aliquots of the octanol and aqueous layers were diluted with internal standard solution (dilution factors chosen per compound properties) and analyzed by LC-MS/MS; concentrations were assessed semi-quantitatively as analyte/IS peak area ratios. LogD_7.4_ was calculated as

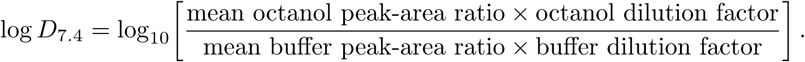

QC compounds (testosterone, propranolol HCl and nadolol) were assayed to monitor performance.

### B.2 Kinetic Solubility

Kinetic aqueous solubility was measured by a miniaturized shake-flask method with HPLC-UV quantification [Lin and Pease, 2016]. Test compounds were prepared as 10 mM stocks in DMSO and dosed (10 *µ*L) into 490 *µ*L of phosphate buffer (50 mM, pH 7.4) in MiniUniPrep vials or 96-well plates to give a nominal concentration of 200 *µ*M. Samples were vortexed 2 min, shaken at 800 rpm at room temperature for 24 h, then centrifuged (4,000 rpm, 25 C, 10 min) and filtered to remove precipitate. Aliquots (200-400 *µ*L, diluted as necessary) of the supernatant were analyzed by HPLC-UV; each run included at least five calibration standards and QC compounds (celecoxib, carbamazepine, chloramphenicol). Kinetic solubility (*µ*M) was calculated as measured concentration *×* dilution factor and reported as *µ*M and *µ*g/mL.

### B.3 Bidirectional permeability in Caco-2 Cells

#### Without inhibitors

Bidirectional permeability of test compounds was assessed in Caco-2 cell monolayers cultured on 96-well permeable inserts (ATCC) for 21-28 days [Hubatsch et al., 2007]. Monolayer integrity was confirmed by Lucifer yellow exclusion and by measuring unidirectional permeability of nadolol (low permeability) and metoprolol (high permeability) and bidirectional transport of digoxin (Pgp substrate) in duplicate. Transport assays were performed in HBSS supplemented with 10 mM HEPES (pH 7.40 *±* 0.05) at 37.0 C, 5% CO2. Test compounds (2.00 *µ*M, DMSO *≤* 1.0%) were applied in both apical *→* basolateral (A *→* B) and basolateral *→* apical (B *→* A) directions (n = 2) and incubated for a single 2 h time point. A T0 sample was prepared by spiking dosing solution into transport buffer and mixing with stop solution containing an internal standard (IS). At 2 h, donor and receiver samples were collected and immediately mixed with stop solution; all samples (including T0) were analyzed by LC-MS/MS and quantified semi-quantitatively as analyte/IS peak area ratios (no calibration curve).

#### With inhibitors

Bidirectional and efflux-inhibition permeability were assessed using Caco-2 monolayers cultured on 96-well permeable inserts (ATCC) for 21-28 days. Monolayer integrity was confirmed by Lucifer yellow exclusion and by measuring unidirectional permeability of nadolol (low permeability) and metoprolol (high permeability) and bidirectional transport of digoxin (Pgp substrate) in duplicate. Transport assays were performed in HBSS with 10 mM HEPES (pH 7.40 *±* 0.05) at 37.0 C, 5% CO2. Test compounds (2.00 *µ*M, DMSO *≤* 1.0%) were applied in both apical *→* basolateral (A*→*B) and basolateral *→* apical (B*→*A) directions (n = 2) for a single 2 h time point, tested both without and with the efflux inhibitor GF120918 [Hyafil et al., 1993] (included in donor and receiver wells at a validated concentration). A T0 sample was prepared by spiking dosing solution into transport buffer and mixing with stop solution containing an internal standard (IS). At 2 h, donor and receiver samples were collected and immediately mixed with stop solution; all samples including T0 were analyzed by LC-MS/MS and quantified semi-quantitatively as analyte/IS peak area ratios (no external calibration curve).

## C Experimental Notes

### C.1 STAT6

Compounds were screened utilizing ready-to-go WuXi (Crelux) biophysical screening assays: MST/SpS and SPR as described in Appendix A.

Orthogonal screening was performed against STAT6 in the aforementioned biophysical assays of 37 compounds from the WuXi OTS collection alongside established STAT6 binders - AK-068, YM-341619 and AS1517499 [McMurray, 2017] - as positive controls. SpS identified a number of putative binders, including 4 compounds with *K_D_*s between 30 and 87 *µ*M. One putative binder SM-GMUQEHEA was identified with an SpS-measured *K_D_* = 30.3 *µ*M (Fig 8). This compound showed no measurable affinity for p38*α* indicating direct STAT6 engagement. Orthogonal confirmation via SPR for STAT6 has remained challenging, due to slow compound dissociation rates. For this reason, a *K_D_* could not be calculated via SPR. Data is presented ‘as is’, with confirmatory studies ongoing to further confirm these findings.

**Figure 8:**
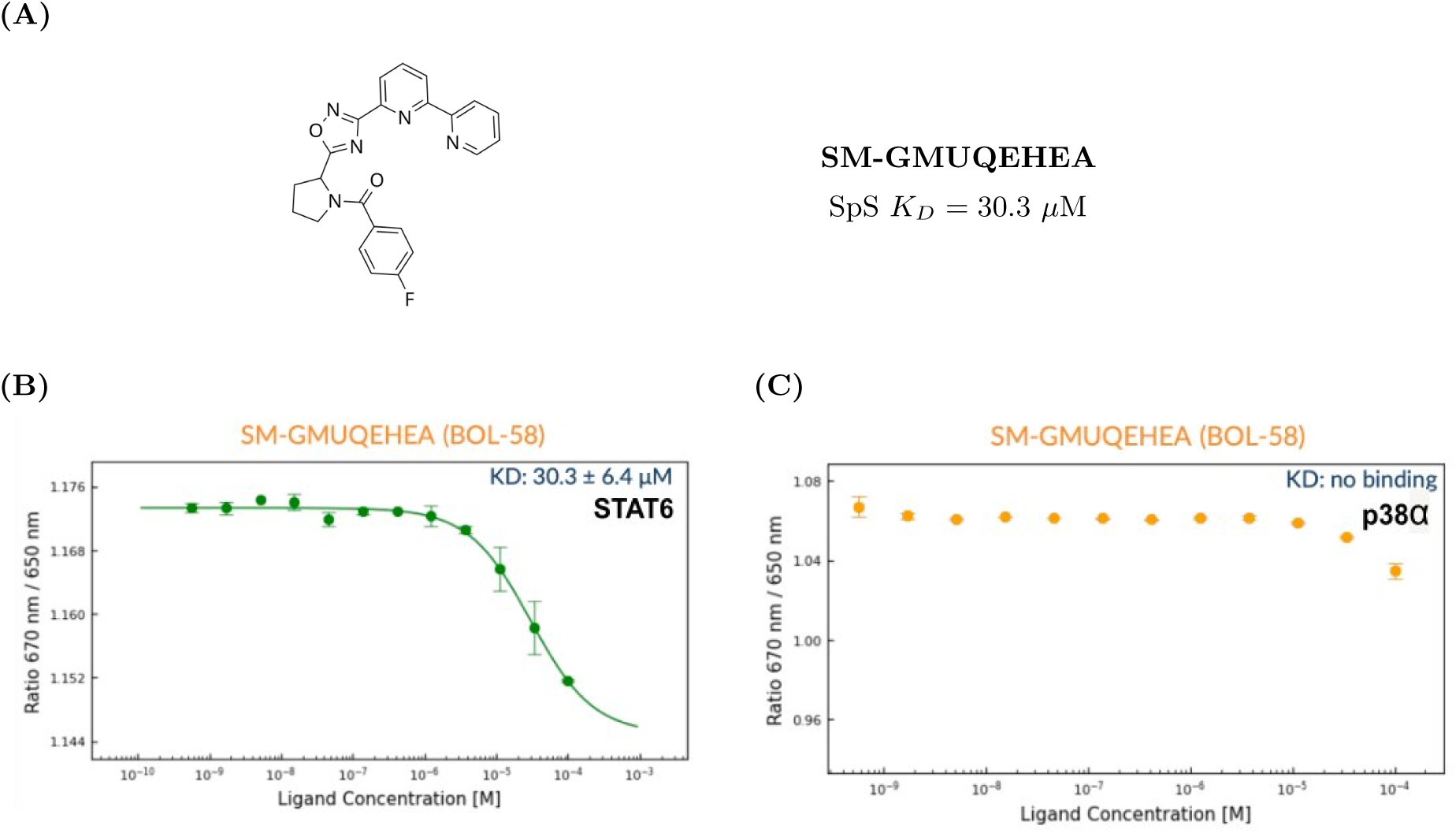
(A) Putative STAT6 binder SM-GMUQEHEA. (B) Binding affinity of SM-GMUQEHEA to STAT6 as measured by spectral shift assay. (C) Binding affinity of SM-GMUQEHEA to p38*α* as measured by spectral shift assay, which showed no binding. Spectral shift measurements were performed on the Monolith^TM^ instrument (NanoTemper).

The WuXi OTS collection was searched for structural analogues of SM-GMUQEHEA using Morgan fingerprint similarity (radius = 2, 2048 bits) [Morgan, 1965] and four readily available compounds spanning a Tanimoto similarity range of 0.43–0.66 relative to the parent were identified for confirmatory SAR analysis. SpS analysis showed that three of the four analogues behaved as weak binders; however, none of the SpS curves reached saturation, precluding accurate *K_D_* determination. This is consistent with variability observed in repeat SpS analysis of the parent compound SM-GMUQEHEA, which was run concurrently with the analogue screen. In the original SpS experiment, SM-GMUQEHEA yielded a *K_D_* of 30.3 µM across 12 concentration points; in the repeat experiment, conducted with 8 concentration points, SM-GMUQEHEA was again detected as a weak binder but the curve similarly failed to saturate, and no reliable KD could be extracted.

Retrospective analysis of co-folding predictions of analogues predicted SM-J13C2EF8 and SM-Y9H7E2LH to bind more strongly compared to parent SM-GMUQEHEA based on their Boltz Optimization Scores (continuous score between 0 and 1, with 1 being the strongest binder) while SM-C66YZ25D and SM-NHPXU2LR were predicted lower. For SM-Y9H7E2LH, this ligand was mispositioned and predicted to engage a distinct binding pocket relative to the parent and analogues. SM-J13C2EF8 and SM-C66YZ25D were predicted to adopt the opposite stereochemical orientation to SM-GMUQEHEA, such that the Cbz benzyl group of SM-J13C2EF8 and SM-C66YZ25D occupies a different region of the pocket than the 4-fluorophenyl group of SM-GMUQEHEA, which occupies a hydrophobic subpocket in the original co-folding prediction. Like the parent compound SM-GMUQEHEA, all analogues contain an oxadiazole moiety flanked by a 2,2’-bipyridyl group. The predictions highlight ligand placement proximal to an Arg residue within reach to form productive stabilising interactions; however, for SM-C66YZ25D the increase in piperidine ring size from a five- to six-membered ring relative to SM-J13C2EF8 causes a shift in ligand placement, perturbing the orientation of the central oxadiazole ring and 2,2’-bipyridyl relative to the Arg residue, which may contribute to the lack of binding observed for Z25D. Substitution of the cyclic pyrrolidine amide in SM-GMUQEHEA for an N-methylpiperazine in SM-NHPXU2LR, linked via a methylene, extends this vector toward an Asp, while the remainder of the ligand positioning is also altered but not displaced from the parent binding region.

Notably, none of the molecules described here were predicted to extend into the classical SH2 phosphotyrosine-binding subpocket - defined by the conserved Arg562, Lys480, and Ser/Thr residues and engaged by canonical phosphate groups or phosphate mimics in previously reported peptidomimetic inhibitors of the STAT6 SH2 domain - presenting a clear opportunity to grow the series into this region and improve binding affinity (Fig 9).

**Figure 9:**
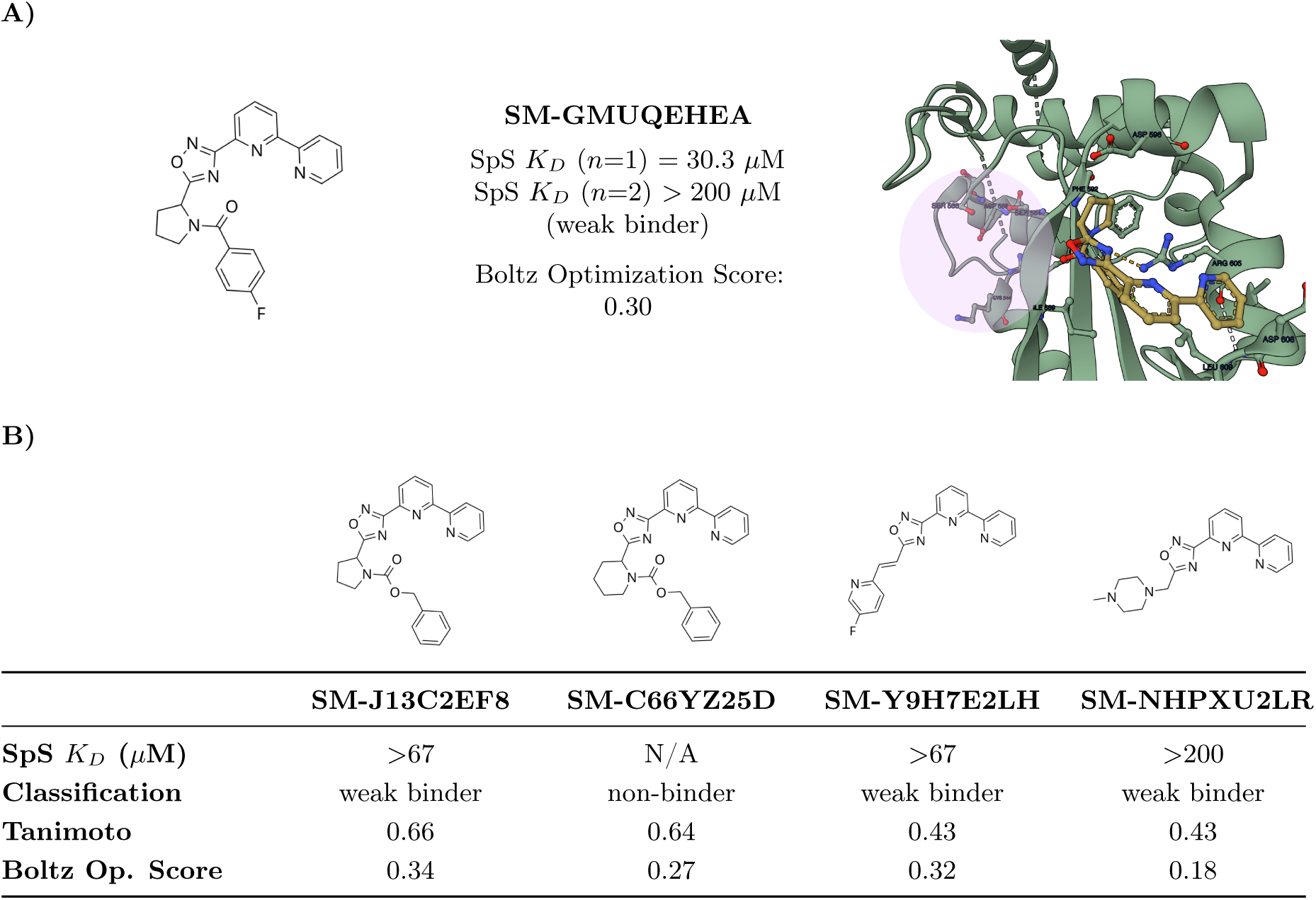
Analogue exploration of SM-GMUQEHEA. (A) Parent compound: 2D structure, SpS *K_D_* binding data, and the predicted protein-ligand binding pose. (B) Four analogues from the scaffold series with SpS *K_D_* values, classification, Tanimoto similarity to the parent, and Boltz Optimization Scores.

### C.2 GLP-2R

Compounds were screened in the cAMP accumulation assay using CHO-K1 cells stably expressing GLP-2R, as described in Appendix A. Primary screening was conducted at three concentrations (0.5, 5, and 50 *µ*M), followed by progression of compounds which gave an initial signal to full dose-response curves for IC_50_ determination. Counter-screening against the parental CHO-K1 cell line was performed in parallel to identify and exclude compounds with non-specific or cell-intrinsic activity.

Fifty-one compounds from the WuXi (OTS) collection predicted to bind the orthosteric GLP-2 peptide pocket were screened against GLP-2R to evaluate agonist and antagonist activity. Primary screening followed by DRC identified twelve compounds demonstrating functional activity in the cAMP assay, all of which displayed antagonist activity. The antagonists identified showed IC_50_ values in the double digit nanomolar to double digit micromolar range (0.07 - 24 *µ*M), with SM-4MPGPRSA as a representative example (Fig 10). Counter-screening in parental CHO-K1 cells confirmed no functional background activity. Six selected compounds from this functionally active group were carried forward for analogue exploration. Structural analogues were retrieved from the WuXi OTS collection using Morgan fingerprint similarity (radius = 2, 2048 bits) and screened in the cAMP GLP-2R assay and parental counter-screen for confirmatory SAR analysis.

**Figure 10:**
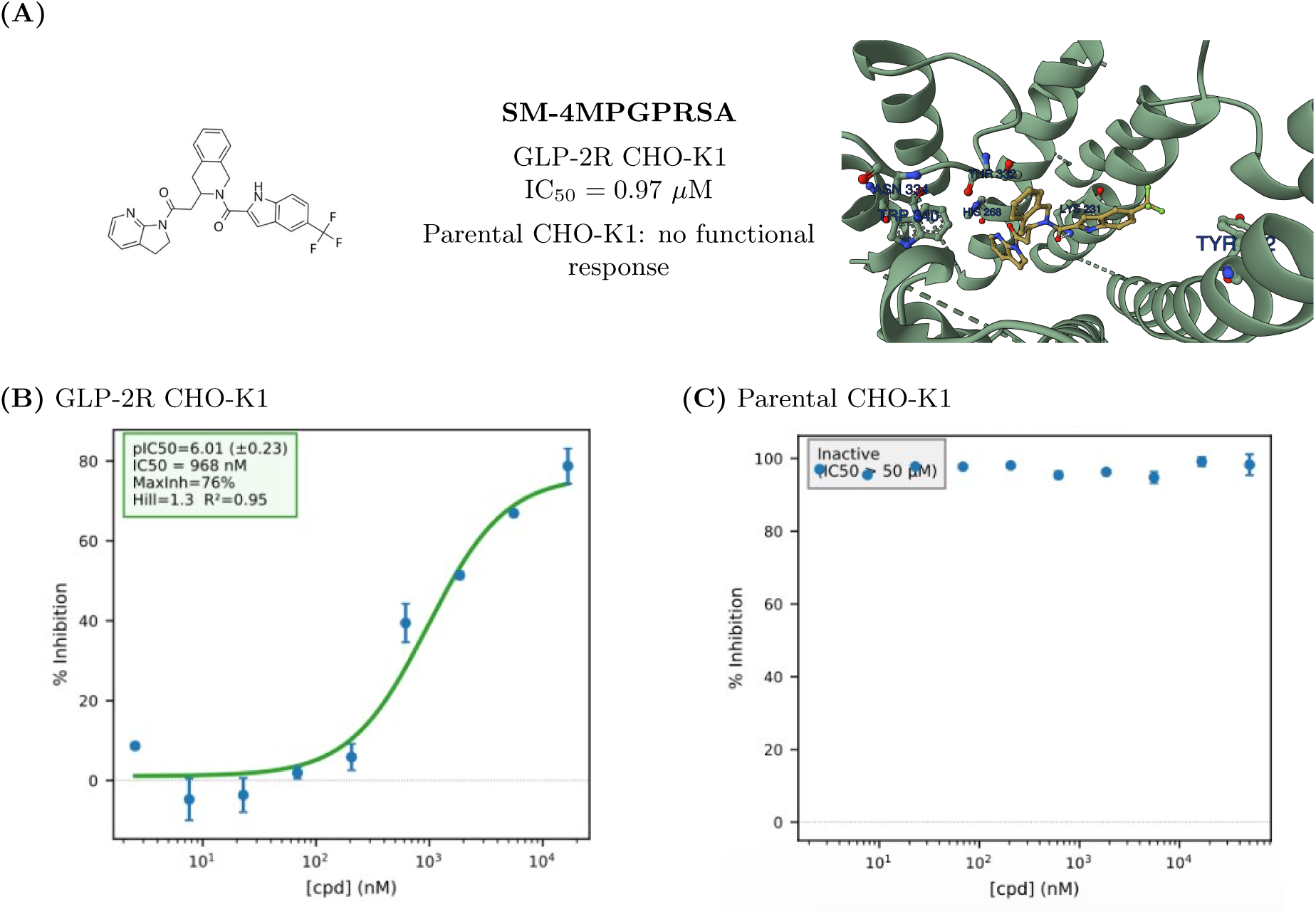
(A) Antagonist binder SM-4MPGPRSA and predicted ligand-protein binding pose placing ligand in the orthosteric GLP-2 peptide pocket. Dose-response curves for SM-4MPGPRSA in (B) GLP-2R-expressing CHO-K1 cells and (C) parental CHO-K1 cells, measured by cAMP accumulation assay.

The results of the antagonist analogue exploration are summarized in Table 2. Five of six parents yielded at least one active analogue. The most productive series was SM-WK86BLMN giving 4 actives, with the parent molecule itself remaining the most potent compound. Two further series began from sub-micromolar parents: SM-4MPGPRSA where 2/4 active analogues retained low-micromolar potency, and SM-SFKXATXC gave 2/5 active analogues in the micromolar range. Among the less polar parents, SM-EX12JGV5 was the most productive (2/5 actives) while SM-T6ABU679 each yielded one active analogue. No pharmacological switches were observed for any analogues. We noted that some parents and their analogues exhibited high Hill slopes (>2) in the cAMP assay DRCs, which is under further investigation. Two parent antagonists (SM-WK86BLMN, SM-SFKXATXC) were re-tested in independent assay runs with determined IC_50_ values in good agreement.

**Table 2:** Summary of analogue exploration of GLP-2R antagonists.

| Parent<br>GLP-2R<br>Antagonists | $IC_{50}$ ($\mu M$) | # Analogues Tested | # Actives | Classification | $IC_{50}$ ranges<br>($\mu M$)<sup>o</sup> |
| --- | --- | --- | --- | --- | --- |
| SM-4MPGPRSA | 0.97 | 4 | 2 | 2 antagonists* | 2.1–2.3 |
| SM-UU87GK35 | 24.0 | 3 | 0 | N/A | N/A |
| SM-WK86BLMN* | 0.07/0.16<sup>†</sup> | 5 | 4 | 4 antagonists | 2.3–4.9 |
| SM-EX12JGV5 | 6.2 | 5 | 2 | 2 antagonists* | 7.8–17 |
| SM-T6ABU679 | 5.5 | 3 | 1 | 1 antagonists* | 9.7 |
| SM-SFKXATXC* | 0.92/0.94<sup>†</sup> | 5 | 2 | 2 antagonists* | 16–23 |

\* = hill slopes (>2) observed in DRC; <sup>†</sup> = assay run1/run2; <sup>o</sup> = Min–Max $IC_{50}$ range observed across active analogues

### C.3 MRGPRX2

Compounds were screened in the MRGPRX2 FLIPR calcium mobilization assay using HEK-293 cells stably expressing MRGPRX2, as described in Appendix A. Primary screening was conducted at three concentrations (0.5, 5, and 50 *µ*M), followed by progression of compounds which gave an initial signal to full dose-response curves for EC_50_ determination. Counter-screening against the parental HEK-293 cell line was performed in parallel to identify and exclude compounds with non-specific or cell-intrinsic activity.

Thirty-eight compounds from the WuXi OTS collection were evaluated against MRGPRX2 for agonist and antagonist activity. Primary screening followed by DRC identified thirteen molecules demonstrating confirmed functional activity against MRGPRX2: ten agonists (EC_50_ 2.2 – 18 *µ*M), with SM-SF1C941G as a representative example (Fig 11), and three antagonists (IC_50_ 9.3 – 19 *µ*M), with SM-6JSFN2T1 as a representative example (Fig 12). Counter-screening in parental HEK-293 cells confirmed the absence of non-specific calcium mobilization.

**Figure 11:**
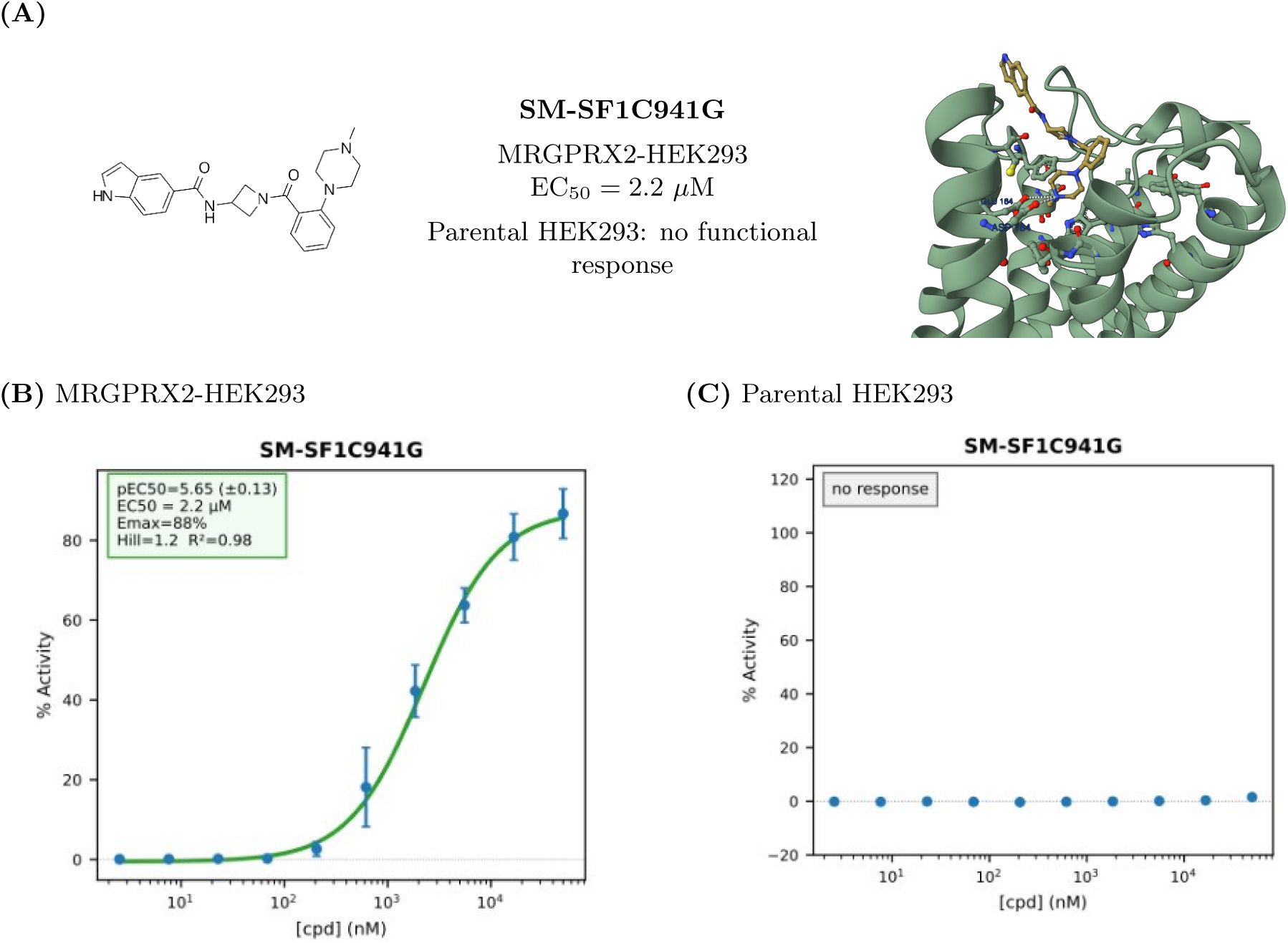
(A) Agonist binder SM-SF1C941G. Dose-response curves for SM-SF1C941G in agonist mode in (B) MRGPRX2-expressing HEK293 cells and (C) parental HEK293 cells, measured by FLIPR calcium mobilization assay.

**Figure 12:**
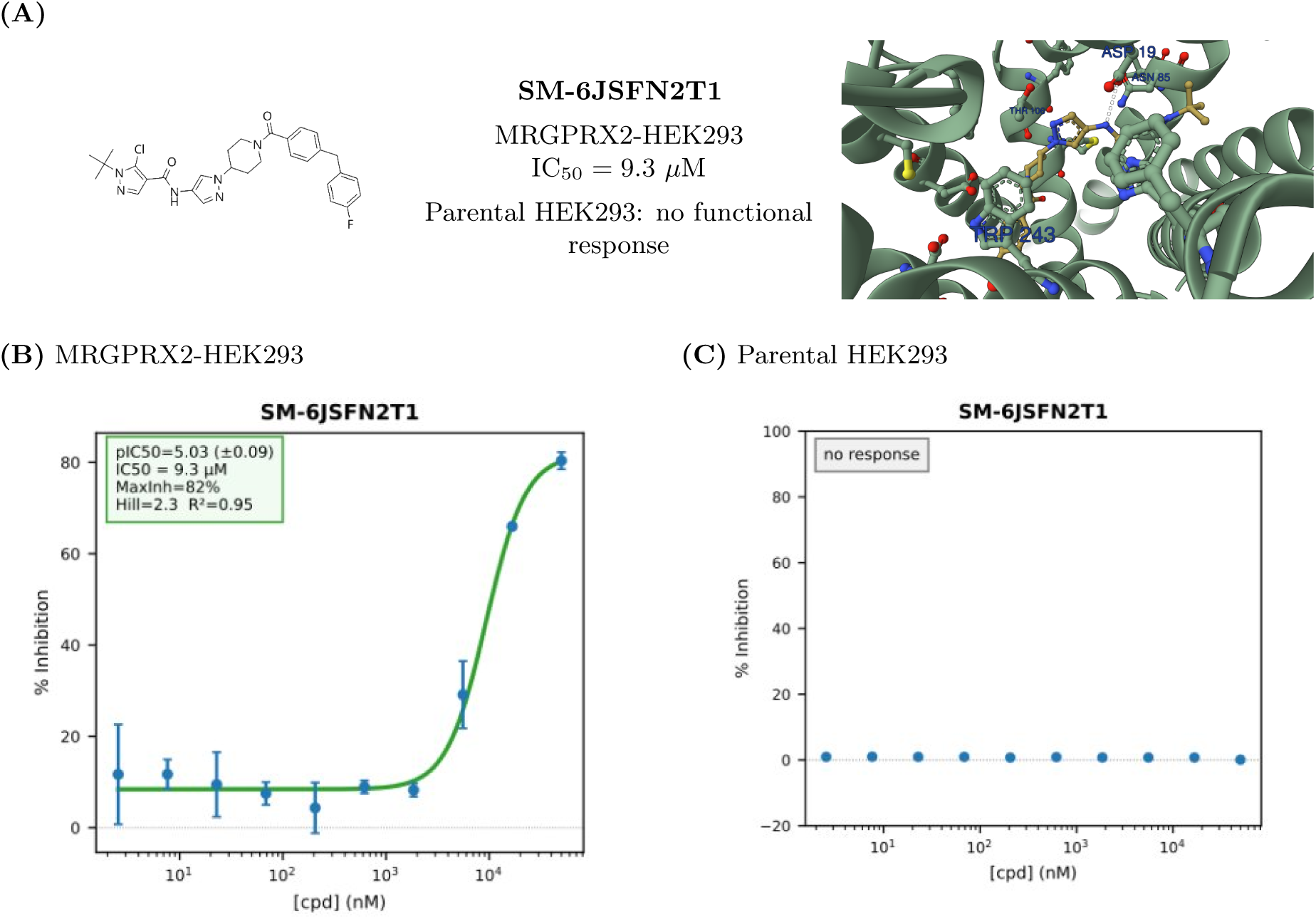
(A) Antagonist binder SM-6JSFN2T1. Dose-response curves for SM-6JSFN2T1 in antagonist mode in (B) MRGPRX2-expressing HEK293 cells and (C) parental HEK293 cells, measured by FLIPR calcium mobilization assay.

Structural analogues of five functionally active agonist parents and three functionally active antagonist parents were retrieved from the WuXi OTS collection using Morgan fingerprint similarity (radius = 2, 2048 bits) and screened in the MRGPRX2 FLIPR assay alongside the parental HEK-293 counter-screen. In total, 18 agonist-parent analogues and 15 antagonist-parent analogues were tested. Across the full set of 33 analogues, none elicited a calcium response in the parental HEK-293 counter-screen, confirming that the functional activity observed was MRGPRX2-dependent. All analogues were retrospectively co-folded using Boltz-2.

The results of the agonist analogue series are summarized in Table 3. Among the agonist series, three of the five yielded at least one active analogue with EC_50_ values broadly comparable to those of their parents. Two of the five did not yield active analogues however for SM-5DLANXS9 only a single analogue was available in the WuXi OTS collection and is therefore not adequately interrogated by this round. Notably, the SM-AGPZRE92 series produced more potent analogues alongside a pharmacological switch. One of its five analogues behaved as an antagonist rather than an agonist. Review of the Boltz-2 predicted binding poses highlighted the agonist-classified analogues converged on a binding mode that closely recapitulated the parent pose, occupying the same subpocket, however, the antagonist-classified analogue was displaced relative to the agonist cluster and predicted to bind in an adjacent region of the pocket. This divergence may offer a post-rationalization for the observed functional reversal. Three parent agonists were re-tested in independent assay runs whereby two demonstrated EC_50_ values in good agreement.

**Table 3:** Summary of analogue exploration of MRGPRX2 agonists.

| Parent MRGPRX2 Agonist | EC<sub>50</sub> (μM) | #Analogues Tested | #Actives | Classification | EC<sub>50</sub> ranges (μM)<sup>φ</sup> |
| --- | --- | --- | --- | --- | --- |
| SM-SF1C941G | 2.2 | 3 | 1 | 1 agonist | 5.0 |
| SM-ZJGLZDJ9 | 9.2/12<sup>†</sup> | 6 | 1 | 1 agonist | 15 |
| SM-5DLANXS9 | 3.2 | 1 | 0 | N/A | N/A |
| SM-51YA5XSM | 15/>50<sup>†</sup> | 3 | 0 | N/A | N/A |
| SM-AGPZRE92 | 18/16<sup>†</sup> | 5 | 3 | 2 agonists<br>1 antagonist | 4.32–4.39<br>21 |

<sup>†</sup> = assay run1/run2; <sup>φ</sup> = Min–Max EC<sub>50</sub> range observed across active analogues

The results of the antagonist analogue series are summarized in Table 4. Just one antagonist parent yielded active analogues, SM-9WNUSCCT where 4/5 analogues were functionally active. This molecule showed good run-to-run IC_50_ reproducibility. Three analogues were identified as antagonists while a single agonist analogue was identified. Further analysis of Boltz-2 predicted binding poses demonstrated the antagonist-classified analogues clustered tightly with the parent pose while the agonist binding pose diverged. Two parent molecules, SM-6JSFN2T1 and SM-E59FAQCU, demonstrated run-to-run variability. These compounds have since been identified as having kinetic solubility below the assay limit of detection (<1.56 µM), which likely accounts for the discrepancy between repeat runs.

**Table 4:** Summary of analogue exploration of MRGPRX2 antagonists.

| Parent<br>MRGPRX2<br>Antagonists | IC<sub>50</sub> (μM) | #Analogues Tested | #Actives | Classification | EC<sub>50</sub> ranges<br>(μM)<sup>o</sup> |
| --- | --- | --- | --- | --- | --- |
| SM-9WNUSCCT | 19/23<sup>†</sup> | 5 | 4 | 1 agonist<br>3 antagonists | 16<br>21–33 |
| SM-6JSFN2T1 | 9.3/>50<sup>†*</sup> | 5 | 0 | N/A | N/A |
| SM-E59FAQCU | 11/>50<sup>†*</sup> | 5 | 0 | N/A | N/A |

<sup>†</sup> = assay run1/run2; <sup>o</sup> = Min–Max IC<sub>50</sub> range observed across active analogues; \* Kin sol = <1.56 μM

### C.4 ROR1

Compounds were screened utilizing ready-to-go WuXi (Crelux) biophysical screening assays: nanoDSF, MST/SpS and SPR as described in Appendix A.

Orthogonal screening of thirty four compounds from the WuXi OTS collection against ROR1 in the aforementioned biophysical assays identified one binder which was active in all three assays. SM-9228HDAG showed a *K_D_* of 5.13 *µ*M by SPR and 1.35 *µ*M by SpS. Furthermore in nanoDSF ROR1 was significantly stabilized with Δ*T_m_* (*^◦^*C) = +4.3. An additional putative binder SM-A776RTUX was identified via measured binding in SPR (*K_D_* = 10.1 *µ*M) and MST (*K_D_* = 8.52 *µ*M) and SpS (*K_D_* = 2.33 *µ*M) however no thermal stabilization was observed in nanoDSF.

Using Morgan fingerprint similarity (radius = 2, 2048 bits) [Morgan, 1965] the WuXi OTS collection was searched for closely related structural analogues of SM-9228HDAG. Analogues spanning Tanimoto similarities of 0.38–0.61 relative to the parent were explored. Three of four analogues were demonstrated to be binders by SpS (*K_D_*s = 10.3 to 48.2 uM) two of which were also demonstrated binding in the MST assay (*K_D_*s 16.9 to 87.7 uM). One analogue SM-RW7VK4A6 induced aggregation of ROR1 in both SpS and MST and thus KD could not be determined. Concurrently, the parent molecule SM-9228HDAG was re-examined in these assays and we note two independent SpS runs were within 2-fold agreement with determined KD values of 0.72 uM and 1.35 uM respectively. SM-9228HDAG has since been determined to have low kinetic solubility and we believe this may contribute to the discrepancy between KD values measured in MST vs SpS for this compound given the sensitivity of MST to aggregation.

Retrospective analysis of the co-folding predictions demonstrated the analogues predicted lower Optimization Scores (relative scoring metric of strongest to weakest binder) than the parent compound SM-9228HDAG, in agreement with the weaker binding affinities observed experimentally by spectral shift assay. The predicted co-folding structure places SM-9228HDAG in a binding pose proximal to key residues, consistent with the capacity to form stabilising interactions (Figure 14). The central amide engages Glu and Asp residues through hydrogen bonding, the terminal pyrazolopyridine is positioned in close contact with the backbone amide of Ile, and the central fluorophenyl ring participates in a *π*-stacking interaction with a neighbouring Phe residue. Analysis of the predicted analogue binding modes enabled the early SAR explored to be rationalized. The predicted hydrogen bonds between the amide and Glu/Asp residues were conserved across all analogues, including those bearing a modified amide position (SM-NDRE3TM5) or a urea bioisostere in place of the amide (SM-ZMEM2L3L). Notably, introduction of pyridyl groups into the aryl rings flanking the central amide or urea in the three active analogues gave rise to additional predicted hydrogen bond interactions. Replacement of the terminal pyrazolopyridine with an acetamide in SM-RW7VK4A6 was predicted to form productive hydrogen bonds with the Ile backbone amide; however, this compound induced aggregation of ROR1, precluding reliable KD determination.The exploration of analogues demonstrated structural modifications were tolerated, and provided preliminary SAR support, further validating that the observed activity is chemically meaningful rather than assay specific.

**Figure 13:**
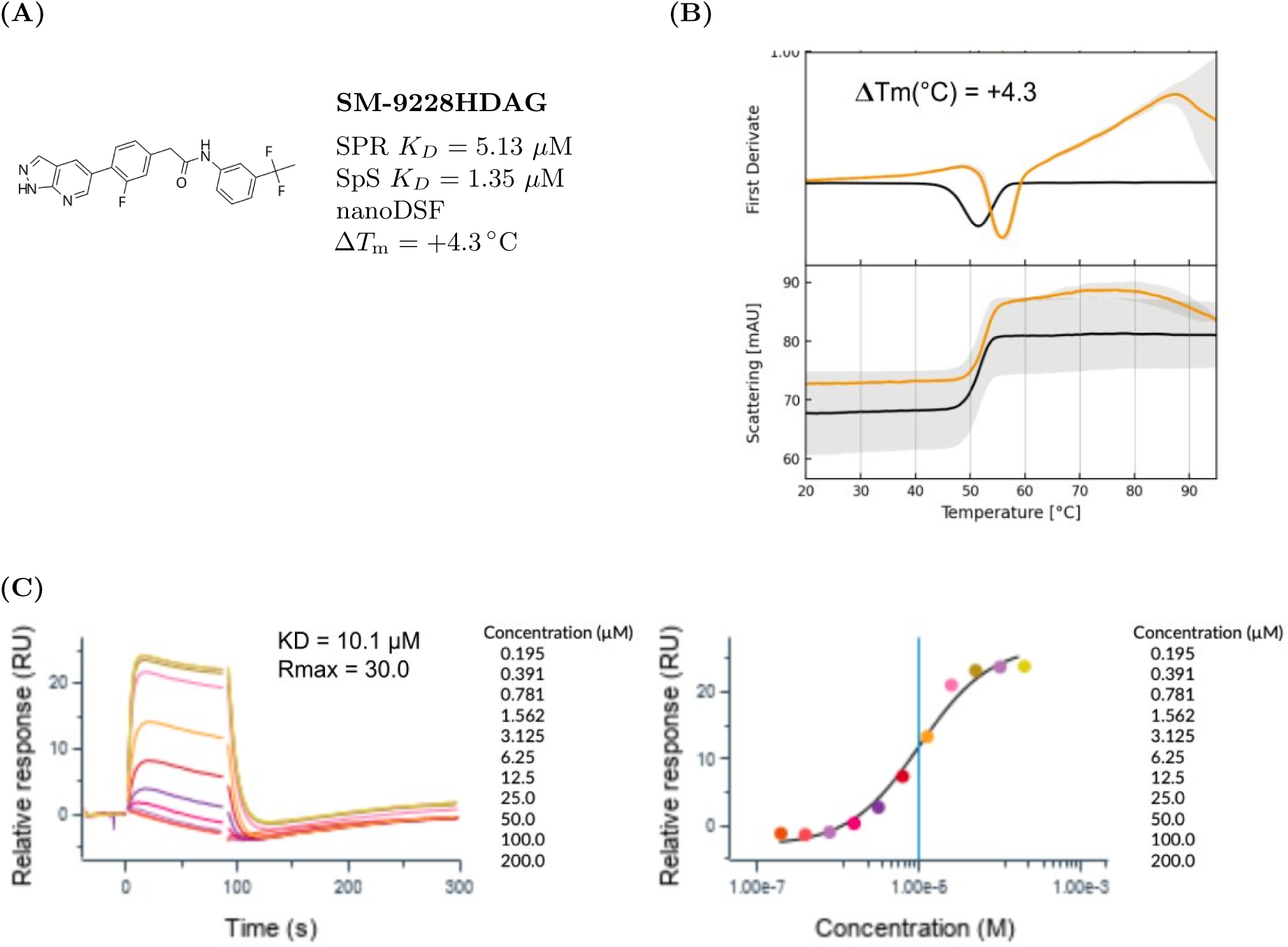
(A) ROR1 binder SM-9228HDAG. (B) Effect of SM-9228HDAG on ROR1 measured by nanoDSF. (C) Binding affinity of SM-9228HDAG to immobilized ROR1 (DFB9, PC16375-1). Serial dilutions of compound (ranging from 200 to 0.195 *µ*M; 2-fold dilution series) were injected onto a SA chip with immobilized ROR1 protein using a multicycle kinetics method.

**Figure 14:**
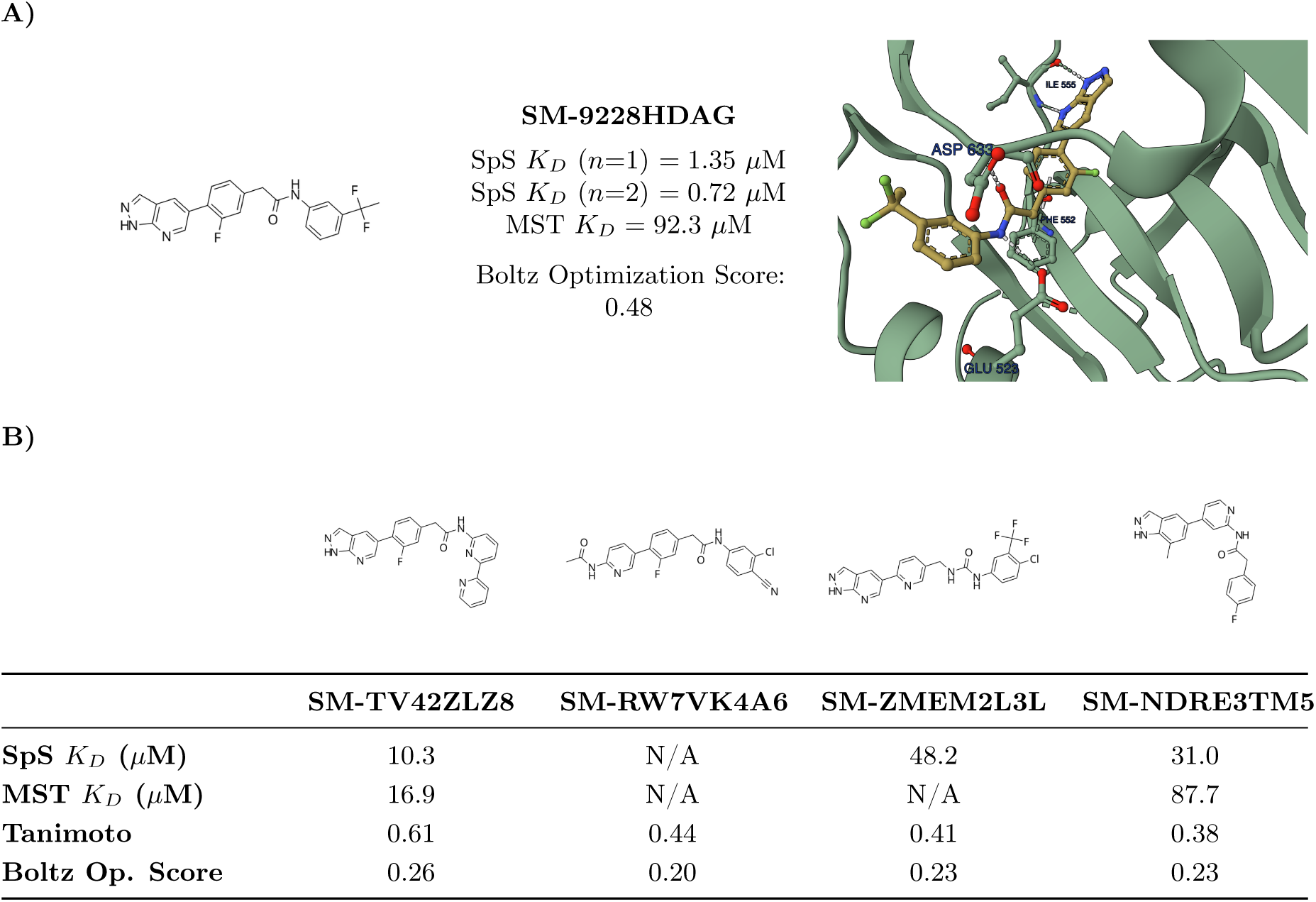
**A)** Boltz Lab predicted structure of SM-9228HDAG bound to ROR1. **B)** Summary of analogues explored of SM-9228HDAG for ROR1 and their experimental results.

### C.5 LC3B and GABARAP

To identify small-molecule binders of LC3B and GABARAP, a generative molecular design screen was performed using Boltz Lab where the molecular generator sampled molecules from Enamine’s 76B REAL space [Enamine Ltd., 2024]. Top-scoring binders were triaged by their ability to satisfy the known pharmacophore. Twenty-eight compounds predicted to engage LC3B or GABARAP were purchased from Enamine and tested in a competitive AlphaScreen assay measuring displacement of biotinylated peptide tracers (FYCO1S for LC3B; K1 for GABARAP) from His-tagged recombinant protein (see Appendix A for assay details). IC50 values determined are averages of *≥* 3 biological replicates.

Of twenty eight compounds tested, three observed inhibition in the AlphaScreen assay with DRCs reaching full saturation and IC50s showing activity in the 15–40 *µ*M range against both targets. In line with previous observations, small molecules are demonstrated to be less potent than peptide binders included here as a positive control (Figure 15c) [Schwalm et al., 2024]. No significant LC3B/GABARAP selectivity was observed; this finding is consistent with the high conservation of HP1 residues across the family [Sora et al., 2020]. Further validation of these compounds as binders via orthogonal assay are being explored.

**Figure 15:**
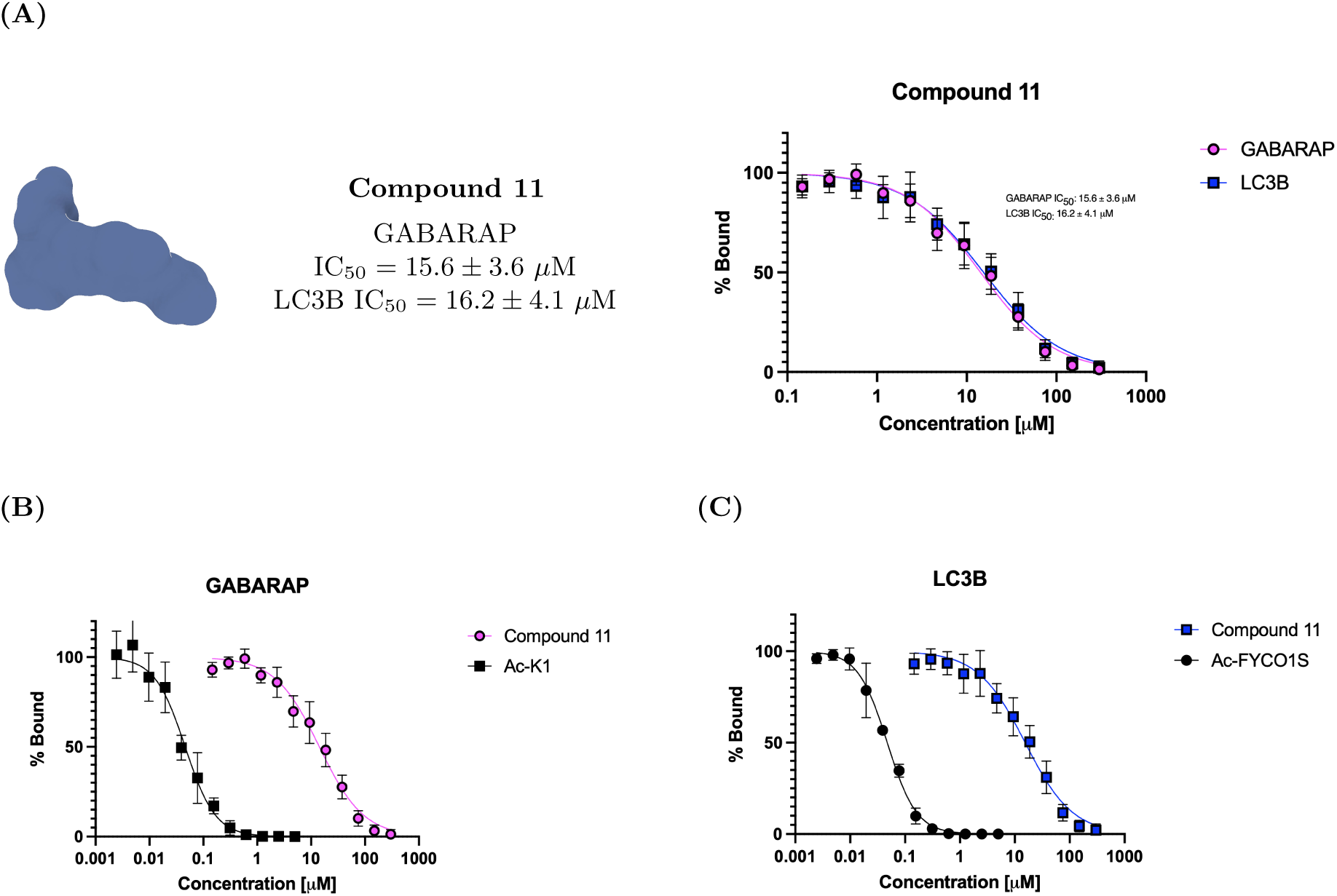
(A) Compound 11 GABARAP and LC3B IC_50_ values observed in the AlphaScreen assay. (B) Compound 11 vs. GABARAP control peptide Ac-K1 (IC_50_ = 45 *±* 4 nM) in the AlphaScreen assay. (C) Compound 11 vs. LC3B control peptide Ac-FYCO1S (IC_50_ = 45 *±* 10 nM) in the AlphaScreen assay. IC_50_ values are averages of *≥* 3 biological replicates.

### C.6 PknB

We initially generated 1,000 candidate inhibitors with the same SynFlowNet pipeline described in [Passaro et al., 2025], with a further 100 analogues identified through scoring the Enamine kinase library with the same pipeline. These were triaged to a purchasable subset using a combination of compound clustering, Tanimoto similarity to known PknB inhibitors, ProLIF interaction-fingerprint analysis of the Boltz-2 poses, RDKit medicinal-chemistry filters (PAINS, BMS, Lilly, GSK), and expert medicinal-chemist review, ultimately selecting 96 compounds for assay. Compounds were first screened in single points in triplicate at a final concentration of 512 *µM* with the KinaseGlo assay (Appendix A.8). The top 18 compounds were then selected for dose-response analysis. Hits were classified as confirmed binders if they showed a K*_i_* < 150*µM*. Sixteen compounds were classified as confirmed actives, with IC_50_ values ranging from 23.2 to 399 *µ*M. K*_i_* values were derived assuming competitive inhibition, ranging from 6.8 to 118 *µ*M. Figure 16 shows the dose-response curve for one of the compounds confirmed by the KinaseGlo assay.

**Figure 16:**
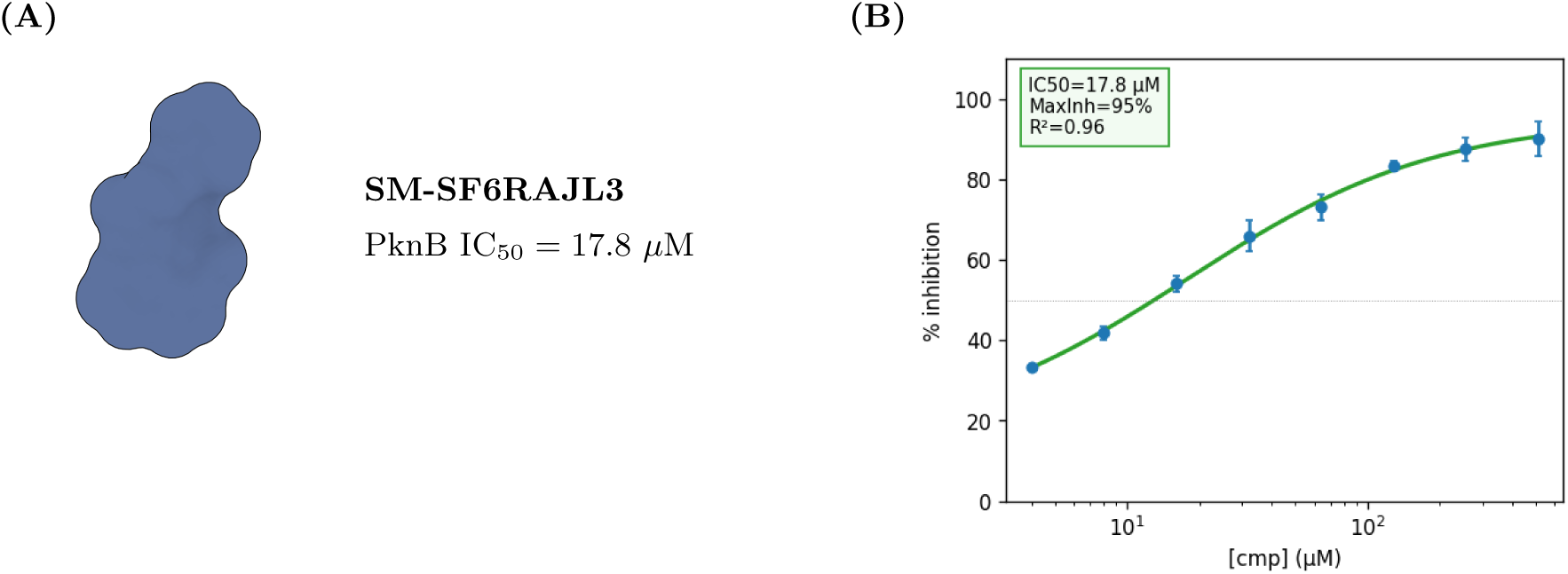
(A) SM-SF6RAJL3 PknB IC_50_ value observed in the KinaseGlo assay. (B) Dose-response curve for SM-SF6RAJL3 in the PknB KinaseGlo assay.

### C.7 MALT1

Compounds were screened utilizing ready-to-go WuXi (Crelux) biophysical screening assays: nanoDSF, MST/SpS and SPR as described in Appendix A.

Orthogonal biophysical screening of thirty-three compounds from the WuXi on-the-shelf (OTS) collection, together with two established MALT1 binders – MLT-985 and mepazine [Schlauderer et al., 2013] – as positive controls, did not yield a confirmed hit. One compound, SM-NQ9S8HFY, was identified as a putative weak binder by SPR (*K_D_* = 101 *µ*M). Orthogonal assessment by MST/SpS did not corroborate binding, and SM-NQ9S8HFY produced no measurable change in MALT1 thermal stability in nanoDSF; signal was further confounded by compound autofluorescence, precluding reliable interpretation of the thermal shift data. This compound was therefore excluded as a putative positive for MALT1.

Collectively, the identification of binders targeting MALT1 has proven challenging within the current screening campaign. One plausible explanation for the low hit rate is the possible under-representation of privileged chemotypes for the MALT1 allosteric pocket within the screened collection, a limitation that may necessitate targeted library expansion or alternative approaches to achieve productive engagement with this site.

### C.8 Experimental ADMET Results

#### Caco-2 permeability

Performance was strongly affected by assay quality: 7 compounds had low solubility and compound recovery below 50%, indicating non-specific loss to plate or membrane and rendering their apparent permeability readouts unreliable. The 8 compounds with measured efflux ratio above 2 were identified as likely efflux substrates whose net A*→*B permeability reflects active transport in addition to passive flux. We accordingly ran the assay in the presence of the efflux inhibitor GF120918 [Hyafil et al., 1993]. Restricting the analysis to the 16 compounds with measurable solubility and using the additional assays substantially increase the performance to *r* = 0.65 and MAE = 0.21. In Figure 17, we compare the Caco-2 experimental results before and after correction using efflux inhibitors.

**Figure 17:**
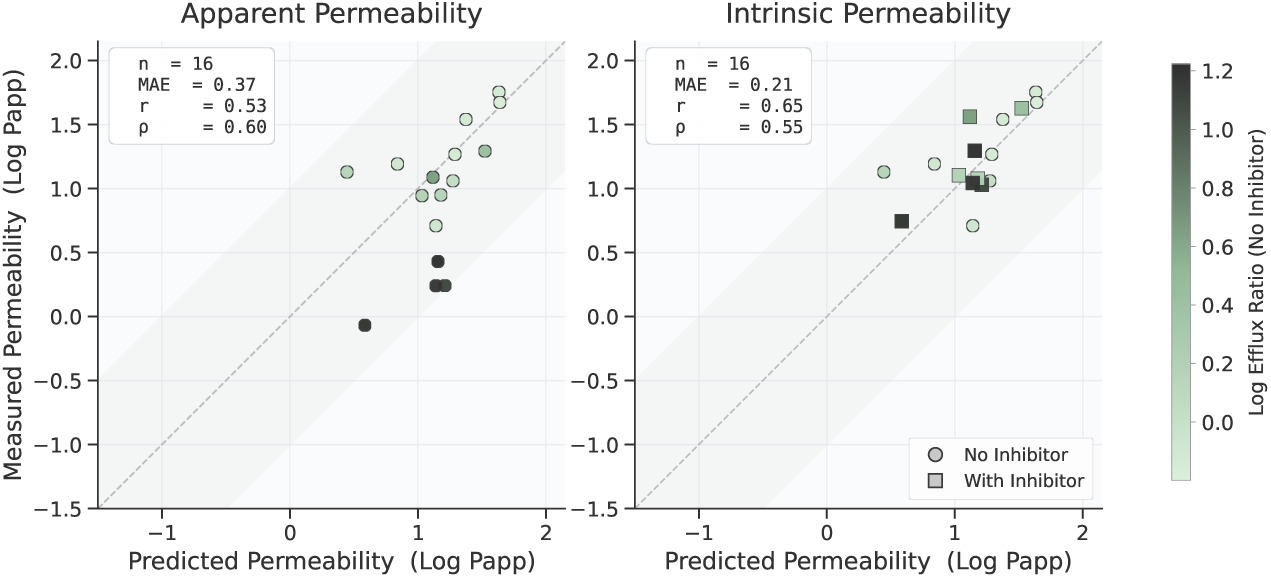
Comparison of predicted vs. experimental Caco-2 permeability before and after correction with efflux inhibitors.

## D ADMET Method

### D.1 Analysis of Data Leakage in TDC

Here we detail the analysis of leakage on the ADMET benchmarks published by the Therapeutics Data Commons (TDC) [Huang et al., 2021] as summarized in Section 3.2.1.

While the scaffold-based splits released by the TDC aim to estimate model performance on unseen chemistry, in practice they are not strict enough to truly assess out-of-distribution generalization. A first issue is structural standardization: when salts, counterions, tautomers, and aromatic representations are not consistently normalized prior to scaffold assignment [Bento et al., 2020], molecules with identical scaffolds (or even fully identical molecules) can end up on opposite sides of the split. A second, more fundamental issue is that scaffold separation does not guarantee that train and test compounds fall outside a single congeneric series; compounds with different Bemis-Murcko scaffolds [Bemis and Murcko, 1996] can still differ by a single atom or substituent, enabling substantial leakage that goes undetected by the traditional zero-scaffold-overlap check.

Table 5 shows representative train-test pairs drawn from the TDC AQSolDB split [Sorkun et al., 2019], illustrating these failure modes directly. The first row is a duplicate compound that appears twice in the dataset with the same label under two distinct tautomeric forms. Under incomplete standardization, these resolve to distinct parent scaffolds and are split between the train and test sets despite being the same compound at assay resolution and having the same measured solubility. The second row shows a pair whose Bemis-Murcko scaffolds are chemically identical but are treated as distinct due to inconsistent aromatic ring representation. The third row is a single-substituent matched molecular pair: distinct scaffolds by the standard definition, but related by a one-fragment substitution in which a pyrimidine ring is swapped for pyridine [Hussain and Rea, 2010]. In these latter two cases, the train and test labels are within a small fraction of a log unit.

**Table 5:**
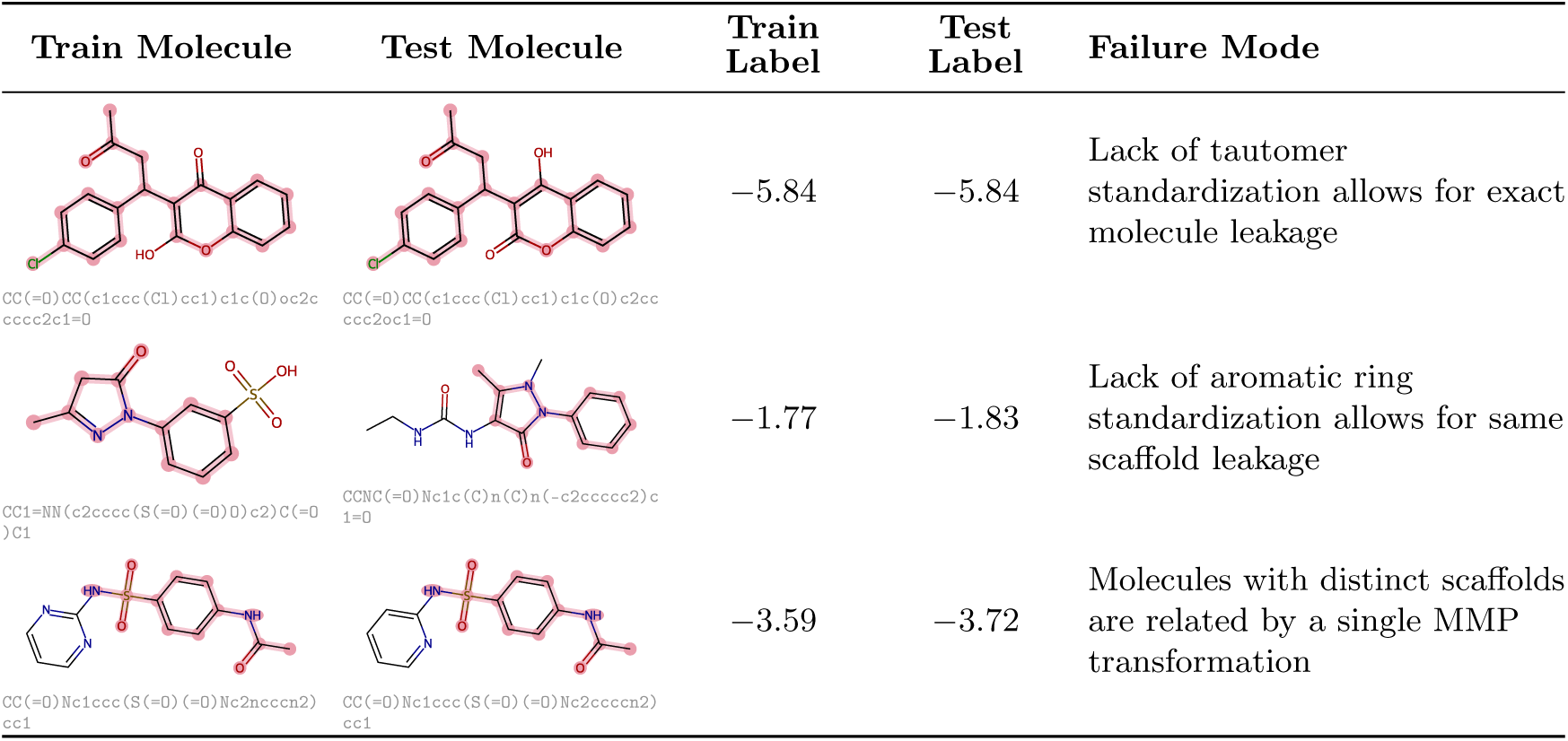
Representative leakage cases from the TDC AQSolDB scaffold splits.

| Train Molecule | Test Molecule | Train Label | Test Label | Failure Mode |
| --- | --- | --- | --- | --- |
| CC(=O)CC(c1ccc(Cl)cc1)c1c(O)oc2ccccc2c1=O | CC(=O)CC(c1ccc(Cl)cc1)c1c(O)c2ccccc2oc1=O | -5.84 | -5.84 | Lack of tautomer standardization allows for exact molecule leakage |
| CC1=NN(c2cccc(S(=O)(=O)O)c2)C(=O)C1 | CCNC(=O)Nc1c(C)n(C)n(-c2ccccc2)c1=O | -1.77 | -1.83 | Lack of aromatic ring standardization allows for same scaffold leakage |
| CC(=O)Nc1ccc(S(=O)(=O)Nc2ncccn2)cc1 | CC(=O)Nc1ccc(S(=O)(=O)Nc2ccccn2)cc1 | -3.59 | -3.72 | Molecules with distinct scaffolds are related by a single MMP transformation |

The most common alternative to scaffold-based splitting is fingerprint similarity based splitting, typically implemented via Butina clustering [Butina, 1999]. This approach guarantees only that every molecule in a cluster falls within a similarity threshold of its centroid; it places no constraint on the pairwise similarity between molecules in different clusters. As a result, two molecules assigned to different clusters (and therefore to different sides of the train-test split) can have unbounded similarity to each other, leaving the same form of near-neighbor leakage that scaffold splits permit.

To address these limitations, we define leakage for each test compound using the union of two complementary criteria. The first is whole-molecule similarity: we flag any test compound whose maximum ECFP4 [Rogers and Hahn, 2010] Tanimoto similarity to the training set exceeds 0.45. The second is local analog overlap: we flag any test compound that is a structural analog of any compound in the training set. Formally, we say that two compounds are structural analogs or connected by a matched molecular pair if their structures differ by exactly one fragment after cutting up to three rotatable bonds in each compound, with the differing fragment comprising at most 50% of the heavy atoms in each molecule [Hussain and Rea, 2010]. The first criterion captures broad similarity that scaffold splits routinely miss; the second captures medicinal-chemistry analog relationships that leave the scaffold nominally distinct. We treat the union as the relevant measure of leakage, since either form of proximity is sufficient to inflate apparent generalization.

When analyzing the LogD, LogS, and Caco-2 ADMET tasks in TDC, using the canonical scaffold splits released with the benchmark, we find evidence of extensive leakage. To standardize compounds prior to the study, we first strip salts and counterions before applying the ChEMBL Structure Pipeline [Bento et al., 2020] to standardize the representations of aromatic rings, charges, and ambiguous functional groups such as sulfoxides. We then strip stereochemistry and use RDKit [Landrum and RDKit Contributors, 2024] to canonicalize tautomers, so that stereoisomers and alternative tautomeric forms of the same compound are not treated as distinct when assessing leakage.

Figure 18 shows the cumulative distribution of maximum train-test ECFP4 Tanimoto similarity across the three tasks. For each test compound, we compute its maximum similarity to any training compound and plot the fraction of the test set falling at or below each threshold. Across all three tasks, a substantial fraction of test compounds have a training neighbor well above the 0.45 threshold: 71.9% on LogD, 55.8% on LogS, and 44.4% on Caco-2. The canonical scaffold splits therefore leave a meaningful portion of the test set within close fingerprint similarity of the training data.

**Figure 18:**
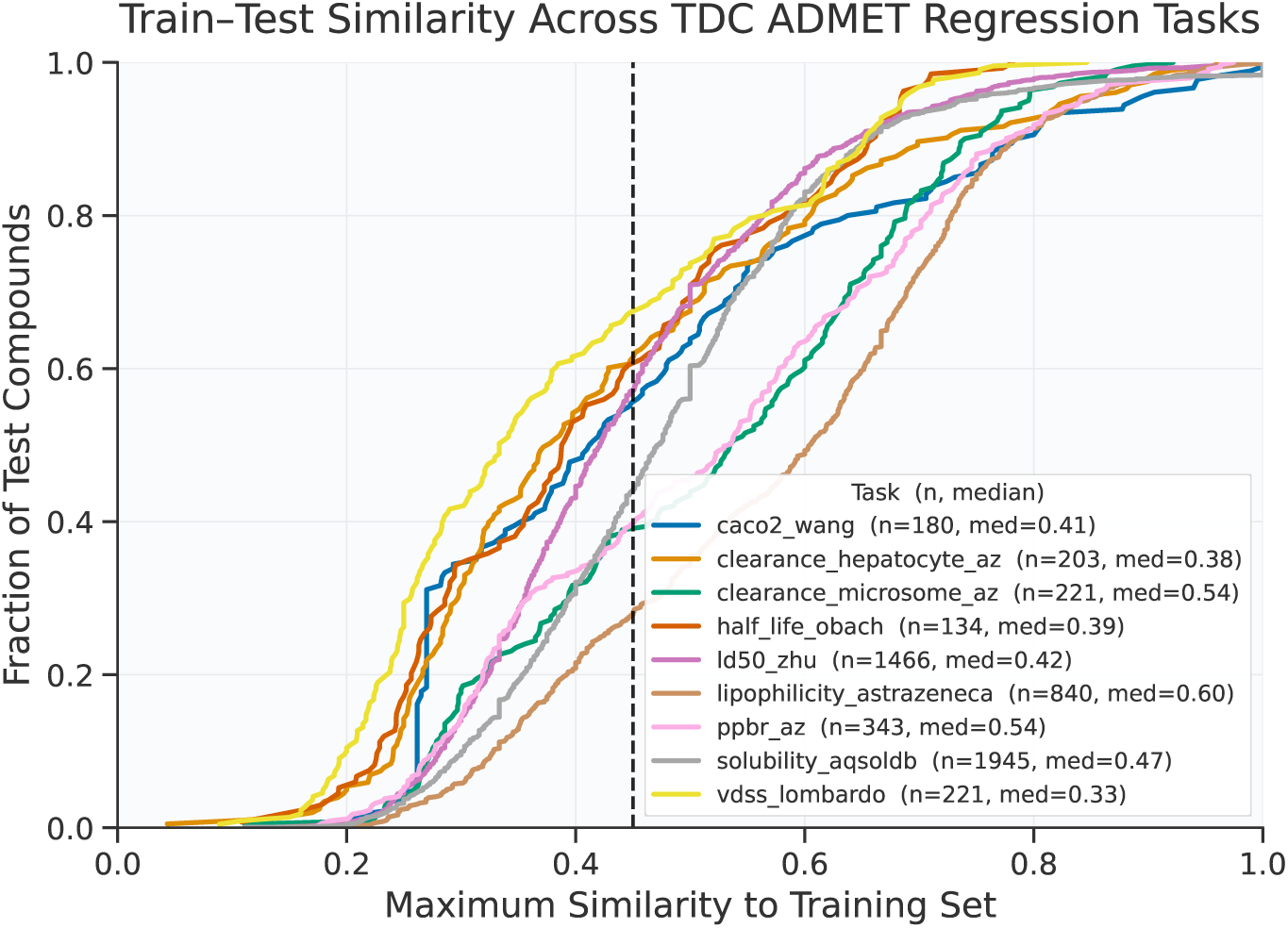
Maximum similarity of test compounds to the training set for TDC ADMET scaffold splits. The dashed vertical line marks the 0.45 threshold used in the leakage definition.

Table 6 reports the prevalence of each leakage criterion across the three tasks. For each test compound, we record whether the training set contains: (i) an exact structural match after standardization, (ii) a compound sharing the same Bemis-Murcko scaffold after standardization, (iii) a matched molecular pair under our analog definition, (iv) any compound with ECFP4 Tanimoto Similarity *>* 0.45, and (v) the union of (iii) and (iv). The union number is the relevant headline figure: across the three tasks 79.9% of compounds in the LogD test set, 71.0% in the LogS test set, and 46.7% in the Caco-2 test set have at least one form of chemically meaningful proximity to the training set despite the canonical scaffold separation.

**Table 6:** Analysis of leakage in TDC ADMET scaffold splits. Each cell reports the percentage of test compounds for which the training set contains a match under the corresponding criterion.

| Task | $n_{\text{test}}$ | Exact Match | Same Scaffold | MMP | Similarity > 0.45 | Union |
| --- | --- | --- | --- | --- | --- | --- |
| Lipophilicity-AZ | 840 | 0.0 | 1.8 | 69.4 | 71.9 | **79.9** |
| Solubility-AqSolDB | 1945 | 0.6 | 27.5 | 54.4 | 55.8 | **71.0** |
| Caco2-Wang | 180 | 1.1 | 1.6 | 32.2 | 44.4 | **46.7** |
| PPBR-AZ | 343 | 0.0 | 2.5 | 51.6 | 60.3 | **64.7** |
| VDss-Lombardo | 221 | 0.0 | 2.7 | 33.9 | 32.6 | **43.4** |
| Half-life-Obach | 134 | 0.0 | 0.0 | 30.6 | 39.6 | **46.3** |
| Clearance-Hep-AZ | 203 | 0.0 | 1.6 | 36.0 | 38.9 | **44.3** |
| Clearance-Mic-AZ | 221 | 0.0 | 0.5 | 61.1 | 61.1 | **67.4** |
| LD50-Zhu | 1466 | 0.0 | 2.5 | 52.7 | 42.8 | **63.6** |

Taken together, these results indicate that strong performance on the canonical TDC scaffold splits is not, on its own, evidence of out-of-distribution generalization for the hit-triage setting. We therefore curated training and evaluation data for our internal models with these failure modes in mind, as described in Section 3.2.2 and Appendix D.2.

### D.2 Data Splitting

As described in Section 3.2.2, our core data corpus is collated from ChEMBL [Zdrazil et al., 2024] and GOSTAR [Excelra (formerly GVK BIO)] across LogS, LogD, Caco-2, and related permeability endpoints. The collated dataset is curated to correct errors in reported values, structural representations, and assay metadata through a combination of manual and automated methods. From the curated pool, we construct training and validation sets across all endpoints, with held-out test sets reserved for the three primary endpoints (LogS, LogD, and Caco-2 A*→*B).

Each validation and test compound is required to be free of leakage to any training compound across the union of all tasks under either of the criteria defined previously in Appendix D.1: maximum ECFP4 Tanimoto similarity greater than 0.45, or the presence of a matched molecular pair. To further strengthen the split, we use the fact that ChEMBL and GOSTAR record the source document for each measurement and partition at the document level rather than the compound level: for any held-out document, we require that no compound in that document has either form of leakage to any compound in any training-assigned document. This guards against the failure mode in which individual pairs all pass the pairwise thresholds but a chain of small changes within an expanded series connects training and held-out chemistry, and ensures that the held-out evaluation is out-of-distribution by construction. Held-out documents are additionally subject to manual review of the underlying sources to ensure that the reported values, units, and assay conditions are consistent and trustworthy.

We additionally handle reference and calibration compounds (e.g., propranolol, verapamil, warfarin) asymmetrically, since their cross-document appearance reflects their role as assay standards rather than genuine chemistry overlap between programs. When a reference compound appears in a training document, the compound is retained in training, but any held-out document containing a compound similar to that reference is excluded from evaluation. However, when a reference compound appears in a held-out document, the compound itself is dropped from evaluation, but the document is retained: excluding entire documents on the basis of containing a calibration standard would systematically remove the most carefully assayed documents from held-out sets, leaving only the least reliable ones.

For additional training data, we incorporate subsets of public and competition datasets that pass the pairwise leakage criteria with respect to our held-out sets. We incorporate solubility data from AqSolDB [Sorkun et al., 2019] and Cui et al. [Cui et al., 2020] and permeability data from PAMPA-NCATS [Sun et al., 2017], primarily to extend coverage of chemistry underrepresented in ChEMBL and GOSTAR. We additionally fold in compounds from recent ADMET prediction efforts under the same leakage constraints: the ASAP Antiviral campaign [ASAP Discovery Consortium, 2024], OpenADMET ExpansionRx [OpenADMET Consortium and Expansion Therapeutics, 2025], and the Biogen ADME-Fang dataset [Fang et al., 2023].

### D.3 ADMET Retrospective Results

As summarized in Section 3.2.4, we benchmark each of our three models against ADMET-AI [Swanson et al., 2024]. Our test sets are constructed under the leakage criteria specified in Appendix D.2 and are out-of-distribution with respect to our training data by construction. However, our splits do not satisfy the same criteria with respect to ADMET-AI, which was jointly trained on nine TDC regression tasks using an ensemble of random splits. To compare how the two models generalize, for each test compound we apply both leakage criteria from Appendix D.1 identifying matched-molecular-pair relationship or global similarity *>* 0.45 against the corpus of TDC training data. We then divide our test set into one subset that is in-distribution for ADMET-AI but out-of-distribution for us, and a subset that is out-of-distribution for both.

#### LogD

Table 7 reports MAE, Pearson *r*, and Spearman *ρ* on the full test set and on the in-distribution and out-of-distribution subsets. LogD is a comparatively easy task and the two models are comparable on the full set (MAE 0.669 vs 0.683; *r* 0.732 vs 0.729). On the in-distribution subset (*n* = 123), ADMET-AI slightly outperforms us across all three metrics (MAE 0.476 vs 0.535; *r* 0.900 vs 0.868); on the out-of-distribution subset (*n* = 626), we slightly outperform ADMET-AI (MAE 0.696 vs 0.724; *r* 0.681 vs 0.659).

**Table 7:**
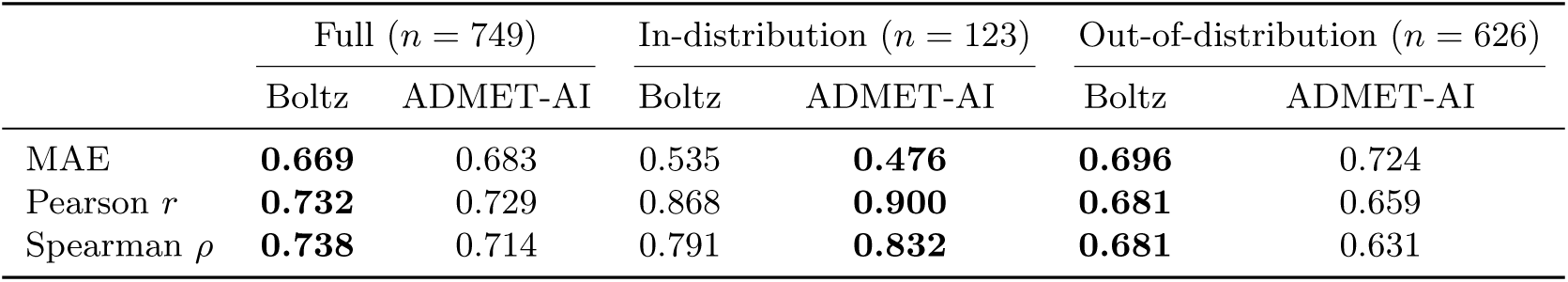
LogD performance for our model and ADMET-AI on the test set, stratified by leakage status with respect to ADMET-AI’s training data. Note that both the splits are out-of-distribution with respect to Boltz.

| | Full ($n = 749$) | | In-distribution ($n = 123$) | | Out-of-distribution ($n = 626$) | |
| --- | --- | --- | --- | --- | --- | --- |
|  | Boltz | ADMET-AI | Boltz | ADMET-AI | Boltz | ADMET-AI |
| MAE | **0.669** | 0.683 | 0.535 | **0.476** | **0.696** | 0.724 |
| Pearson $r$ | **0.732** | 0.729 | 0.868 | **0.900** | **0.681** | 0.659 |
| Spearman $\rho$ | **0.738** | 0.714 | 0.791 | **0.832** | **0.681** | 0.631 |

#### LogS

As stated in Section 3.2.4, to evaluate ADMET-AI as a classifier under the same precision and recall thresholds used for our model, we rank-align its continuous predictions to match the bin sizes our model produces on each subset. Table 8 reports the result on the full test set as well as on the in-distribution and out-of-distribution subsets. Comparing performance across the two settings, ADMET-AI’s high-solubility precision drops from 0.885 in-distribution to 0.451 out-of-distribution, whereas ours drops from 0.808 to 0.726. Low-solubility recall at LogS *< −*5 falls from 0.850 to 0.469 for ADMET-AI and from 0.900 to 0.706 for us. For fully insoluble compounds with LogS *< −*6 recall falls from 1.000 to 0.615 for ADMET-AI versus 1.000 to 0.923 for us. While precision at the top and recall at the bottom are the most informative for the usefulness of a classifier in our virtual screening setting, we also greatly outperform ADMET-AI out-of-distribution on precision and recall for the remaining bins, as listed in Table 8. The comparison is not fully apples-to-apples as ADMET-AI was not optimized for asymmetric triage, while our model was, but the central finding that ADMET-AI’s performance degrades on chemistry far from its training distribution is robust to the specifics of how the models are aligned at evaluation time.

**Table 8:** LogS performance for our model and ADMET-AI on the test set, stratified by leakage status with respect to ADMET-AI’s training data. Note that both the splits are out-of-distribution with respect to Boltz.

| | Full ($n = 751$) | | In-distribution ($n = 140$) | | Out-of-distribution ($n = 611$) | |
| --- | --- | --- | --- | --- | --- | --- |
|  | Boltz | ADMET-AI | Boltz | ADMET-AI | Boltz | ADMET-AI |
| High Solubility Precision | **0.742** | 0.547 | 0.808 | **0.885** | **0.726** | 0.451 |
| High Solubility Recall | **0.404** | 0.298 | 0.356 | **0.390** | **0.421** | 0.261 |
| Moderate Solubility Precision | **0.504** | 0.390 | **0.270** | 0.243 | **0.540** | 0.404 |
| Moderate Solubility Recall | **0.430** | 0.332 | **0.164** | 0.148 | **0.492** | 0.368 |
| Low Solubility Precision ($\text{LogS} < -5$) | **0.407** | 0.282 | **0.234** | 0.221 | **0.456** | 0.303 |
| Low Solubility Recall ($\text{LogS} < -5$) | **0.726** | 0.503 | **0.900** | 0.850 | **0.706** | 0.469 |
| Low Solubility Precision ($\text{LogS} < -6$) | **0.074** | 0.051 | **0.026** | **0.026** | **0.088** | 0.058 |
| Low Solubility Recall ($\text{LogS} < -6$) | **0.929** | 0.643 | **1.000** | **1.000** | **0.923** | 0.615 |

#### Caco-2 A*→*B

When evaluating the Caco-2 performance we additionally stratify by measured efflux ratio (ER), since net A*→*B permeability is a composite of passive physicochemistry and active efflux, and the two regimes pose different prediction problems. ER *<* 2 isolates compounds where passive permeability dominates and the Caco-2 measurement directly reflects the prediction target; ER *>* 2 contains compounds where active efflux materially shapes the readout.

Table 9 reports MAE, Pearson *r*, and Spearman *ρ* for each cell. On the full test set our model exceeds ADMET-AI across every metric (MAE 0.46 vs 0.55; *r* 0.64 vs 0.48; *ρ* 0.65 vs 0.46). The out-of-distribution gap is the dominant driver: on the out-of-distribution ER *<* 2 cell, we exceed ADMET-AI on all metrics (*r* 0.681 vs 0.346); on the in-distribution ER *<* 2 cell, ADMET-AI is stronger (*r* 0.423 vs 0.205; *ρ* 0.470 vs 0.301). In the ER *>* 2 regime, absolute performance collapses for both models (Pearson *r* in the 0.05–0.54 range across cells), and on the in-distribution ER *>* 2 cell (*n* = 55) neither model carries signal (*r* 0.055 vs *−*0.034); we do not interpret differences within this cell. Capturing active efflux remains an open problem, and neither model is yet at a level where it can triage efflux-substrate chemistry without orthogonal data.

**Table 9:** Caco-2 A*→*B permeability performance for our model and ADMET-AI on the test set, stratified by leakage status with respect to ADMET-AI’s training data. Note that both the splits are out-of-distribution with respect to Boltz.

|  | Full |  | In-distribution |  | Out-of-distribution |  |
| --- | --- | --- | --- | --- | --- | --- |
|  | Boltz | ADMET-AI | Boltz | ADMET-AI | Boltz | ADMET-AI |
| *All Compounds (Full $n = 445$, ID $n = 115$, OOD $n = 330$)* |  |  |  |  |  |  |
| MAE | **0.458** | 0.553 | 0.546 | **0.493** | **0.428** | 0.574 |
| Pearson $r$ | **0.639** | 0.483 | **0.298** | 0.281 | **0.719** | 0.533 |
| Spearman $\rho$ | **0.646** | 0.457 | 0.315 | **0.376** | **0.731** | 0.494 |
| *ER < 2 (Full $n = 180$, ID $n = 60$, OOD $n = 120$)* |  |  |  |  |  |  |
| MAE | **0.408** | 0.429 | 0.558 | **0.430** | **0.334** | 0.428 |
| Pearson $r$ | **0.535** | 0.358 | 0.205 | **0.423** | **0.681** | 0.346 |
| Spearman $\rho$ | **0.406** | 0.248 | 0.301 | **0.470** | **0.487** | 0.120 |
| *ER > 2 (Full $n = 259$, ID $n = 55$, OOD $n = 204$)* |  |  |  |  |  |  |
| MAE | **0.497** | 0.643 | **0.533** | 0.562 | **0.487** | 0.665 |
| Pearson $r$ | **0.464** | 0.382 | **0.055** | −0.034 | **0.536** | 0.461 |
| Spearman $\rho$ | **0.420** | 0.340 | 0.039 | **0.080** | **0.484** | 0.389 |

Together, these results suggest that the leakage prevalent in standard ADMET benchmarks materially inflates reported performance, and that the magnitude of the inflation is substantial enough to change conclusions about model utility. The clearest illustration is LogS, where ADMET-AI’s high-solubility precision falls from 0.885 on the in-distribution subset to 0.451 on the out-of-distribution subset, a near halving of the metric most relevant to virtual-screening triage. Similar degradations are visible across LogD and Caco-2 A*→*B, and are not limited to ADMET-AI: any model evaluated under the canonical TDC splits will benefit from the same close-neighbor relationships that our criteria flag as leakage. For ADMET models to be trusted as a triage layer in virtual screening, where the relevant chemistry is by construction far from any individual training program, evaluation must be conducted under splits that reflect what a medicinal chemist would recognize as a genuinely novel compound and not merely a different scaffold, but one without a close analog or near-neighbor in the training set under both global similarity and local SAR criteria. Performance reported under weaker splits should be understood as an upper bound rather than as evidence of generalization to hit-discovery chemistry.

## E Target Analysis

To contextualize the difficulty of each prospective target for Boltz-2’s affinity module, we performed a chemical similarity analysis between the experimentally identified binders disclosed in this work and the compounds the affinity model was trained on. Specifically, for each target we (i) computed the protein sequence similarity between the target and every protein in the affinity training set, (ii) selected, at a series of sequence similarity cutoffs (0.4–0.9), the pool of training proteins meeting that cutoff, (iii) gathered all training-set compounds annotated against those proteins, and (iv) computed the maximum Tanimoto similarity on Morgan fingerprints (radius = 2, 2048 bits) [Morgan, 1965] between each of our experimental binders and that compound pool. The resulting curves report, for each binder, the closest chemical match available to the model among ligands assayed against proteins of progressively higher homology to the target. The shaded band at Tanimoto *≈* 0.4–0.6 (dashed line at 0.5) marks the conventional threshold above which two molecules are typically considered chemically similar; values below this band indicate that the experimental binder is chemically novel relative to anything the model has seen against comparable proteins. We refer to these curves throughout the target-specific discussion below.

Protein sequence similarities were computed with MMseqs2 [Steinegger and Söding, 2017] using easy-search, querying the prospective target sequences against the set of proteins present in the affinity training set, with a minimum sequence identity threshold of 0.4 and –cov-mode 0. The coverage mode setting ensures that when one sequence is a crop or subdomain of another, the alignment coverage is computed relative to the shorter of the two sequences, so that domain-level homologs are not penalized for length differences. The exact command used is:

~~~
mmseqs easy-search query_proteins.fasta target_proteins.fasta \ target_vs_query.m8 tmp --min-seq-id 0.4 --cov-mode 0
~~~

In addition, we verified that none of the experimental binders disclosed in this work form a matched molecular pair with any compound in the affinity training set, applying the MMP definition described in Appendix D.1. This rules out the most common form of trivial chemical leakage (a near-identical training analog differing only by a small R-group substitution) and ensures that the similarity profiles reported below reflect genuinely novel chemical matter rather than memorized scaffolds.

### E.1 STAT6

STAT6 SH2 domain binding centers on a protein-protein interaction surface rather than an enzyme catalytic pocket. Effective ligands must engage the phosphotyrosine-recognition region while balancing polarity (needed for pTyr-like anchoring) to remain in acceptable LogD space, and maintain drug-like permeability and developability.

Homolog coverage is present for STAT3 and 5 in the affinity training set, with only STAT5 (similarity 0.501) that has sequence similarity *>* 0.4 with STAT6. The full set of training-set proteins above the 0.4 cutoff is shown in Table 10.

**Table 10:** Proteins in the affinity training set with sequence similarity *≥* 0.4 to STAT6.

| UniProt | Gene | Similarity | Protein |
| --- | --- | --- | --- |
| P42226 | STAT6 | 1.000 | Signal transducer and activator of transcription 6 (target itself) |
| P51692 | STAT5B | 0.501 | Signal transducer and activator of transcription 5B |

The disclosure of small molecule inhibitors for STAT6 remains low. Publications in 2014, 2015 and 2018 by Morlacchi, Mandal, and Knight et al [Morlacchi et al., 2014, Mandal et al., 2015, Knight et al., 2018] focus on small-molecule phosphopeptidomimetic inhibitors of the SH2 domain of STAT6. Additional disclosure by Nagashima et al. [Nagashima et al., 2007] describes a series of pyrimidine-5-carboxamide and pyrrolopyrimidine derivatives with STAT6 inhibitory activity, progressed across three publications between 2007 and 2009, and AS1517499, a selective tool compound widely used to pharmacologically interrogate STAT6 function in disease models [Chiba et al., 2009]. The recently disclosed PROTAC AK-1690 crystal structure and SAR series developed by Wang et al. [Kaneshige et al., 2025] was reported post-training data cut-off.

The compound similarity analysis (Figure 19) reflects the limited but persistent paralog coverage. The single experimental STAT6 binder (SM-GMUQEHEA) shows a flat maximum Tanimoto of 0.21 across all sequence similarity cutoffs from 0.4 to 0.9, indicating that even at high homology thresholds at least one related STAT family member contributes ligands to the comparison pool. The Tanimoto value sits well below the chemical similarity threshold, however, meaning the disclosed binder is structurally distinct from any STAT-family chemotype the affinity model has been trained on.

**Figure 19:**
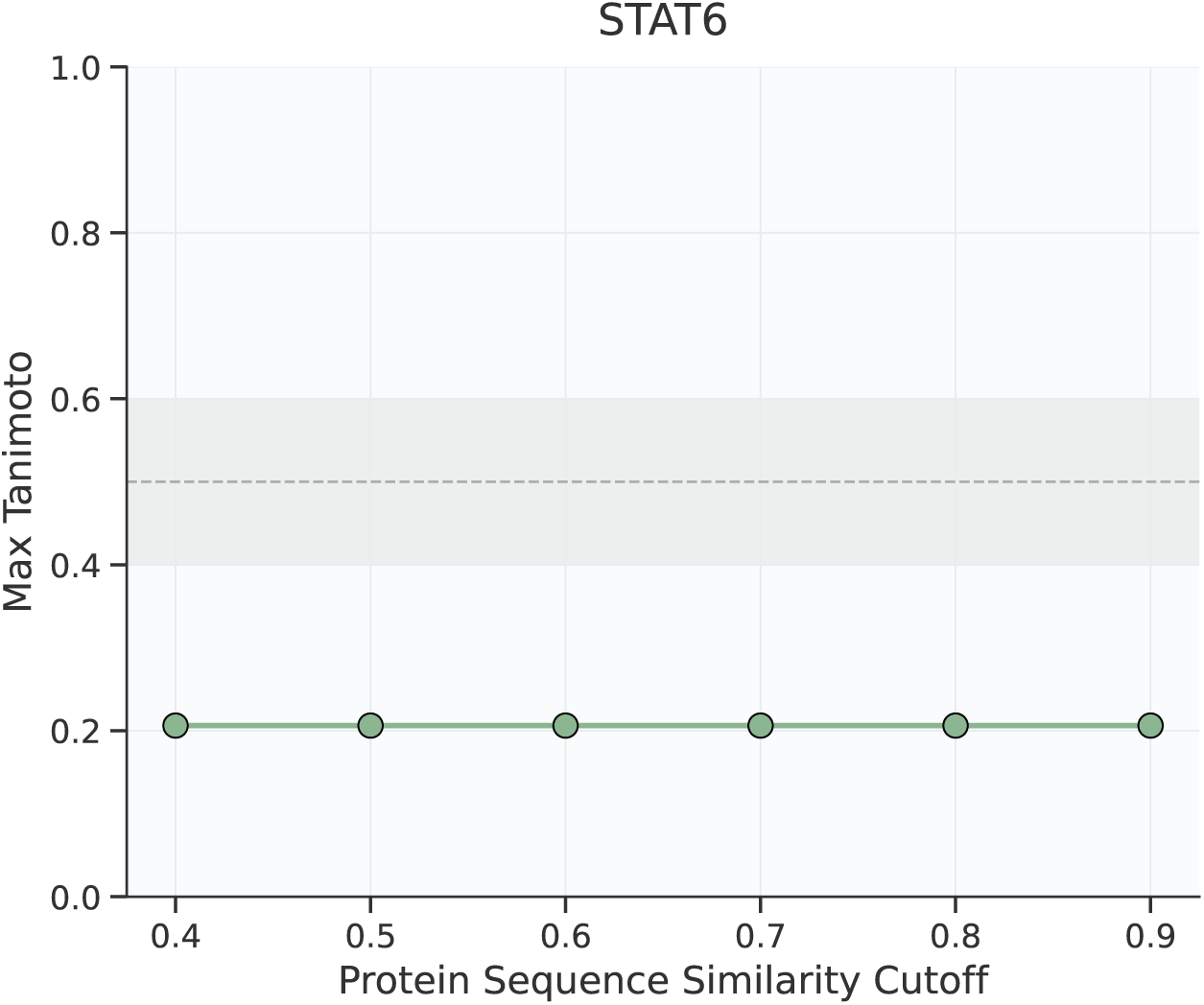
Maximum Tanimoto similarity between the STAT6 experimental binder and affinity training compounds associated with proteins above the indicated sequence similarity cutoff to STAT6.

### E.2 GLP-2R Agonist / Antagonist

The receptor has no known GLP-2R small-molecule co-structure and no experimentally resolved apo state, so affinity prediction must infer whether a druggable transmembrane allosteric pocket exists in a receptor represented only by peptide-bound active-state data. The best template signal comes indirectly from homologous class B receptors, especially GLP-1R, where multiple non-peptide agonist structures demonstrate that oral chemotypes can stabilize transmembrane binding pockets [Kawai et al., 2020]; however, transferring that insight to GLP-2R requires nontrivial generalization across receptor-specific loop and helix differences.

The GLP-2R antagonist similarity profile (Figure 20) supports this picture. The twelve experimental binders show maximum Tanimoto values of 0.21–0.39 at a sequence similarity cutoff of 0.4 and therefore capturing the broader class B GPCR neighborhood including GLP-1R, GCGR (human, mouse, rat) and GIPR — and collapse to zero at cutoff 0.5 and above, reflecting the absence of any GLP-2R paralog in the affinity training set above this threshold (Table 11). At the only cutoff where comparison ligands exist, none of the binders exceed the chemical similarity threshold, so the disclosed antagonists are chemically novel relative to the class B ligand information available to Boltz-2.

**Figure 20:**
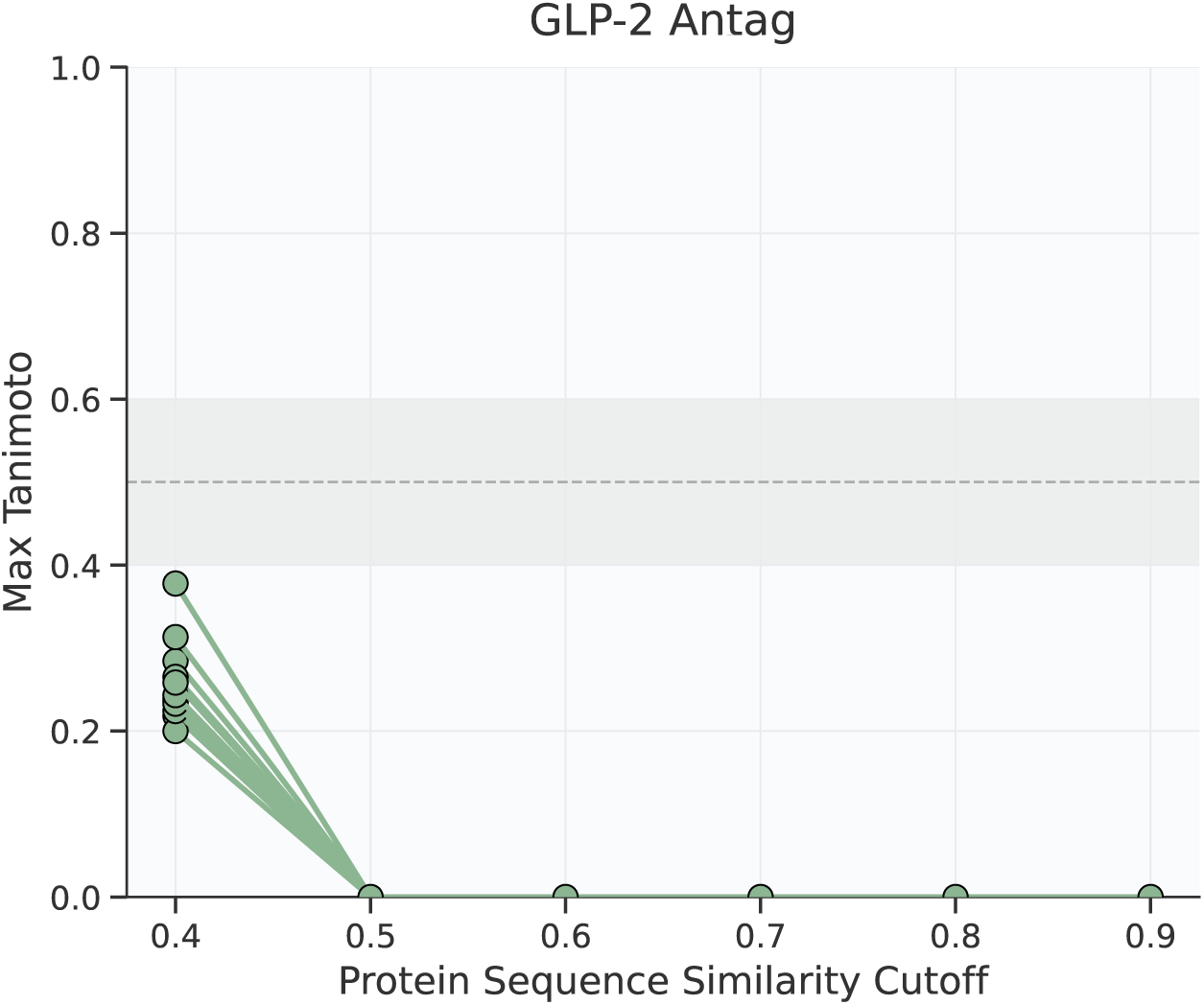
Maximum Tanimoto similarity between GLP-2R antagonist experimental binders and the affinity training compounds associated with proteins above the indicated sequence similarity cutoff to GLP-2R.

**Table 11:**
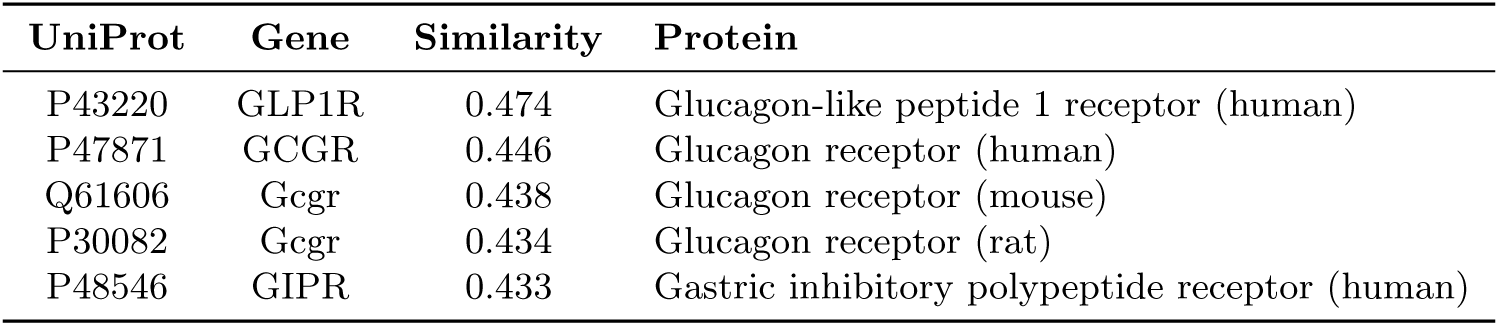
Proteins in the affinity training set with sequence similarity *≥* 0.4 to GLP-2R.

### E.3 MRGPRX2 Agonist / Antagonist

Available MRGPRX2 agonist bound structures are informative but mechanistically incomplete for antagonist design. Current protein backbone arrangements adopt only agonist-bound conformations, with no known small-molecule antagonist structures to draw upon. The orthosteric pocket itself is potentially difficult due to being shallow and highly solvent exposed, and so making it permissive to chemically diverse ligands. It may be expected that in combination with the training data, agonist hits would be favored, or out-rank any antagonist signal within the limited diversity library screened.

Recently disclosed post-training antagonists such as PSB-172656 [Al Hamwi et al., 2025] introduces a potent chemotype, but whose receptor-bound pose remains computationally inferred rather than structurally resolved.

Compound similarity analysis (Figure 21) further illustrates the chemical novelty of the disclosed binders. For both the agonist (ten) and antagonist (three) sets, the maximum Tanimoto similarity to the training-compound pool plateaus around 0.21–0.23 up to a sequence similarity cutoff of 0.6, then collapses to zero at cutoff 0.7 and above. This indicates that no MRGPRX2 paralog with *>* 0.7 sequence similarity contributes ligands to the affinity training set. Across all retained cutoffs the values lie well below the chemical-similarity threshold, so the experimental hits (both agonists and antagonists) represent chemotypes that are essentially out-of-distribution relative to whatever GPCR ligands the model has seen for related receptors. The training-set proteins above the 0.4 sequence similarity cutoff are listed in Table 12.

**Figure 21:**
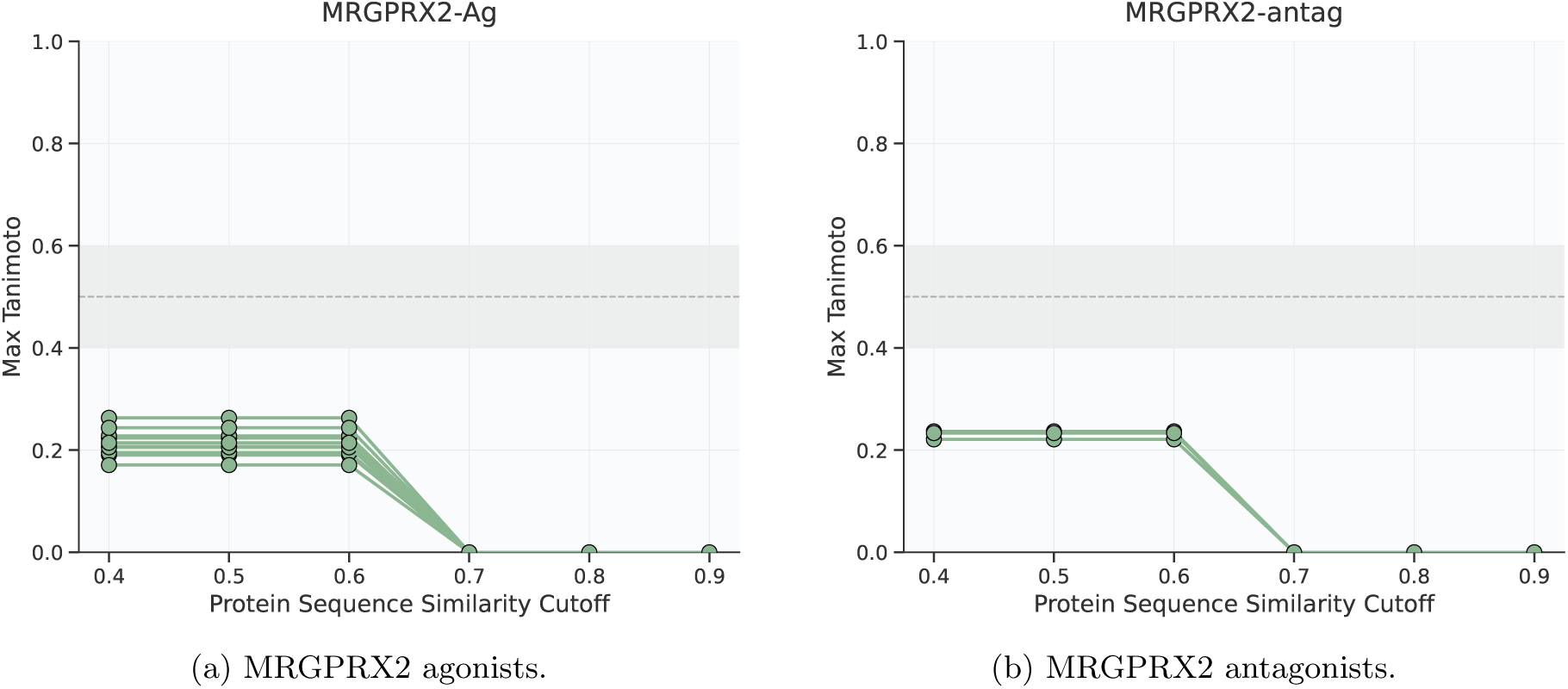
Maximum Tanimoto similarity between MRGPRX2 experimental binders and the affinity training compounds associated with proteins above the indicated sequence similarity cutoff. Above a cutoff of 0.6, no related proteins are present in the training set and the maximum Tanimoto drops to zero.

**Table 12:** Proteins in the affinity training set with sequence similarity *≥*0.4 to MRGPRX2.

| UniProt | Gene | Similarity | Protein |
| --- | --- | --- | --- |
| Q96LB2 | MRGPRX1 | 0.646 | Mas-related G-protein coupled receptor member X1 |
| Q96LA9 | MRGPRX4 | 0.625 | Mas-related G-protein coupled receptor member X4 |

### E.4 ROR1

Utilizing DSF, a screen of 1,486 published kinase inhibitors identified structurally related Ponatinib and GZD824 (both multi-targeted type-II inhibitors) as stabilizers of ROR1. Of these only the crystal structure of Ponatinib has been solved, and remains the sole liganded PDB entry for this target [Sheetz et al., 2020].

ROR1 therefore presents an interesting prospective test for affinity prediction. First, the occluded ATP site disfavors classical type-I kinase chemotypes, so useful molecules must be prioritized for type-II-like occupation of the adjacent hydrophobic pocket. Second, the PDB structure of Ponatinib (PDB: 6TU9) is in the training data, alongside an apo-ROR1 structure (PDB: 5Z55) [Sheetz et al., 2020] therefore any ligand-protein bias is effectively anchored to a single co-crystal (ponatinib-bound 6TU9), creating uncertainty about alternative receptor conformations and ligand poses. Third, ponatinib itself is reported to weakly engage with ROR1, so biased protein-ligand interactions are not optimal to identify strong binders.

While ROR1 has gained traction as a druggable target with a number of recent reports of potent small molecule inhibitors reported [Ghaderi et al., 2023]. Their disclosure and related data post-dates the affinity training date cut-off.

Sequence similarity analysis of the ROR1 pseudokinase domain (UniProt Q01973) at the 0.40–0.47 identity range identifies ten structurally related receptor tyrosine kinases spanning three subfamilies: the Trk neurotrophin receptors, the Eph receptors, and the ALK/LTK family. This is consistent with ROR1’s evolutionary origin as originally designated NTRKR, and reflects shared kinase domain topology across these RTK subfamilies [Lemmon and Schlessinger, 2010].

**Table 13:**
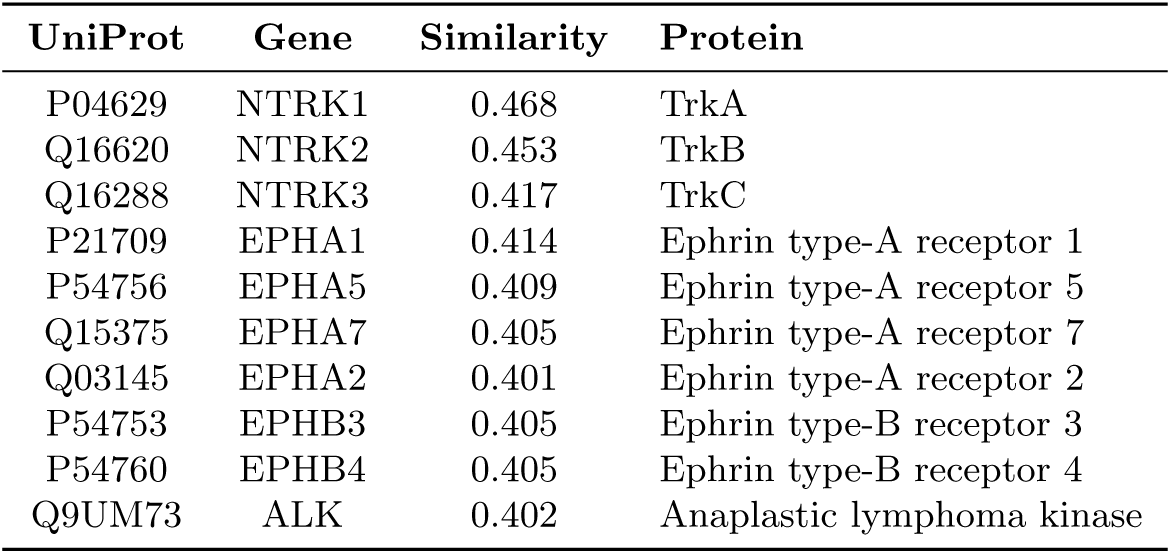
Proteins in the affinity training set with sequence similarity *≥* 0.4 to the ROR1 pseudokinase domain (UniProt Q01973). Reported similarity is the maximum across MMseqs2 hits per UniProt accession.

The chemical similarity profile for ROR1 (Figure 22) is consistent with this homology landscape. At a sequence similarity cutoff of 0.4, capturing the ten RTK homologs above, the maximum Tanimoto similarity between the two experimental ROR1 binders (SM-9228HDAG and SM-A776RTUX) and the corresponding training compound pool reaches 0.38–0.39, just at the edge of the conventional chemical similarity band. Above a cutoff of 0.5, however, no training proteins remain and the maximum Tanimoto collapses to zero, reflecting the complete absence of close ROR1 paralogs in the affinity training set. Boltz-2 therefore approaches ROR1 with chemotype information that is borrowed entirely from cross-subfamily RTKs, and the experimental binders sit on the boundary of what could be considered chemically familiar from this transferred pool.

**Figure 22:**
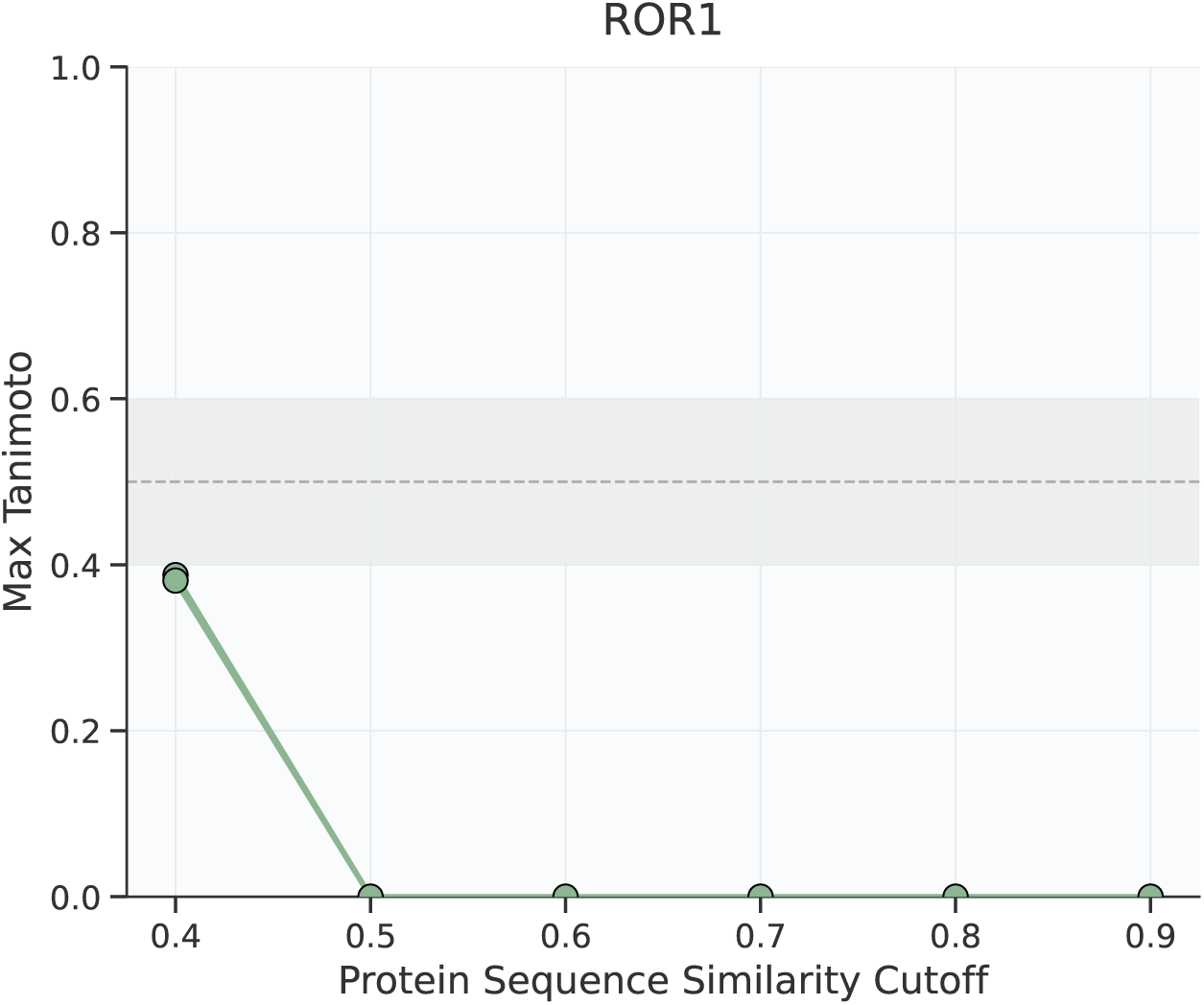
Maximum Tanimoto similarity between ROR1 experimental binders and the affinity training compounds associated with proteins above the indicated sequence similarity cutoff to ROR1. The shaded band (0.4–0.6, dashed line at 0.5) marks the conventional chemical-similarity threshold.

### E.5 LC3B and GABARAP

The LIR docking site is shallow, hydrophobic and historically difficult to drug [Schwalm et al., 2024]. The baseline expectation was not to discover abundant, saturating sub-*µ*M matter on the first pass, and initial data should be interpreted in the context of this challenge.

With respect to prior structural and affinity data, small molecule ligands for LC3/GABARAP have been described since 2019. GW5074 was first reported as an LC3B-recruiting handle in that year [Li et al., 2019], and novobiocin was subsequently reported as a binder of LC3A and LC3B by [Hartmann et al., 2021], with a crystal structure of the LC3A–dihydronovobiocin complex deposited as PDB 6TBE. Three fragment-bound LC3A structures from [Steffek et al., 2023] (PDB 7R9W, 7R9Z, 7RA0) and one covalent LC3B adduct at K49 from [Fan et al., 2021] (PDB 7ELG) were also publicly available before the June 2023 cutoff. The first systematic SAR study of the arylidene-indolinone scaffold, including GW5074 and 18 analogs against both LC3B and GABARAP, was published in January 2025 [Leveille et al., 2025] (bioRxiv February 2024), after Boltz-2’s training cutoff. The XChem fragment screen on LC3B ([Schwalm et al., 2024], PDB 7GA8–7GAS) is similarly post-cutoff. No crystal structure of an arylidene-indolinone bound to LC3B or GABARAP exists at all. The HP1 binding mode is partially engaged by novobiocin but fully shown to be targetable only from post-cutoff 2D-NMR CSP mapping [Leveille et al., 2025]. Boltz-2 therefore had access to several pre-cutoff LC3-family small-molecule structures (predominantly LC3A, and predominantly at HP2 or K49) but no structural data for small-molecule binding at HP1.

Of note, the LIR docking surface has proven tractable to macrocyclic peptide binders: the RFpeptides deep learning design pipeline [Rettie et al., 2025] was used to design macrocycles against GABARAP that achieved *K_D_* values of 6 nM and 36 nM by SPR, with X-ray crystal structures confirming atomic-level agreement with the design models. GAB_D8 and GAB_D23 are post-training-cutoff for Boltz-2 (paper received December 2024, published June 2025) [Rettie et al., 2025].

As shown in Tables 14 and 15, several ATG8 paralogs are present in the affinity training set above the 0.4 cutoff: LC3B and GABARAP each see the other plus LC3A at similarity 0.82, and GABARAP additionally sees GABARAPL2 (GATE-16) at the threshold. Despite this paralog coverage, the compound-side Tanimoto values never approach the chemical-similarity threshold at any cutoff (see Figure 23), indicating that the disclosed hits are chemically novel relative to the historic ATG8-family ligand set the model has been trained on, consistent with the post-cutoff disclosure of the arylidene-indolinone scaffold class [Leveille et al., 2025].

**Figure 23:**
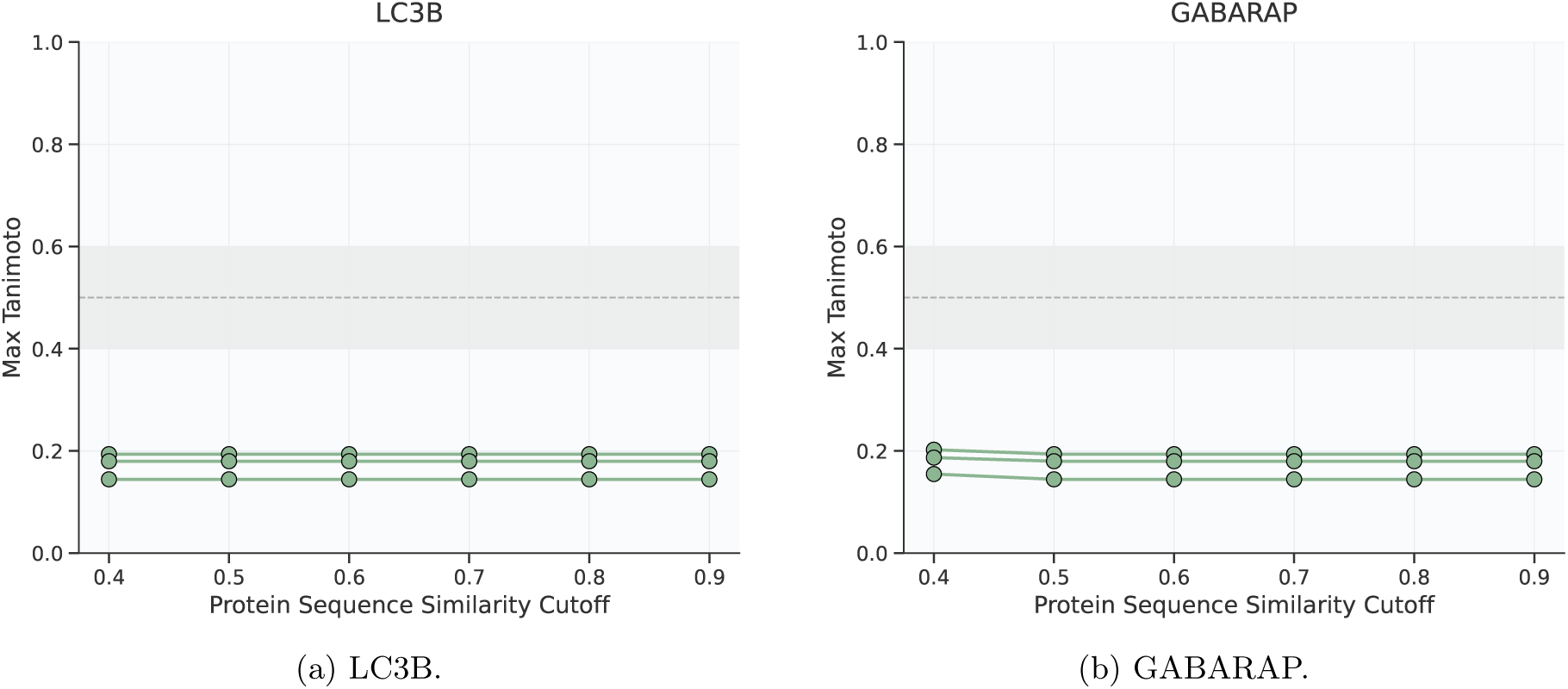
Maximum Tanimoto similarity between LC3B / GABARAP experimental binders (compounds 1, 11, 24) and the affinity training compounds associated with proteins above the indicated sequence similarity cutoff. ATG8 paralogs persist across all cutoffs, but the disclosed binders remain well below the chemical-similarity threshold throughout.

**Table 14:**
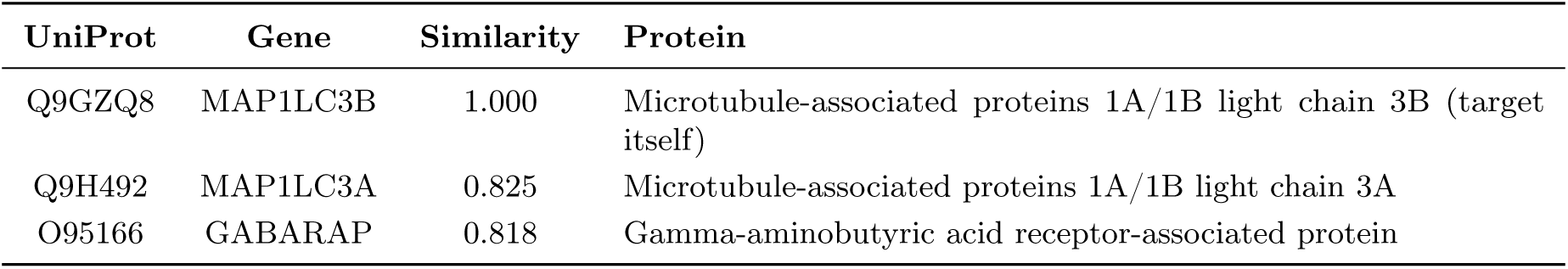
Proteins in the affinity training set with sequence similarity *≥* 0.4 to LC3B.

**Table 15:** Proteins in the affinity training set with sequence similarity *≥* 0.4 to GABARAP.

| UniProt | Gene | Similarity | Protein |
| --- | --- | --- | --- |
| O95166 | GABARAP | 1.000 | Gamma-aminobutyric acid receptor-associated protein (target itself) |
| Q9H492 | MAP1LC3A | 0.818 | Microtubule-associated proteins 1A/1B light chain 3A |
| Q9GZQ8 | MAP1LC3B | 0.818 | Microtubule-associated proteins 1A/1B light chain 3B |
| P60520 | GABARAPL2 | 0.407 | Gamma-aminobutyric acid receptor-associated protein-like 2 (GATE-16) |

### E.6 PknB

PknB is one of eleven serine/threonine protein kinases in *Mycobacterium tuberculosis* and an essential regulator of cell-wall synthesis, division, and metabolism, making it an attractive antitubercular target. Its eukaryotic-like kinase (ELK) domain adopts the canonical bilobal Hanks-type fold and binds ATP in the interlobe cleft, so productive chemistry must address an ATP-competitive site that bears closer resemblance to human kinases than to most bacterial drug targets.

With respect to prior structural data, PknB is comparatively well characterized. Boltz-2’s structure module therefore has direct, in-distribution exposure to liganded PknB ATP-site geometry [Wlodarchak et al., 2018], in contrast to several of the allosteric or undercharacterized sites elsewhere in this study.

As shown in Table 16, the only training-set protein above the 0.4 sequence similarity cutoff is PknB itself, so Boltz-2 approaches this target with in-distribution experience of its own ligand-protein interactions.

**Table 16:**
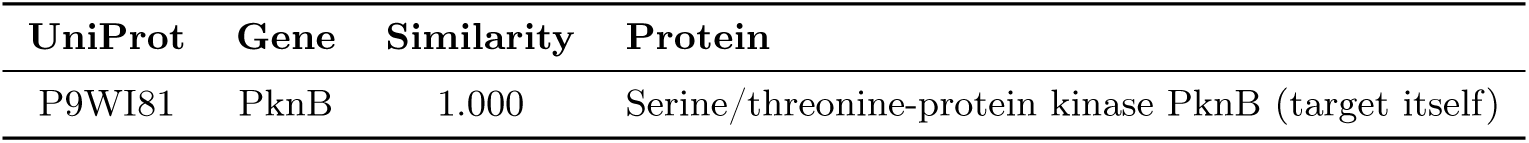
Proteins in the affinity training set with sequence similarity *≥* 0.4 to PknB.

| UniProt | Gene | Similarity | Protein |
| --- | --- | --- | --- |
| P9WI81 | PknB | 1.000 | Serine/threonine-protein kinase PknB (target itself) |

The compound similarity analysis (Figure 24) reflects this. The maximum Tanimoto similarity between the experimental PknB binders and the training-compound pool reaches 0.69; however, most of the binders sit below the 0.5 chemical-similarity.

**Figure 24:**
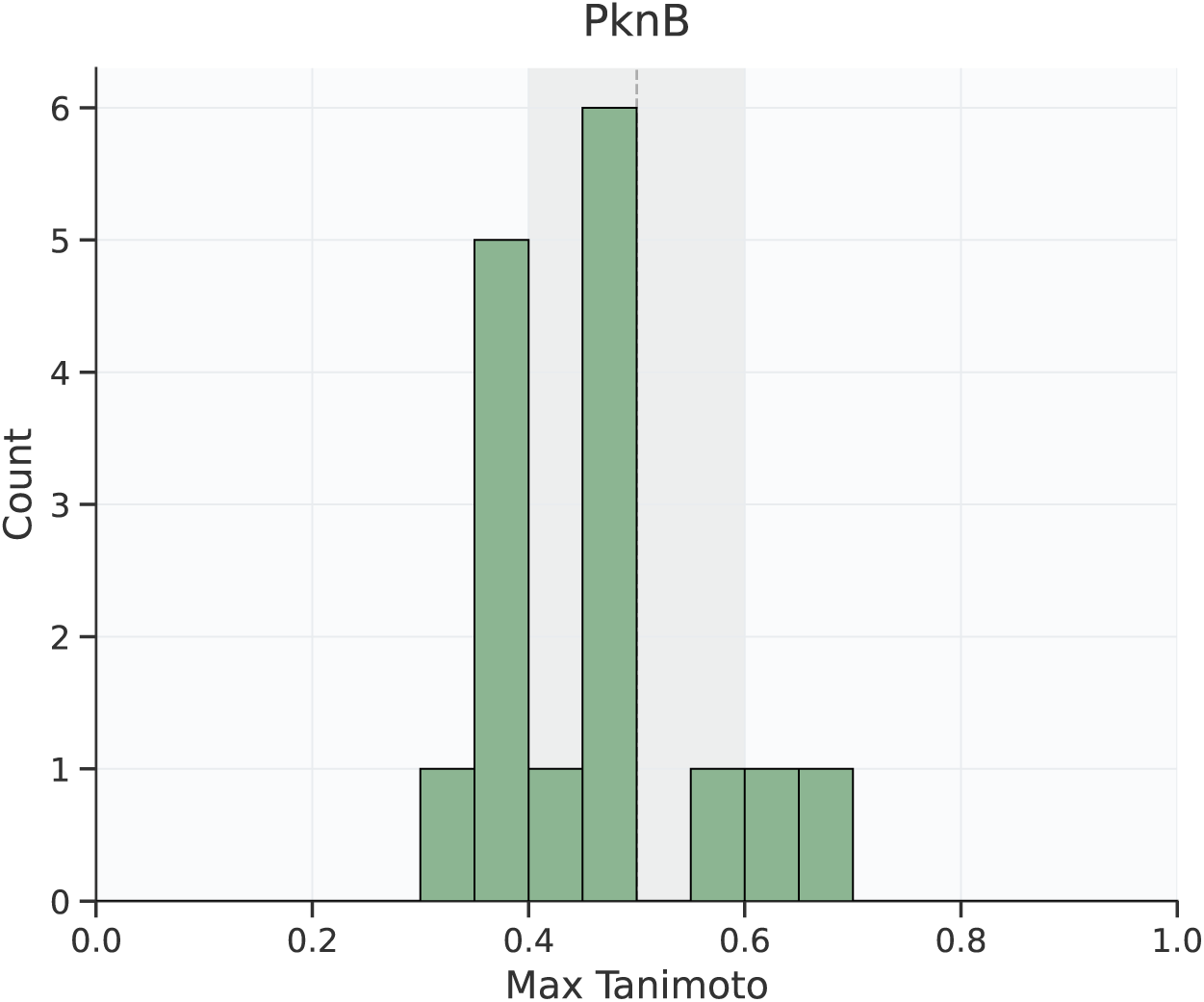
Maximum Tanimoto similarity between PknB experimental binders and the affinity training compounds associated with proteins with sequence similarity above 0.4.

### E.7 mGlu4 PAM

mGlu4 PAM discovery is challenging because productive chemistry must engage a conformationally specific allosteric pocket and support positive modulation rather than simple binding. This makes the screen sensitive to:

1. The structural hypothesis used: Recent work shows mGlu4 PAMs can bind either within a single subunit’s 7TM domain or at the dimer interface between subunits [Niswender and Conn, 2010]. Different PAM chemotypes may prefer different sites, and the same pocket can accommodate both PAMs and agonists.
2. The receptor state represented during ranking: mGlu receptors undergo large conformational rearrangements during activation, transitioning through multiple intermediate states [Niswender and Conn, 2010]. A compound’s functional profile depends on which states it stabilizes.
3. The distinction between binders, modulators, and assay artifacts: Docking can identify compounds that bind the allosteric pocket, but cannot reliably distinguish PAMs (which potentiate glutamate response) from ago-PAMs or pure agonists (which activate directly) [Niswender and Conn, 2010].

In contrast to most other targets in this study, mGlu4 enjoys broad family-level coverage in the affinity training set (Table 17). All eight mGlu subtypes (mGlu1-mGlu8) are present, spanning Group I (mGlu1, mGlu5), Group II (mGlu2, mGlu3), and Group III (mGlu4, mGlu6, mGlu7, mGlu8) receptors [Niswender and Conn, 2010], across human, rat and mouse orthologs. The closest neighbors to mGlu4 are the rat ortholog (P31423, similarity 0.989) and the other Group III receptors mGlu6 (P70579 rat, 0.832; O15303 human, 0.765) and mGlu7 / mGlu8 (similarities 0.75–0.83). Group I and Group II mGlus enter at lower similarities (0.43–0.54). Boltz-2 therefore has access to a rich, in-class chemotype landscape for mGlu PAMs and agonists; the open question for this target is whether the model can use that pool to discriminate the conformationally specific PAM activity profile required, rather than simple class C binding.

**Table 17:** Proteins in the affinity training set with sequence similarity *≥* 0.4 to mGlu4 (GRM4). Group III receptors (mGlu4/6/7/8) cluster at the highest similarities; Group II (mGlu2/3) and Group I (mGlu1/5) follow at progressively lower values.

| UniProt | Gene | Similarity | Protein |
| --- | --- | --- | --- |
| Q14833 | GRM4 | 1.000 | Metabotropic glutamate receptor 4, human (target itself) |
| P31423 | Grm4 | 0.989 | Metabotropic glutamate receptor 4, rat |
| P70579 | Grm6 | 0.832 | Metabotropic glutamate receptor 6, rat |
| O00222 | GRM8 | 0.828 | Metabotropic glutamate receptor 8, human |
| P35400 | Grm7 | 0.785 | Metabotropic glutamate receptor 7, rat |
| Q14831 | GRM7 | 0.785 | Metabotropic glutamate receptor 7, human |
| O15303 | GRM6 | 0.765 | Metabotropic glutamate receptor 6, human |
| P35349 | Grm6 | 0.750 | Metabotropic glutamate receptor 6, mouse |
| Q14416 | GRM2 | 0.538 | Metabotropic glutamate receptor 2, human |
| P31421 | Grm2 | 0.538 | Metabotropic glutamate receptor 2, rat |
| Q14832 | GRM3 | 0.483 | Metabotropic glutamate receptor 3, human |
| P31422 | Grm3 | 0.476 | Metabotropic glutamate receptor 3, rat |
| P41594 | GRM5 | 0.472 | Metabotropic glutamate receptor 5, human |
| P31424 | Grm5 | 0.472 | Metabotropic glutamate receptor 5, rat |
| Q3UVX5 | Grm5 | 0.472 | Metabotropic glutamate receptor 5, mouse |
| H0VDS0 | GRM5 | 0.472 | Metabotropic glutamate receptor 5 (rabbit / Ensembl entry) |
| Q13255 | GRM1 | 0.436 | Metabotropic glutamate receptor 1, human |
| P23385 | Grm1 | 0.432 | Metabotropic glutamate receptor 1, rat |
| P97772 | Grm1 | 0.432 | Metabotropic glutamate receptor 1, mouse |

### E.8 Amylin 3 Receptor Agonist

Beyond conventional GPCR agonist or antagonist hit identification, AMY3 presents an additional complexity due to its composure as a receptor complex rather than a single receptor chain.

No small-molecule-bound crystal structures have been disclosed for AMY3, with known biology dominated by peptide agonism. The available structural data (cryo-EM structures of AMY1, AMY2, and AMY3 in complex with peptide agonists and Gs protein) [Cao et al., 2022] were published in 2022–2023, meaning they fall at or just within the Boltz-2 training cutoff and may be incompletely represented in the model’s learned structural priors for this receptor complex.

This is particularly relevant for affinity prediction, where the absence of any small-molecule co-crystal structure means Boltz-2 has no direct AMY3 small-molecule binding geometry to generalize from, and subtype selectivity dependent on accurately representing RAMP3-shaped binding preferences will be an additional challenge. For a small-molecule screen, that creates uncertainty about the tractability and precise location of productive non-peptide binding modes.

The chemical neighborhood available to Boltz-2 for AMY3 is correspondingly narrow (Table 18). Above the 0.4 sequence similarity cutoff, the only training-set protein meeting the threshold is the calcitonin receptor-like receptor (CALCRL, the receptor scaffold of the CGRP/AM1/AM2 receptor complexes) at similarity 0.609. The calcitonin receptor itself (CALCR), which together with RAMP3 constitutes the AMY3 complex, does not appear in the affinity training set; CALCRL is therefore the closest available paralog providing chemotype information.

**Table 18:** Proteins in the affinity training set with sequence similarity *≥* 0.4 to AMY3 (CALCR).

| UniProt | Gene | Similarity | Protein |
| --- | --- | --- | --- |
| Q16602 | CALCRL | 0.609 | Calcitonin receptor-like receptor (CGRP/AM1/AM2 receptor scaffold) |

### E.9 MALT1

Despite validated active-site and allosteric strategies, current MALT1 medicinal chemistry remains concentrated in a limited set of scaffold families [O’Neill et al., 2023]. The allosteric Trp580 pocket has been the dominant focus of structural drug discovery, and the published co-crystal landscape reflects this. Of the 12 small-molecule-liganded MALT1 structures in the PDB released before the Boltz-2 structural training cutoff (June 2023), 10 occupy the allosteric Trp580 pocket and only 2 covalent small molecules bound active site structures have been deposited.

Boltz-2’s affinity module is trained on a hybrid dataset drawn from ChEMBL, BindingDB [Liu et al., 2007], and PubChem [Kim et al., 2023], filtered to high-confidence biochemical and functional assays. MALT1 small molecule ligands are present in this affinity training set, and remains the sole protein within the dataset with sequence similarity *>* 0.4 to itself. The Boltz-2 model therefore has direct, in-distribution experience with MALT1 ligand-protein interactions, with no closely related surrogate targets diluting or confounding the learned representations.

Boltz-2’s structure module was trained on all PDB entries released before June 2023 [wwPDB consortium, 2023]. For MALT1, this encompasses 16 structures total, of which 12 carry drug-like or probe ligands.

Four additional liganded structures (8V4X, 9MKC, 9MKD, 9MKE - all allosteric) were released after the training cutoff and are not in the Boltz-2 structural training set.

Affinity training data (ChEMBL/BindingDB/PubChem): ChEMBL contains 3,521 bioactivity records against MALT1 (CHEMBL3632452 ChEMBL) [Zdrazil et al., 2024], of which 1,503 are quantitative measurements (IC50, Ki, Kd, EC50) across 114 biochemical or functional assays, covering 1,202 unique compounds. Approximately 900 of these quantitative records derive from publications prior to 2023 and are therefore likely represented in the affinity training set. The affinity data spans a range of privileged chemotypes to the MALT1 allosteric Trp580 binding pocket, including phenothiazines, pyrazolopyrimidines, cyclohexane-diamines, chromane ureas, and others [Zhang et al., 2025].

Boltz-2’s pocket representation for MALT1 is well-calibrated for the allosteric Trp580 site. It has seen 10 diverse co-crystal structures and 900 quantitative affinity measurements, giving it strong in-distribution confidence for ranking allosteric candidates. The primary focus for including this target was to see if the model could rank and find new chemotypes for a well characterized pocket, but that lacks a large family representation (such as with the kinome). Consistent with this, the only training-set protein above the 0.4 sequence similarity cutoff is MALT1 itself (Table 19); no caspase-, paracaspase-, or other cysteine-protease relatives meet the threshold.

**Table 19:** Proteins in the affinity training set with sequence similarity *≥* 0.4 to MALT1.

| UniProt | Gene | Similarity | Protein |
| --- | --- | --- | --- |
| Q9UDY8 | MALT1 | 1.000 | Mucosa-associated lymphoid tissue lymphoma translocation protein 1 (target itself) |

### E.10 Nav 1.8

Targeting Nav1.8 via a small molecule screen focussed on the VSD2 domain presents a significant structural gap for affinity prediction. Despite suzetrigine’s clinical success and sub-nanomolar potency (IC50 = 0.68 nM in human DRG neurons) [Osteen et al., 2025], no co-crystal structure of a small molecule bound to VSD2 exists in the PDB. The available structures include apo Nav1.8 (PDB: 7WFW, 9DBK) [Huang et al., 2022] and a complex with ProTx-I (PDB: 9DBN) [Neumann et al., 2025], a 35-residue tarantula venom peptide that engages VSD2. While ProTx-I confirms the binding site location and identifies contact residues (P691, T692, E694, A695, Q698, I702, L744, G745, V746, A747, K748, L752), peptide-protein interfaces differ fundamentally from small molecule binding modes in geometry, buried surface area, and interaction chemistry.

The sole liganded small molecule structures for Nav1.8 (PDB: 7WE4, 7WFR) [Huang et al., 2022] capture A-803467, a pore-blocking inhibitor that binds the central cavity of the pore domain, and thus not within the target-crop used for this work, and presents a mechanistically and spatially distinct site from VSD2. These pore-binder co-crystals provide no information about VSD2 ligand poses or receptor conformations relevant to suzetrigine-class molecules. A VSD2-directed virtual screen may however extrapolate from the ProTx-I peptide interface and cross-family VSD structures (e.g., Nav1.7 VSD4 with small molecules) [Ahuja et al., 2015].

Nav1.8 VSD2 thus represents a test case for structure prediction in a clinically validated but structurally undercharacterized allosteric site.

Although the VSD2 site itself lacks small-molecule co-crystal data, the broader Nav family is extensively represented in the affinity training set (Table 20). Nav subtypes (Nav1.1-Nav1.8) appear, spanning human, rat and mouse orthologs, with sequence similarities to Nav1.8 ranging from 0.70 (across the Nav1.1-Nav1.7 cluster) up to 0.88 (rat / mouse Nav1.8 orthologs at 0.875 and 0.868). This means Boltz-2 has been exposed to a deep pool of pore-blocker chemotypes from related Nav channels - but, as discussed above, those structural priors derive almost exclusively from central-cavity binders (e.g. A-803467 class) rather than VSD-engagers, so paralog coverage at the sequence level does not translate into prior knowledge of the relevant binding mode. Three lower-similarity hits (Q9P0X4, NP_066921 / O95180) correspond to the T-type voltage-gated calcium channel (CACNA1I) and reflect the shared 24-helix transmembrane architecture between Nav and Cav *α*-subunits [Catterall et al., 2020] rather than functionally relevant ligand overlap.

**Table 20:** Proteins in the affinity training set with sequence similarity *≥* 0.4 to Nav1.8. Sodium channel paralogs dominate; the lower-similarity entries correspond to T-type voltage-gated calcium channels sharing the canonical four-domain channel architecture.

| UniProt | Gene | Similarity | Protein |
| --- | --- | --- | --- |
| Q9Y5Y9 | SCN10A | 1.000 | Sodium channel Nav1.8, human (target itself) |
| Q62968 | Scn10a | 0.875 | Sodium channel Nav1.8, rat |
| Q6QIY3 | Scn10a | 0.868 | Sodium channel Nav1.8, mouse |
| Q14524 | SCN5A | 0.757 | Sodium channel Nav1.5, human (cardiac) |
| P35498 | SCN1A | 0.726 | Sodium channel Nav1.1, human |
| P35499 | SCN4A | 0.720 | Sodium channel Nav1.4, human (skeletal muscle) |
| Q9NY46 | SCN3A | 0.720 | Sodium channel Nav1.3, human |
| Q9UQD0 | SCN8A | 0.720 | Sodium channel Nav1.6, human |
| P04775 | Scn2a | 0.718 | Sodium channel Nav1.2, rat |
| Q99250 | SCN2A | 0.718 | Sodium channel Nav1.2, human |
| P15390 | Scn1a | 0.713 | Sodium channel Nav1.1, rat |
| Q15858 | SCN9A | 0.713 | Sodium channel Nav1.7, human |
| Q62205 | Scn8a | 0.705 | Sodium channel Nav1.6, mouse |
| O08562 | Scn9a | 0.697 | Sodium channel Nav1.7, mouse |
| Q01118 | SCN7A | 0.488 | Sodium channel Nav2.1 / Nax, human (atypical Na sensor) |
| Q9P0X4 | CACNA1I | 0.429 | Voltage-gated calcium channel Cav3.3 (T-type) |

## F Boltz API Usage

Full details and instructions for implementation of the Boltz API can be found online: https://api.boltz.bio/docs/guides/small-molecule-library-screen

API Keys must be generated prior to launching a request: https://api.boltz.bio/docs/guides/getting-started

All Boltz API inputs in Appendix F were validated against the Boltz Python SDK (boltz-api 0.34.1); each target submitted and completed successfully on the live API.

An up-to-date WuXi OTS compound collection can be requested from the WuXi LabNetwork: https://www.labnetwork.com.cn/library

### F.1 Example API Call

Following is an example API input for the BoltzMol-1 screen for ROR1 using a defined list of SMILES strings.

Listing 1: Submitting a defined list of ROR1 small-molecule inhibitors with the Boltz API.

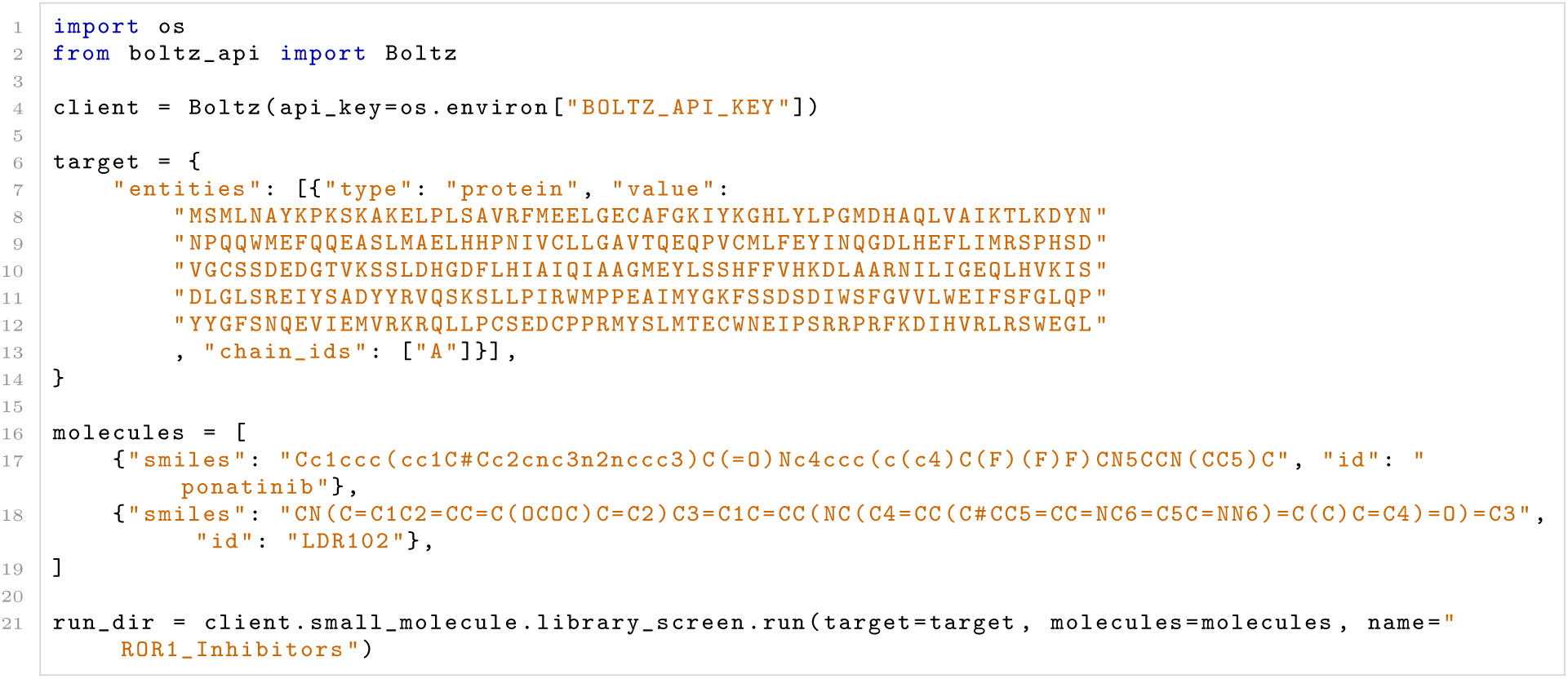

### F.2 Full ROR1 BoltzMol-1 API Call

Following is the full API call to replicate the BoltzMol-1 screen on ROR1, importing a CSV file containing the WuXi OTS screening collection as reported in this manuscript.

Listing 2: Boltz library-screen request body (full input format) for ROR1.

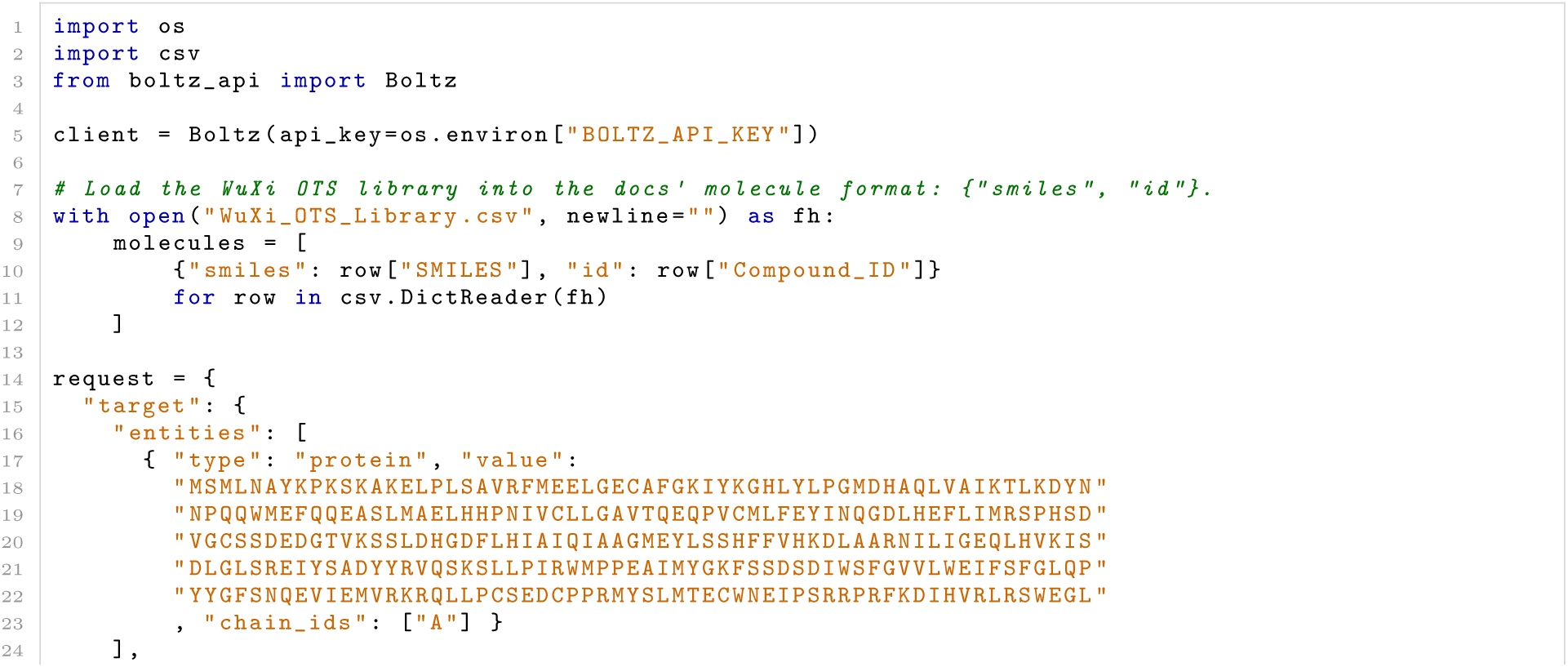
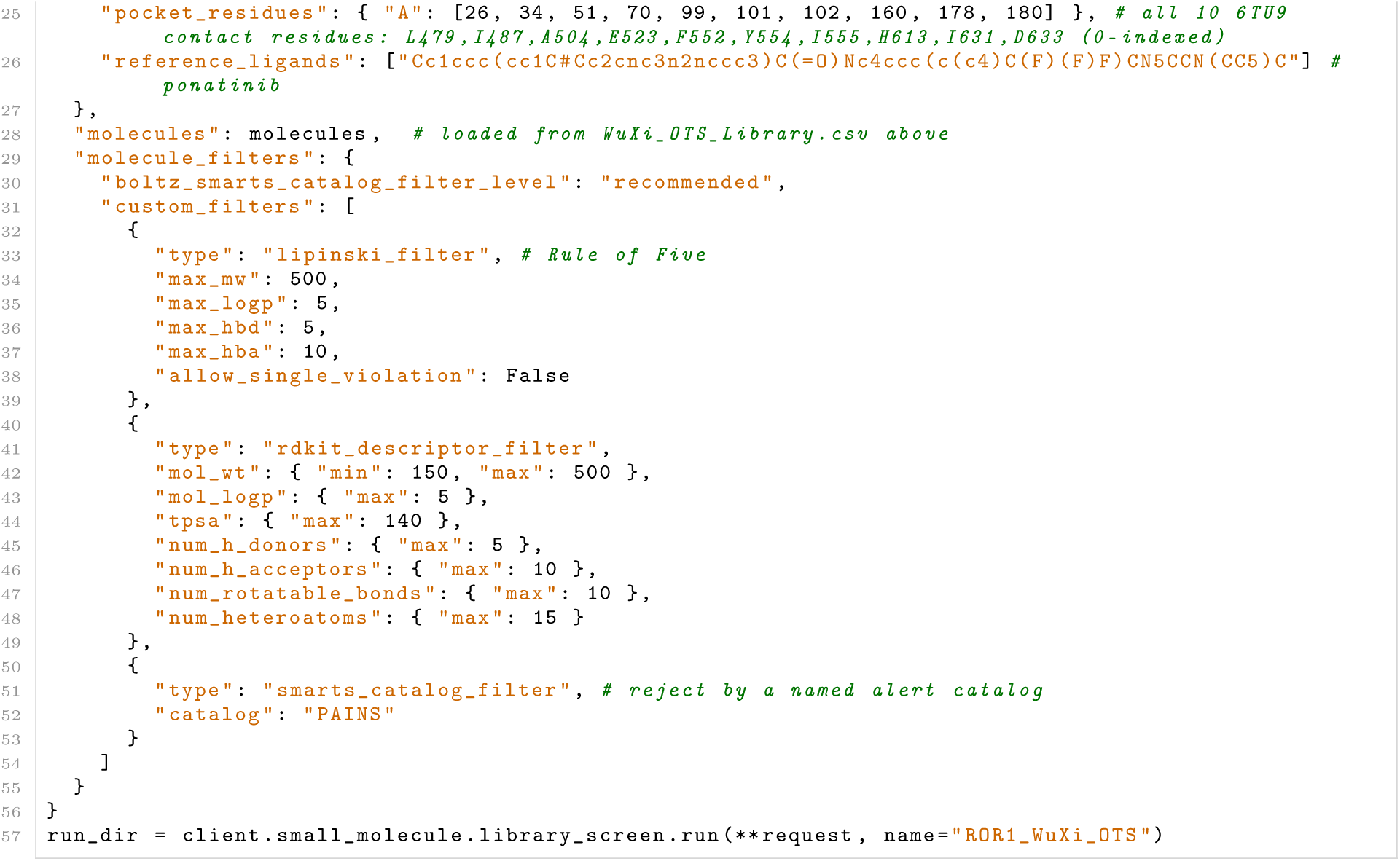

### F.3 Target specific inputs for successful non-collaboration targets

#### MALT1

Listing 3: Boltz Input target Parameters for MALT1

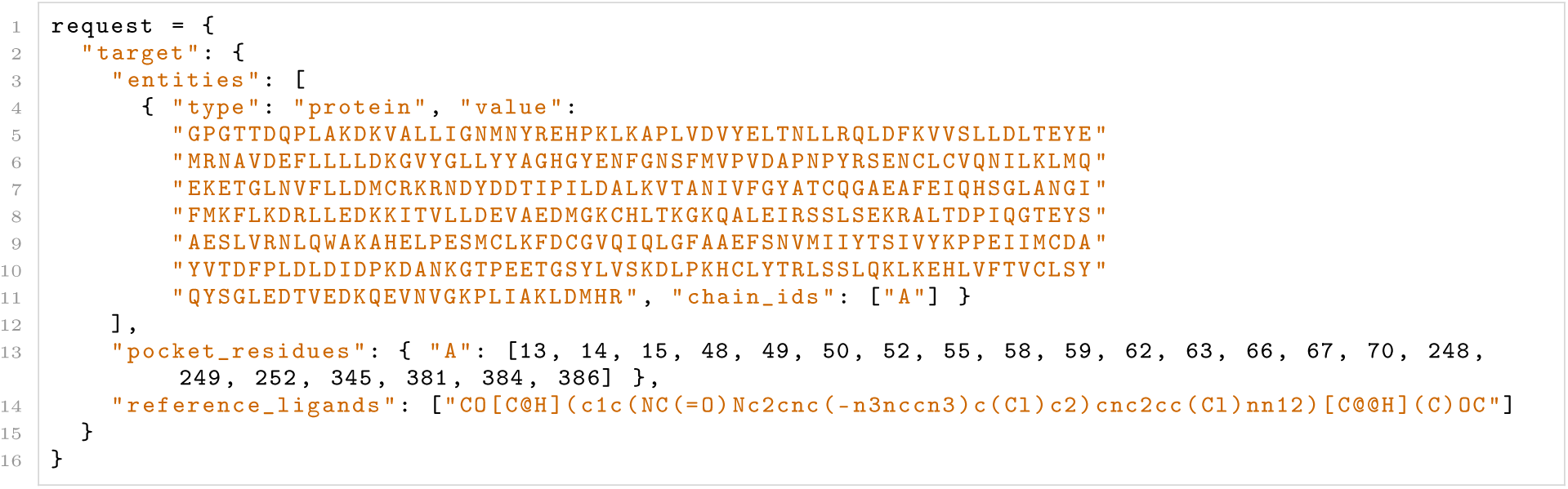

#### STAT6

Listing 4: Boltz Input target Parameters for STAT6

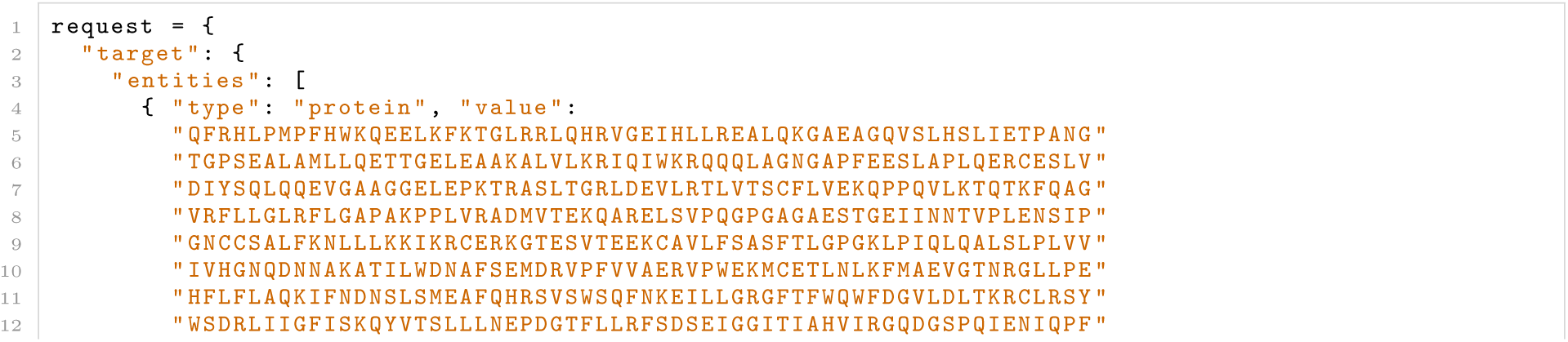
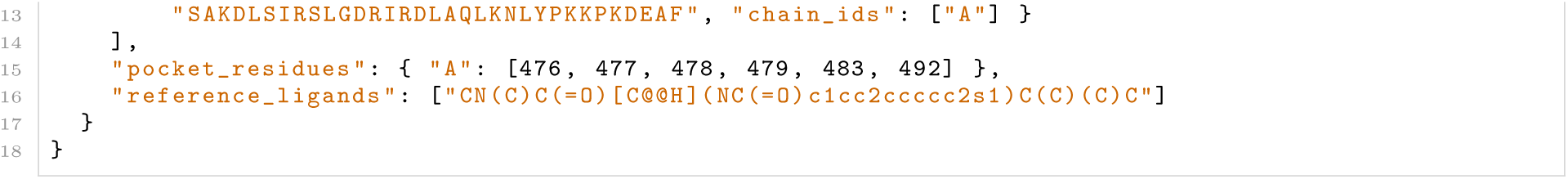

#### MRGPRX2

Listing 5: Boltz Input target Parameters for MRGPRX2

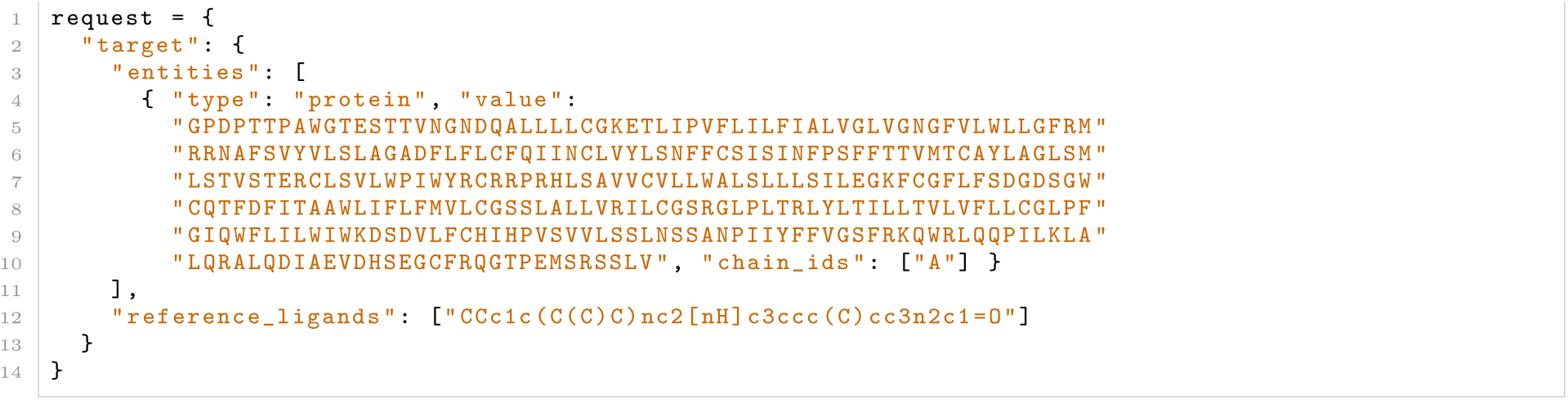

#### GLP-2

Listing 6: Boltz Input target Parameters for GLP-2

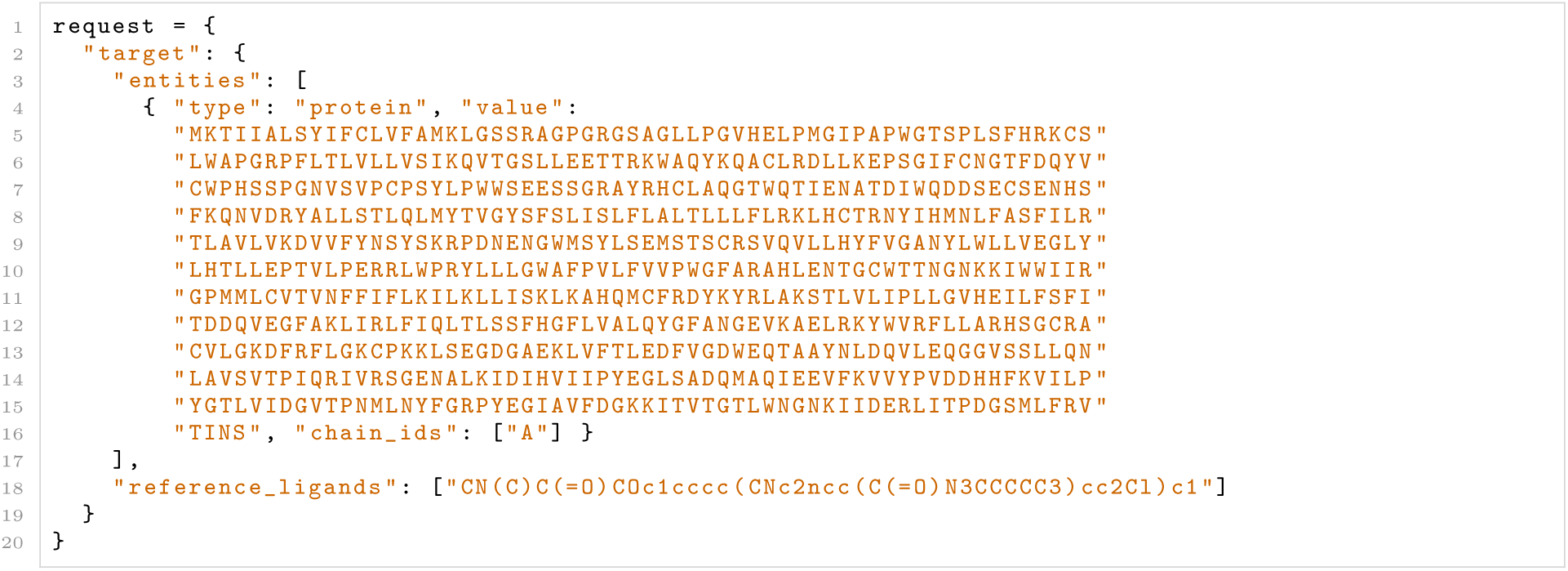

### F.4 Target specific inputs for unsuccessful non-collaboration targets

The input target parameters to replicate the screens using Boltz API for which no binders or actives from the WuXi OTS screening collection are described below:

#### Amylin 3 Receptor

Listing 7: Boltz Input target Parameters for the Amylin 3 Receptor (CTR + RAMP complex)

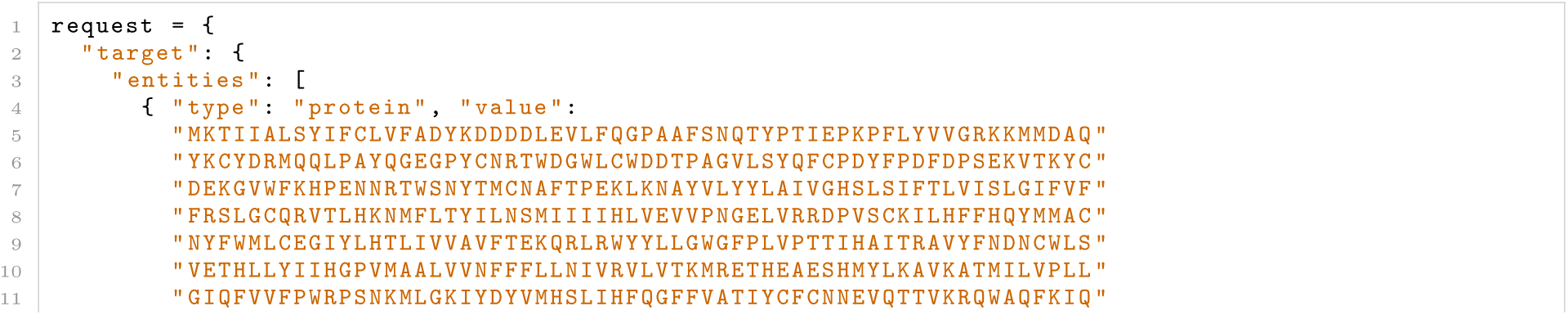
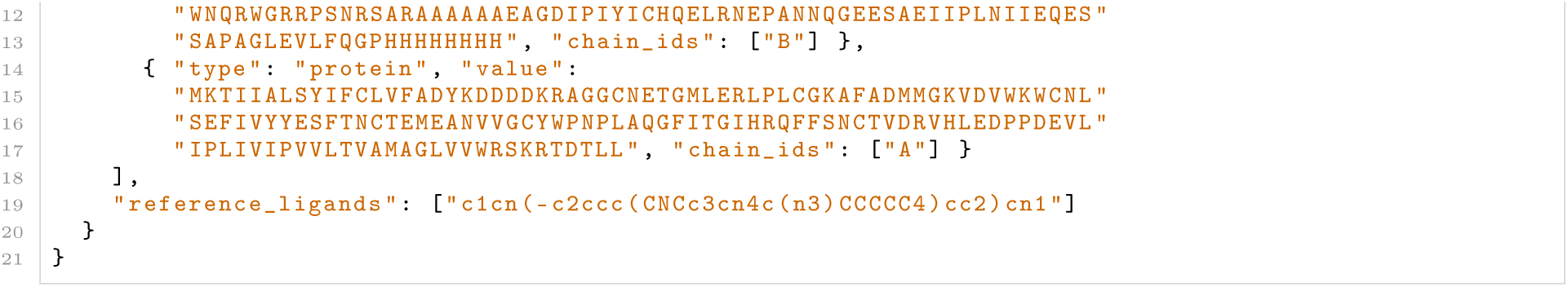

#### Nav1.8 VSD2

Listing 8: Boltz Input target Parameters for the Nav1.8 VSD2 domain

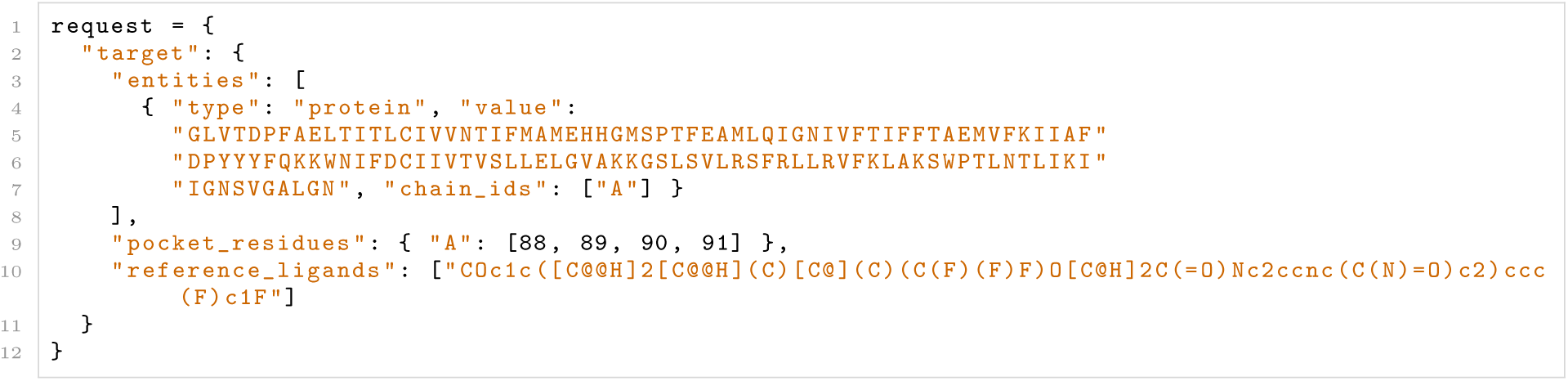

#### mGlu4 PAM

Listing 9: Boltz Input target Parameters for the mGlu4 PAM allosteric site

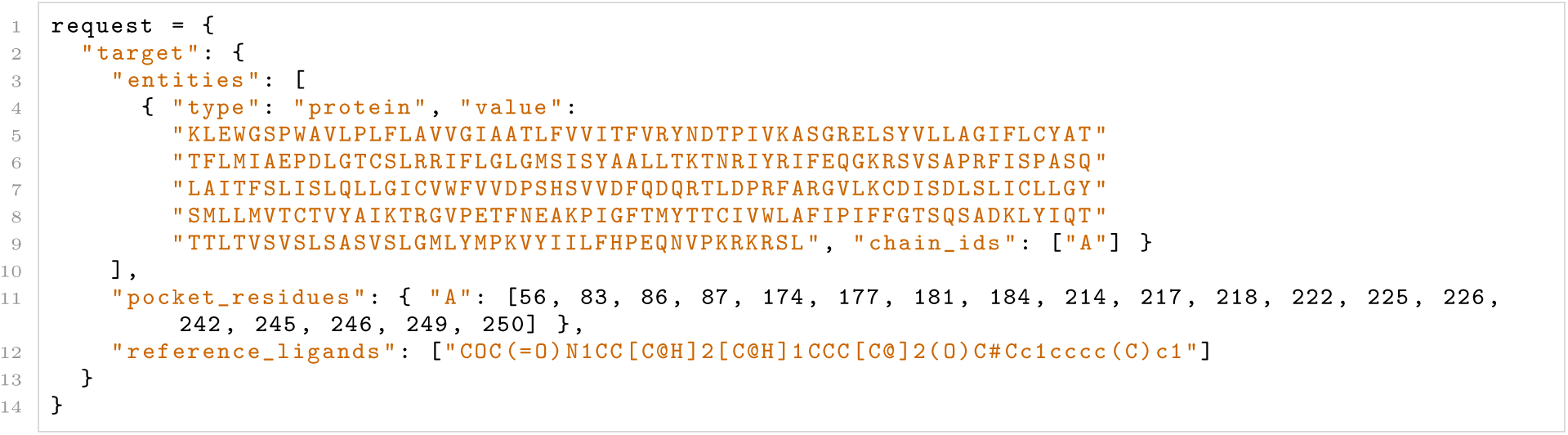

**Note**: For GPCR orthosteric receptors, no pocket residues were specified. This was to allow for either agonist or antagonist binding sites to be identified. For mGlu4 putative 7TM PAM site, pocket residues should be defined to exclude the extracellular orthosteric site.

### F.5 Boltz API ADME Endpoints

ADME predictions are included during small molecule screening using BoltzMol-1.

For submission of molecules to obtain ADME endpoints only, the following example call can be used:

Listing 10: Predicting small-molecule ADME with the Boltz API.

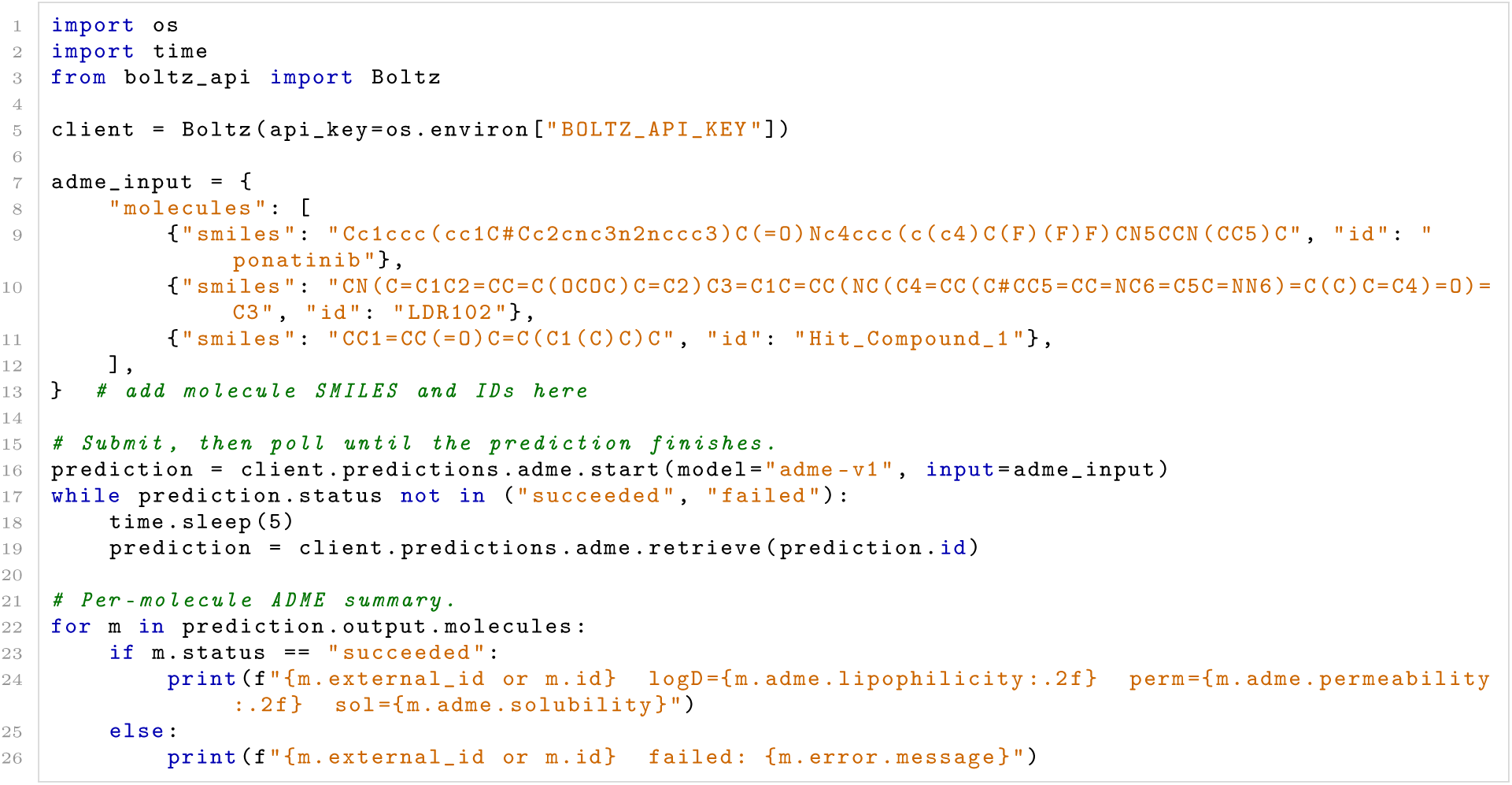

